# Integrated analyses of single-cell atlases reveal age, gender, and smoking status associations with cell type-specific expression of mediators of SARS-CoV-2 viral entry and highlights inflammatory programs in putative target cells

**DOI:** 10.1101/2020.04.19.049254

**Authors:** Christoph Muus, Malte D. Luecken, Gokcen Eraslan, Avinash Waghray, Graham Heimberg, Lisa Sikkema, Yoshihiko Kobayashi, Eeshit Dhaval Vaishnav, Ayshwarya Subramanian, Christopher Smilie, Karthik Jagadeesh, Elizabeth Thu Duong, Evgenij Fiskin, Elena Torlai Triglia, Meshal Ansari, Peiwen Cai, Brian Lin, Justin Buchanan, Sijia Chen, Jian Shu, Adam L Haber, Hattie Chung, Daniel T Montoro, Taylor Adams, Hananeh Aliee, J. Samuel, Allon Zaneta Andrusivova, Ilias Angelidis, Orr Ashenberg, Kevin Bassler, Christophe Bécavin, Inbal Benhar, Joseph Bergenstråhle, Ludvig Bergenstråhle, Liam Bolt, Emelie Braun, Linh T Bui, Mark Chaffin, Evgeny Chichelnitskiy, Joshua Chiou, Thomas M Conlon, Michael S Cuoco, Marie Deprez, David S Fischer, Astrid Gillich, Joshua Gould, Minzhe Guo, Austin J Gutierrez, Arun C Habermann, Tyler Harvey, Peng He, Xiaomeng Hou, Lijuan Hu, Alok Jaiswal, Peiyong Jiang, Theodoros Kapellos, Christin S Kuo, Ludvig Larsson, Michael A. Leney-Greene, Kyungtae Lim, Monika Litviňuková, Ji Lu, Leif S Ludwig, Wendy Luo, Henrike Maatz, Elo Madissoon, Lira Mamanova, Kasidet Manakongtreecheep, Charles-Hugo Marquette, Ian Mbano, Alexi Marie McAdams, Ross J Metzger, Ahmad N Nabhan, Sarah K. Nyquist, Lolita Penland, Olivier B Poirion, Sergio Poli, CanCan Qi, Rachel Queen, Daniel Reichart, Ivan Rosas, Jonas Schupp, Rahul Sinha, Rene V Sit, Kamil Slowikowski, Michal Slyper, Neal Smith, Alex Sountoulidis, Maximilian Strunz, Dawei Sun, Carlos Talavera-López, Peng Tan, Jessica Tantivit, Kyle J Travaglini, Nathan R. Tucker, Katherine Vernon, Marc H. Wadsworth, Julia Waldman, Xiuting Wang, Wenjun Yan, William Zhao, Carly G. K. Ziegler, The NHLBI LungMAP Consortium, The Human Cell Atlas Lung Biological Network

**Author notes:** These authors contributed equally (Christoph Muus, Malte D. Luecken, Gokcen Eraslan, Avinash Waghray, Graham Heimberg, Lisa Sikkema, Yoshihiko Kobayashi, Eeshit Dhaval Vaishnav, Ayshwarya Subramanian, Christopher Smilie, Karthik Jagadeesh, Elizabeth Thu Duong, Evgenij Fiskin, Elena Torlai Triglia, Meshal Ansari, Peiwen Cai, Brian Lin, Justin Buchanan, Sijia Chen, Jian Shu, Adam L Haber, Hattie Chung, Daniel T Montoro).

## Abstract

The COVID-19 pandemic, caused by the novel coronavirus SARS-CoV-2, creates an urgent need for identifying molecular mechanisms that mediate viral entry, propagation, and tissue pathology. Cell membrane bound angiotensin-converting enzyme 2 (ACE2) and associated proteases, transmembrane protease serine 2 (TMPRSS2) and Cathepsin L (CTSL), were previously identified as mediators of SARS-CoV2 cellular entry. Here, we assess the cell type-specific RNA expression of *ACE2*, *TMPRSS2*, and *CTSL* through an integrated analysis of 107 single-cell and single-nucleus RNA-Seq studies, including 22 lung and airways datasets (16 unpublished), and 85 datasets from other diverse organs. Joint expression of *ACE2* and the accessory proteases identifies specific subsets of respiratory epithelial cells as putative targets of viral infection in the nasal passages, airways, and alveoli. Cells that co-express ACE2 and proteases are also identified in cells from other organs, some of which have been associated with COVID-19 transmission or pathology, including gut enterocytes, corneal epithelial cells, cardiomyocytes, heart pericytes, olfactory sustentacular cells, and renal epithelial cells. Performing the first meta-analyses of scRNA-seq studies, we analyzed 1,176,683 cells from 282 nasal, airway, and lung parenchyma samples from 164 donors spanning fetal, childhood, adult, and elderly age groups, associate increased levels of *ACE2*, *TMPRSS2*, and *CTSL* in specific cell types with increasing age, male gender, and smoking, all of which are epidemiologically linked to COVID-19 susceptibility and outcomes. Notably, there was a particularly low expression of ACE2 in the few young pediatric samples in the analysis. Further analysis reveals a gene expression program shared by *ACE2^+^TMPRSS2^+^* cells in nasal, lung and gut tissues, including genes that may mediate viral entry, subtend key immune functions, and mediate epithelial-macrophage cross-talk. Amongst these are IL6, its receptor and co-receptor, *IL1R*, TNF response pathways, and complement genes. Cell type specificity in the lung and airways and smoking effects were conserved in mice. Our analyses suggest that differences in the cell type-specific expression of mediators of SARS-CoV-2 viral entry may be responsible for aspects of COVID-19 epidemiology and clinical course, and point to putative molecular pathways involved in disease susceptibility and pathogenesis.

## INTRODUCTION

COVID-19 is a global health threat due to its rapid spread, morbidity, and mortality. Despite progress in viral identification, sequencing of the full viral genome, creation of initial diagnostics, and the development of therapeutic hypotheses, many outstanding hurdles remain. These include deciphering the basis of the increased risk associated with certain demographic groups and identifying molecular mechanisms of disease pathogenesis.

The clinical presentation and transmission of COVID-19 is complex. Common symptoms include fever, cough, shortness of breath, chest pain, malaise, fatigue, headache, myalgias, anosmia, and diarrhea, while laboratory and radiographic findings include lymphopenia and ground-glass opacities on chest imaging, respectively^1–8^. Of an initial cohort of 1,099 hospitalized patients diagnosed with COVID-19, many developed diffuse alveolar damage (DAD) ^1^, pneumonia (79.1%), not infrequently complicated by acute respiratory distress syndrome (ARDS, 3.4%), and shock (1.0%), with 5.0% of patients requiring ICU admission and 2.2% requiring ventilation^4^. As the number of patients has surged, multi-system pathologies have been increasingly described, including kidney injury^9^, liver injury, gastrointestinal symptoms^4^, cardiac injury and dysfunction^2, 5, 10^, and multiorgan failure^1–4^. In addition to nasal and throat secretions, SARS-CoV-2 RNA has also been detected in saliva and stool specimens^10, 11^, suggesting possible alternative routes of transmission beyond respiratory droplets^12, 13^. SARS-CoV-2 may also infect the testis, similarly to SARS-CoV^14, 15^. Vertical transmission from mother to fetus remains a possibility. At least five neonates born to pregnant women with COVID-19 pneumonia were reported to test positive for SARS-CoV-2 infection after birth^16–18^ and other studies report newborns with elevated virus-specific antibodies to SARS-CoV-2 born to mothers with COVID-19^16, 19^. However, several other studies have thus far failed to find evidence for intrauterine transmission from pregnant women with COVID-19 to their newborns in cohorts as large as 38 patients^20–22^. Additionally, newborns from 9 COVID-19 patients who had cesarean deliveries in their third trimester tested negative for SARS-CoV-2^17^.

There is substantial variation in the clinical consequences of infection across individuals, ranging from asymptomatic carrier status to death. It has been suggested that undocumented subclinical infection contributes to the rapid dissemination of the virus^23^. As of April 14, 2020, COVID-19 has caused 1,970,225 confirmed infections and 124,544 deaths worldwide (https://coronavirus.jhu.edu/map.html). While true case fatality rate (CFR) is difficult to assess early in an epidemic^24–26^, estimates from modeling studies range from 0.9% - 3.3%^27, 28^. Disease severity and mortality rates show a striking rise with age^29^, with CFR estimates ranging from <0.1% for patients under 30 years old to >10% for those over 70^24^, with a slightly higher incidence and mortality in men^4, 30^. Children are significantly less likely than adults to develop severe disease, and reported pediatric deaths are rare^31^. Smoking is most likely associated with more severe disease^4, 32^. Finally, adults with pre-existing cardiovascular disease and acute myocardial injury have higher rates of disease acuity and death^33, 34^.

The coronavirus non-segmented, positive sense RNA genome of ∼30 kb contains coding regions for the expression of structural proteins including spike (S), envelope (E), membrane (M), and nucleocapsid (N) proteins. Virion recognition of host cells is initiated by interactions between the S protein and its receptor^35^. ACE2^36–38^, an essential regulator of the renin-angiotensin system^39^, is the receptor for both SARS-CoV^40^ and SARS-CoV-2. The receptor-binding domain of the SARS-CoV-2 S-protein has a higher binding affinity for human ACE2 than SARS-CoV^36, 41^, whereas the interaction with CD147 (encoded by the gene BSG), another reported receptor for the SARS-CoV-2 S-protein, is weak (CD147: Kd, 0.185 µM vs hACE2: Kd, ∼15 nM)^42^. Following receptor binding, the virus gains access to the host cell cytosol through acid-dependent proteolytic cleavage of the S protein. For SARS-CoV, a number of proteases including TMPRSS2 and CTSL cleave at the S1 and S2 boundary and S2 domain (S2’)^43^ to mediate membrane fusion and virus infectivity. For SARS-CoV-2, both pharmacological inhibition of endogenous TMPRSS2 protein and *TMPRSS2* overexpression support a role for TMPRSS2-mediated cellular entry^37, 44^.

The identification of the specific cell types that can be infected by SARS-CoV-2 will inform our understanding of disease transmission and pathogenesis, which are often cell context-specific. Studies suggest that key infection routes involve the nasal passages, airways, and alveoli, where epithelial cells play a key barrier role. Identifying putative target cells in other organs could inform our understanding of extra-pulmonary COVID-19 associated organ failure or of potential placental transmission. Early analyses of the Human Lung Cell Atlas revealed that some of the cells of the nasal passages, airways, alveoli, and gut co-express *ACE2* and *TMPRSS2*^45, 46^.

Here, we perform integrated analysis of 107 single-cell and single-nucleus RNA-Seq studies, including 22 studies of the lung and airways, and 85 additional studies of other diverse tissues, spanning both published and unpublished datasets. We comprehensively define the expression patterns of the *ACE2* viral receptor and accessory proteases genes. We test how their expression is related to age (from prenatal to old age), sex, and smoking status. We identify gene expression programs associated with cells that can be infected by the virus and compare these programs across specific cell types, organs, and species. To further inform future studies, we assess the conservation of these human features in mouse models and explore the expression of other proteases that may play a role in the viral replication cycle.

## RESULTS

### A cross-tissue survey identifies dual-positive *ACE2^+^TMPRSS2^+^* cells in nasal, airway, alveolar, and intestinal epithelia, as well as in other organs associated with COVID-19 pathology or transmission

Previous analyses of Human Cell Atlas datasets established that ACE2, the viral receptor, and one of its entry-associated proteases, TMPRSS2, are expressed in nasal, lung, and gut epithelial cells^45^. Specifically, nasal goblet cells and multiciliated cells comprised the highest fraction of dual-positive *ACE2^+^TMPRSS2^+^* cells^45^, consistent with a plausible role for a nasal viral reservoir that supports transmissivity. In the distal lung, co-expression occurred in AT2 cells^45, 47, 48^. Previous surveys across other tissues also showed a relatively high portion of *ACE2^+^TMPRSS2^+^* cells within colonic enterocytes, another potential viral reservoir that promotes viral transmission^49^.

To perform a comprehensive survey, we enumerated the proportion of dual-positive *ACE2^+^TMPRSS2^+^* cells and *ACE2^+^CTSL^+^* cells^50^ across 92 human studies (including seven of the lung and airways) with single-cell or single-nucleus RNA-seq (sc/snRNA-seq) (**Fig. 1, Methods, Supplementary Table 1 and 2**). These included a large survey of published datasets from diverse tissues, which we assigned to five broad cell categories, (**Fig. 1a,b, Extended Data Fig. 1, 2, Supplementary Table 1**). We further analyzed more finely annotated published and unpublished datasets (**Methods, Fig. 1c,d, Supplementary Table 3**). Consistent with previous reports^45^, dual-positive *ACE2^+^TMPRSS2^+^* cells in the proximal airways were largely secretory goblet and multiciliated cells, and dual-positive cells in the distal lung were largely AT2 cells (**Fig. 1c, Extended Data Fig. 3a**).

**Figure 1.**
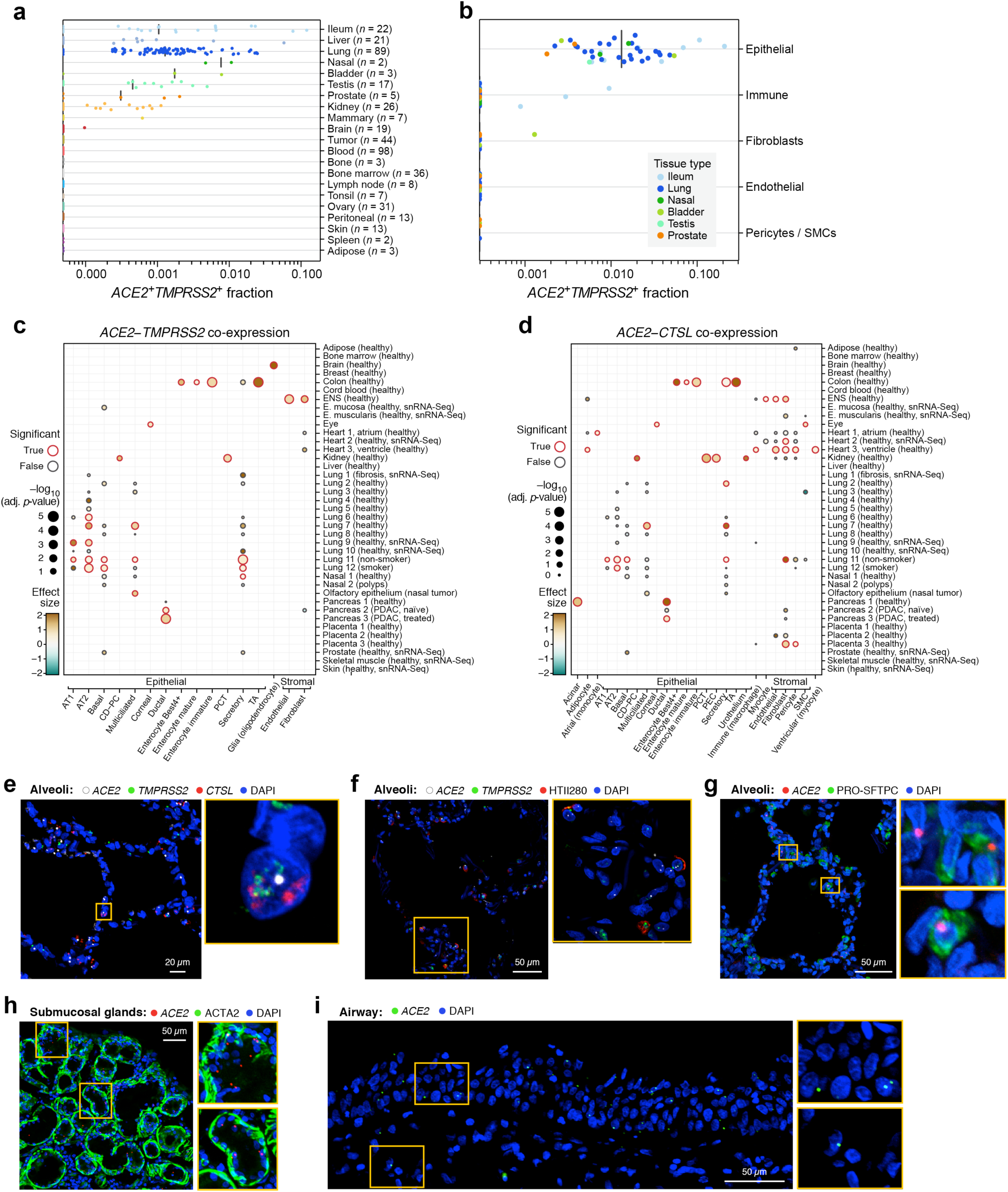
A cross-tissue survey of *ACE2^+^TMPRSS2^+^* cells shows enrichment in cells at reported sites of disease transmission or pathogenesis. (a,b) Dual positive cells are more prevalent in epithelial organs and cells. (a) Proportion of *ACE2^+^TMPRSS2^+^* cells (y axis) per dataset (dots) from 21 tissues and organs (rows). (b) Proportion of *ACE2^+^TMPRSS2^+^* cells (y axis) within cell clusters (dots) annotated by broad cell-type categories (rows) within each of the top 7 enriched datasets (color legend, inset). (c,d) Significant co-expression of *ACE2^+^TMPRSS2^+^* or *ACE2^+^CTSL^+^* highlights cells from tissues implicated in transmission or pathogenesis. Significance of co-expression (dot size - log10(adjusted P-value), Methods; red border: FDR<0.1) of *ACE2^+^TMPRSS2^+^* (c) or *ACE2^+^CTSL^+^* (d) and effect size (dot color, color bar) for finely annotated cell classes (columns) from diverse tissues (rows). Only tissues and cells in at least one significant co-expression relationship are shown (Methods). (e-i) *In situ* validation of dual positive cells in the lung, airways, and submucosal gland. (e-g) PLISH and immunostaining in human adult lung alveoli for (e) *ACE2* (white), *TMPRSS2* (green) and *CTSL* (red), (f) *ACE2* (white), *TMPRSS2* (green) and HTII-280 (red), (g) *ACE2* (red) and Pro-SFTPC (green). (h) PLISH and immunostaining in human large airway submucosal glands. *ACE2* (red), ACTA2 (green) and DAPI (blue). (i) PLISH in human adult large airway. *ACE2* (green) and DAPI (blue).

**Figure 2.**
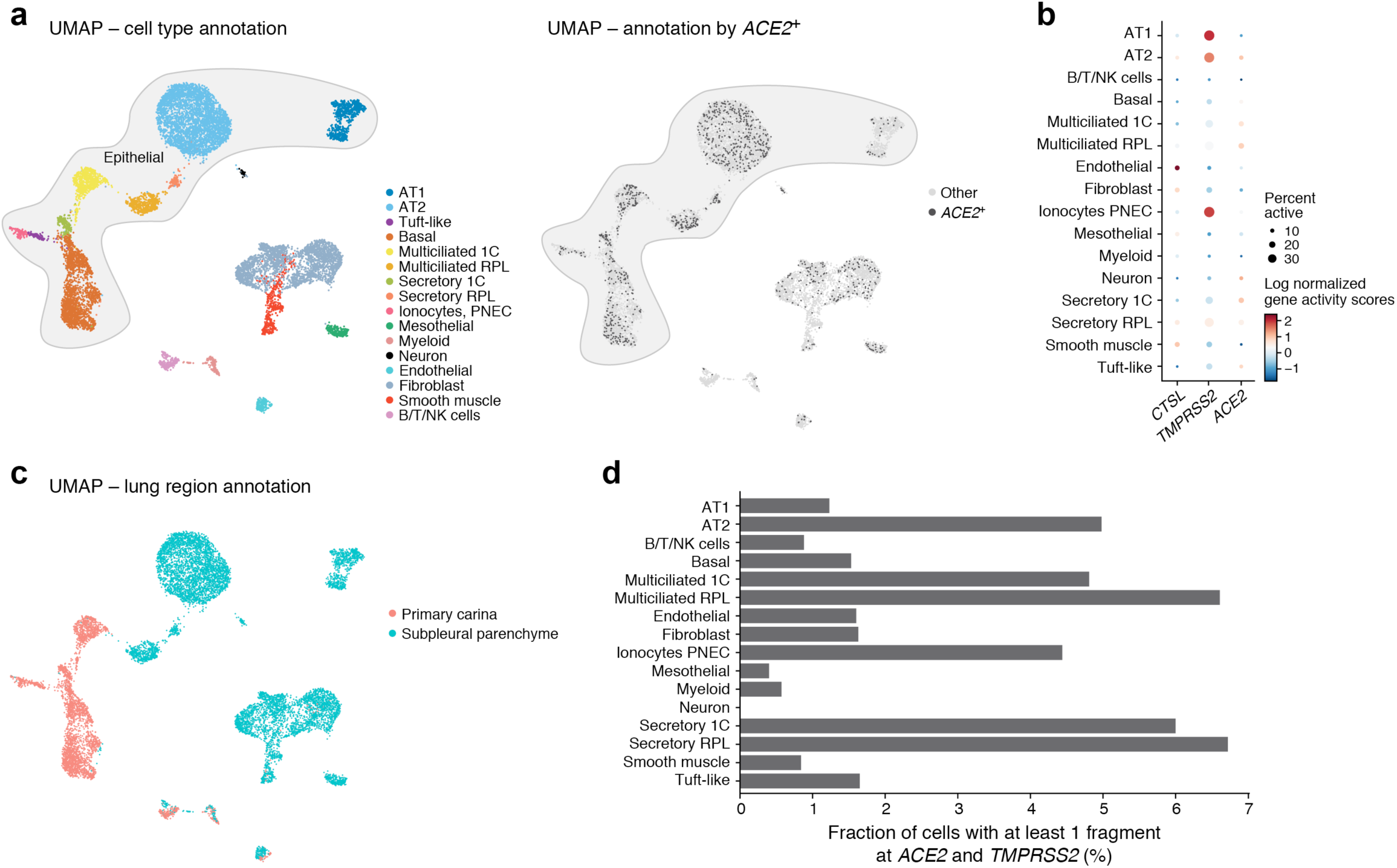
Open chromatin at the *ACE2* and *TMPRSS2* loci in AT2, ciliated and secretory cells in the lung and airways (a,c) Single-cell ATAC-seq of lung samples from primary carina (1C) and subpleural parenchyme (RPL) (*n*=1 patient, *k*=3 samples, 3,366 cells from 1C, 8,340 cells from RPL). Uniform Manifold Approximation and Projection (UMAP) embedding of scATAC-seq profiles (dots) colored by (a, left) cell types, (a, right) cells with at least 1 fragment (indicating accessibility, open chromatin) mapping to the *ACE2* gene locus (defined as −2kb upstream the Transcription Start Site to Transcription End Site), or by sample location (c). Grey shaded area: epithelial cell types. (b) Inferred gene activity of *ACE2, TMPRSS2, CTSL* across cell types. Log normalized mean “scATAC activity score” (quantified from accessibility, open chromatin) (dot color) and proportion of cells with active scores (dot size) for *ACE2, TMPRSS2,* and *CTSL* (columns) across different cell types (rows) from the primary carina (1C) and subpleural parenchyme (RPL). (d) Some AT2, ciliated and secretory cells have accessible chromatin at both *ACE2* and *TMPRSS2* loci. Proportion of cells (*x* axis) in each cell type (*y* axis) with accessible chromatin (at least 1 fragment) at both the *ACE2* and *TMPRSS2* loci (defined as −2kb upstream of the Transcription Start Site to Transcription End Site).

**Figure 3.**
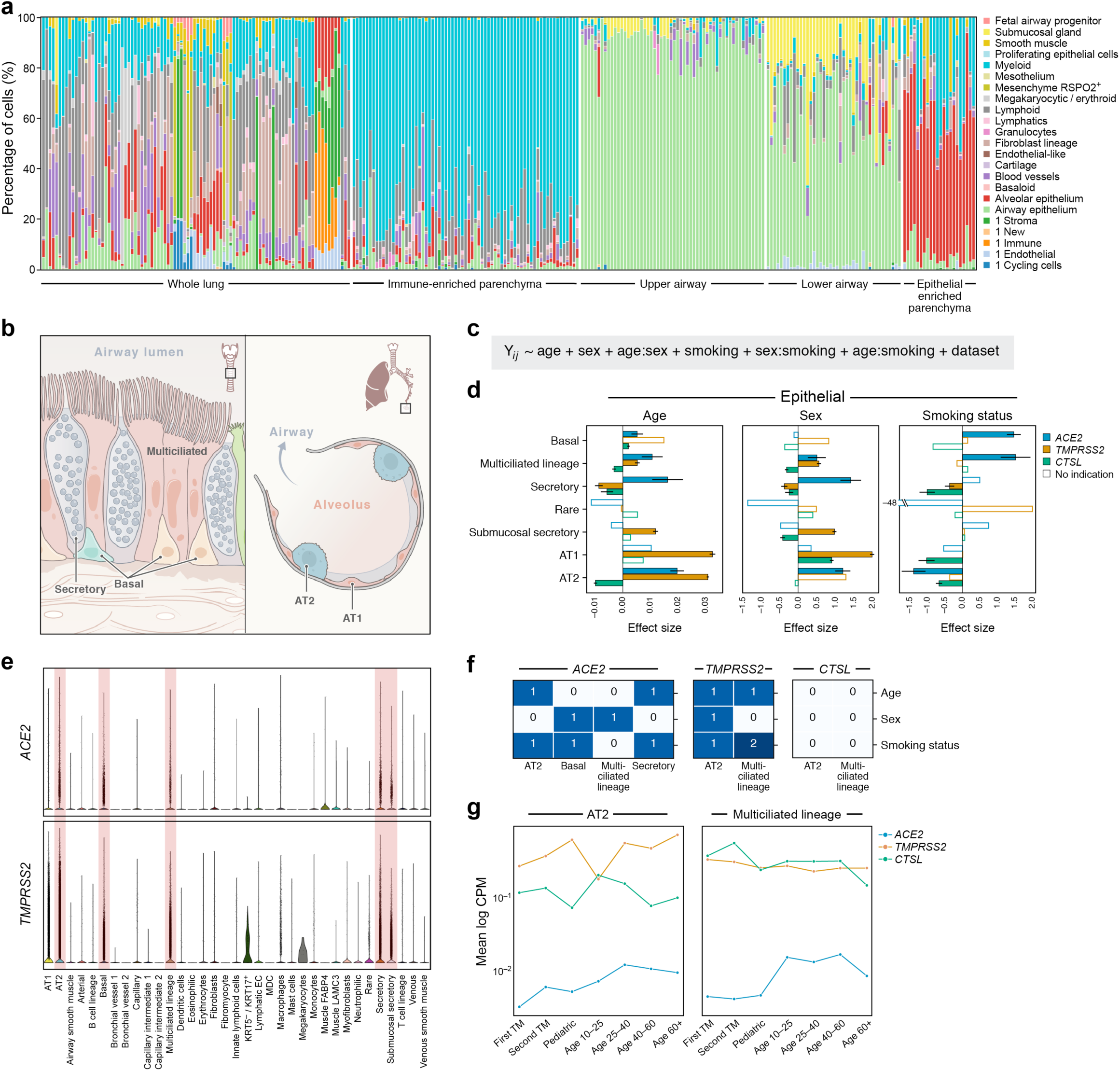
*ACE2*, *TMPRSS2,* and *CTSL* expression in AT2 cells increases with age and smoking status, and in men (a) Samples in the aggregated lung and airway dataset partition to several classes by their cell composition. Percentage of cells (y axis) by level 2 cell annotations (Annotations with a preceding “1” indicate coarse annotations) across samples (x axis). The 282 samples are ordered by sample composition clusters. (b) Schematic of key lung and airway cell types highlighted in the study. (c) Statistical model. Model fitted to the data to assess sex, age, and smoking status associations with expression of the three genes. *Y_i,j_* denotes gene counts and *nUMI* denotes the total UMI counts per cell. (d) Age, sex, and smoking status associations with expression of *ACE2* (blue), *TMPRSS2* (orange), and *CTSL* (green) in epithelial cells. Effect size (x axis) of the association, in log fold change (sex, smoking status) or slope of log expression with age. Colored bars: associations with an FDR-corrected p-value<0.05, where pseudo-bulk analysis shows a consistent effect direction. Error bars: standard errors around coefficient estimates. (e) Distribution of *ACE2* and *TMPRSS2* expression across level 3 lung cell types. Red shading indicates the main cell types that express *ACE2* and *TMPRSS2*. (f) Hold out analysis shows the robustness of associations to holding out a dataset. The values show the number of held-out datasets that result in loss of association between a given covariate (rows) and *ACE2, TMPRSS2, or CTSL* expression in a given cell type (columns). Robust trends are determined by significant effects that are robust to holding out any dataset (0 values). (g) Low expression in pediatric samples. Mean expression level (log CPM, y axis) of *ACE2* (blue), *TMPRSS2* (orange), and *CTSL* (green) across age bins (x axis) in AT2 (left) and ciliated (right) cells. Pediatric samples: 0-10 years. Samples from past or current smokers were removed from this plot to avoid smoking confounders. Error bars are omitted due to y-axis limitations. They are typically 10-fold the mean value (Supplementary Table 5). Multiciliated and AT2 cells are shown as these cell types are present in fetal data, and show significant age associations with *ACE2* expression.

*ACE2* expression in secretory (especially goblet) and AT2 cells is also supported by scATAC-seq from the primary carina and subpleural parenchyma, respectively (**Fig. 2**, *n*=3 samples per location, *n*=1 patient), showing accessibility at the *ACE2* locus in a portion of AT2 cells (10.8%, 364 out of 3,371 cells, **Methods**), as well as secretory and multiciliated cells, and to a lesser extent some basal and tuft cells (**Fig. 2a-c**). The proportion of AT2 cells with an open *ACE2* locus is somewhat higher than of *ACE2^+^* AT2 cells by scRNA-seq in the same patient and region (pleura: 10.8%, 364 out of 3,371 cells *vs*. 6.2%, 199 out of 3,215 cells). Cells with accessible chromatin at both the *ACE2* and *TMPRSS2* loci were also most commonly found in epithelial cells, especially AT2 cells (**Fig. 2d**, 4.98%, 168 of 3,371 AT2 cells from a subpleural sample; comparable to 3.9%, 124 of 3,215 in matched scRNA-Seq), secretory cells (6.72%, 8 of 119 secretory cells from small airway of the subpleural region, and 6%, 12 of 200 secretory cells from primary carina of the large airway), and multiciliated cells (6.61%, 32 of 484 multiciliated cells from subpleural sample, and 4.81%, 31 out of 644 multiciliated cells from primary carina).

**Figure 4:**
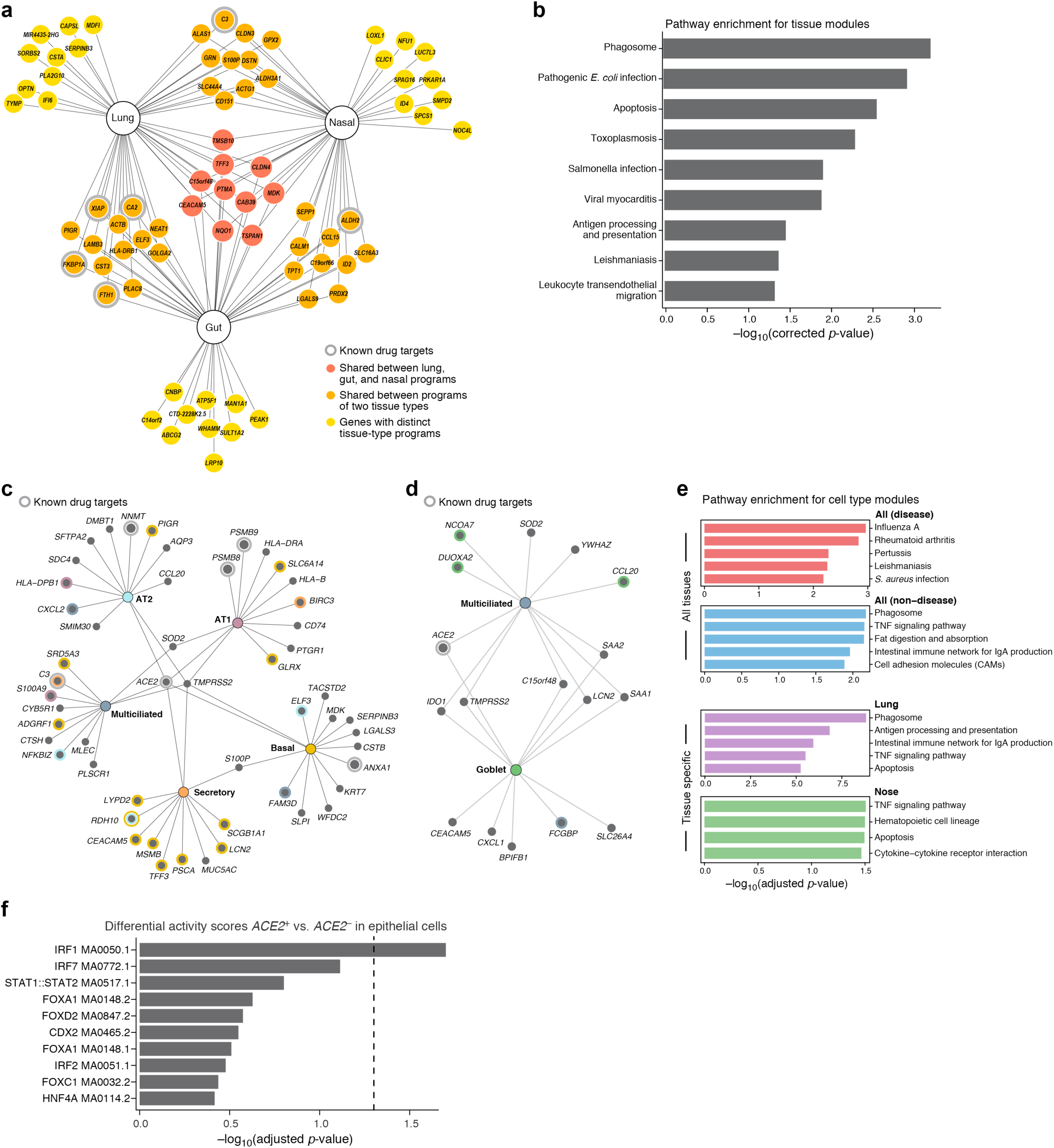
Tissue and cell-type-specific gene modules in *ACE2^+^TMPRSS2^+^* cells highlight immune and inflammatory features (**a,b**) Tissue programs of *ACE2^+^TMPRSS2^+^* cells in lung, gut, and nasal samples. (**a**) Selected tissue program genes. Node: gene; Edge: program membership. Genes are selected heuristically for visualization (Methods). (**b**) Enrichment (-log10(adj P-value), x axis) of KEGG pathway gene sets (y axis) in the full tissue programs. (**c-e**) Cell programs of *ACE2^+^TMPRSS2^+^* cells. (c,d) Top 12 genes from each cell program recovered for different lung (**c**) or (**d**) nasal epithelial cell-type (nodes, colors). Colored concentric circles: overlap with a gene in the top 250 significant genes in other cell types. ACE2 and TMPRSS2 are included even if not among the top 12. (**e**) Enrichment (-log10(adj P-value), x axis) of KEGG disease and non-disease pathway gene sets in either highly significant genes across all tissues (top) or in specific tissues (lung, nose, bottom). (**f**) Motif activity in immune TFs in *ACE2*^+^ cells. Significance (-log10(adjusted p-value), x axis) of the top 10 differential “motif activity scores” (Methods) between epithelial *ACE2^+^* cells or *ACE2^-^* cells (y axis). (Epithelial cells are: AT1, AT2, secretory, ciliated, ionocytes, and neuroendocrine cells, highlighted in the gray shaded area in Fig. 2c). (n=2 locations: primary carina and lung lobes, n=3 samples per location, n=1 patient). Motifs are extracted from the JASPAR2020 database, motif code is shown in each row. Dashed line: threshold for significance (adjusted p-value of 0.05).

There were dual-positive *ACE2^+^TMPRSS2^+^* cells in tissues beyond the respiratory system (**Fig. 1a-c**), including enterocytes, pancreatic ductal cells, prostate luminal epithelial cells, cholangiocytes^51^, oligodendrocytes in the brain, inhibitory enteric neurons, heart fibroblasts/pericytes^52^, and fibroblasts and pericytes in multiple other tissues (**Fig. 1c**). *ACE2^+^TMPRSS2^+^* epithelial cells were most prevalent (in order) within the ileum, liver, lung, nasal mucosa, bladder, testis, prostate, and kidney (**Fig. 1a**). Enterocytes had a substantial proportion of dual-positive cells (**Fig. 1c**), and are possibly part of a renin-angiotensin multicellular circuit^53^. In line with the kidney’s role in the renin-angiotensin-aldosterone system, dual-positive cells are enriched in the proximal tubular cells and in principal cells of the collecting duct (**Fig. 1a,c**). Interestingly, brain oligodendrocytes, multiciliated and sustentacular cells in olfactory epithelium, AT1 cells in non-smoker lung, and ductal cells in pancreas – all *ACE2^+^TMPRSS2^+^* – were also all enriched for *MYRF*, a transcription factor necessary for myelination in the brain and sufficient to induce expression of the myelin proteins *MOG* (myelin oligodendrocyte glycoprotein) and *MBP* (myelin basic protein)^54^. *ACE2*^+^*CTSL*^+^ cells were enriched in additional subsets, associated with COVID-19 pathology, most notably the olfactory epithelium, ventricular cardiomyocytes, heart macrophages, and pericytes in multiple tissues, including the heart, lung, and kidney (**Fig. 1d**).

The presence of dual-positive cells in the lung, heart, and kidney may reflect that cells in these organs may be direct targets of viral infection and pathology^5, 55^. Dual-positive cells in the sustentacular and basal cells of the olfactory epithelium (**Fig. 1c**) may be associated with a loss of the sense of smell^56^. Dual-positive cells in the corneal and conjunctival epithelium, may contribute to viral transmission^4, 57^. Dual positive cardiomyocytes may be related to “direct” cardiomyocyte damage (see Tucker et al. companion manuscript^58^), whereas heart pericytes may indicate a vascular component to the cardiac dysfunction, and could contribute to increased troponin leak in patients without coronary artery disease. Notably, ACE2-expressing heart pericytes in another dataset (Tucker et al) is even higher (30%) than any other tissue dataset analyzed here (max. 21.7% in kidney, **Extended Data Fig. 4**). Despite the lymphopenia observed with COVID-19^2, 4, 59, 60^, we did not typically observe *ACE2* mRNA expression in scRNA-seq profiles in the bone marrow or cord blood (**Fig. 1a,b**), although there was ACE2 expression in some tissue macrophages, including alveolar and heart macrophages (**Extended Data Fig. 4**). Further studies of ACE2 RNA and protein expression in COVID-19 disease tissue will help elucidate its expression in immune cells^61^.

### Validation of ACE2, TMPRSS2 and CTSL mRNA and protein expression in airway and alveolar epithelium

To validate our findings from scRNA-seq analysis and to determine the spatial expression patterns of *ACE2*, *TMPRSS2*, and *CTSL*, and their corresponding proteins we performed fluorescence *in situ* hybridization and immunohistochemistry on tissue sections of airway and alveoli from healthy donor lungs that were rejected for lung transplantation.

First, we performed triple fluorescence *in situ* hybridization to identify *ACE2*, *CTSL* and *TMPRSS2* on alveolar sections. We observed co-expression, albeit at low levels, of all three genes in alveolar cells (**Fig. 1e**). We then performed co-staining with cell type-specific markers. We observed *ACE2* transcripts in a subset of type 2 (AT2) cells identified by canonical AT2 protein markers, HTII-280 and pro-SFTPC (**Fig. 1f,g**). Similarly, we observed *TMPRSS2* gene expression in HTII-280+ AT2 cells (**Fig. 1f**). Immunostaining for TMPRSS2 protein further confirmed AT2 cell expression (**Extended Data Fig. 5a**). We also observed TMPRSS2 protein expression at low levels in some AT1 cells identified by the canonical AT1 protein marker AGER (**Extended Data Fig. 5a**). Of note, some non-epithelial cells also expressed these three genes. We further validated the expression of *ACE2* by bulk mRNA-seq in sorted AT2 cells, including those from long-term cultured alveolar organoids (**Extended Data Fig. 5b**). We then performed immunohistochemistry and deployed three different available putative ACE2 antibodies to establish ACE2 protein expression (**Supplementary Table 4**). One of these 3 antibodies, the one used previously to functionally block cellular viral entry, specifically labeled adult pro-SFTPC-positive AT2 cells (**Extended Data Fig. 5c**). As a cautionary note, the lack of agreement between antibody staining patterns suggests that some of these antibodies may be non-specific.

**Figure 5:**
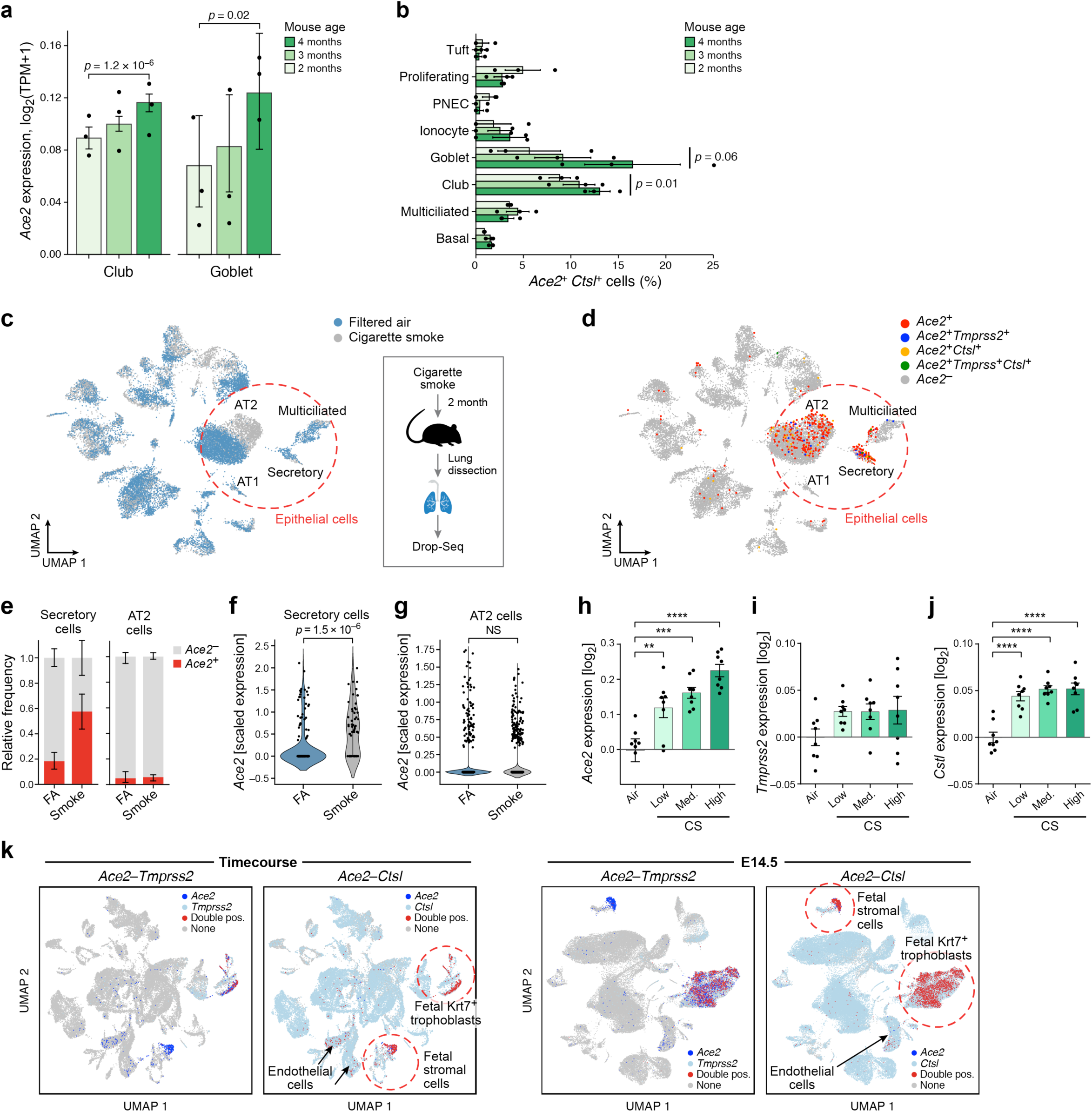
***Ace2, Tmprss2 and Ctsl2 expression in similar cell types and follows similar patterns with age and smoking.*** (**a**) Gradual increase in *Ace2* expression by airway epithelial cell type with age. Mean expression (y axis) of *Ace2* in different airway epithelial cells (*x* axis) of mice of three consecutive ages (color legend, upper right). Shown are replicate mice (dots), mean (bar), and error bars (standard error of the mean (SEM)). (**b**) Increase in proportion of *Ace2^+^Ctsl^+^* goblet and club cells with age. Percent of *Ace2^+^Ctsl^+^* cells (x axis) in different airway epithelial cell types (y axis) of mice of three consecutive ages (color legend, upper right). Shown are replicate mice (dots), mean (bar), and error bars (SEM). (**c-j**) Increase in *Ace2* expression in secretory cells with smoking. Mice were daily exposed to cigarette smoke or filtered air as control for two months after which cells from whole lung suspensions were analyzed by scRNA-seq (Drop-Seq). (**c,d**) UMAP of scRNA-seq profiles (dots) colored by experimental group (c) or by *Ace2*^+^ cells and indicated double positive cells (d). Alveolar epithelial cells (AT1 and AT2) and airway epithelial secretory and ciliated cells are marked. (**e**) The relative frequency of *Ace2^+^* cells is increased by smoking in airway secretory cells but not AT2 cells. Relative proportion (y axis) of *Ace2^+^* (red) and *Ace2^-^* (grey) cells in smoking and control mice of different cell types (x axis). (**f, g**) Expression of *Ace2* is increased in airway secretory cells, but not in AT2 cells. Distribution of *Ace2* expression (y axis) in secretory (f) and AT2 (g) cells from control and smoking mice (x axis). (**h-j**) Re-analysis of published bulk mRNA-Seq^101^ of lungs exposed to different daily doses of cigarette smoke show increased expression of (**h**) *Ace2*, (**i**) *Tmprss2*, and (**j**) *Ctsl* after five months of chronic exposure. (**k**) Expression in placenta. UMAP plots showing expression of *Ace2, Tmprss2* and *Ctsl* in single and double positive cells from embryonic days 9.5 to 18 (left) and embryonic day 14.5 (right) of mouse placenta development

Previous studies have revealed that ACE2 is highly enriched in mucous cells of the nasal and lower respiratory tract epithelium. In healthy lungs, both large and small airways contain mucous cells in surface airway epithelium, albeit at low numbers. However, submucosal glands (SMGs) that reside deep within the airway tissues are composed of abundant mucous cells. To test whether these mucous cells also express *ACE2, TMPRSS2, and CTSL*, we performed an integrated analysis on scRNA-seq datasets obtained from microdissected SMGs of the healthy donors. We observed overlapping expression and relatively high enrichment of *AEC2*, *TMPRSS2* and *CTSL* in mucous cells of the SMGS (**Extended Data Fig. 5e**). *In situ* transcript analysis for *ACE2* further confirmed the presence of transcripts in acinar epithelial cells of the SMGs (**Fig. 1h**), and cells expressing *ACE2* in the large airway epithelium (**Fig. 1i**).

### *ACE2* and *TMPRSS2* expression in airway epithelial and AT2 cells increases with age and is higher in men, and airway epithelial cell expression increases with smoking

We next sought to understand how the expression of each of these three key genes -- *ACE2*, *TMPRSS2*, and *CTSL* -- in specific cell subsets may relate to three key covariates that have been associated with disease severity: age (older individuals are more severely affected), sex (males are more severely affected), and smoking (smokers are more severely affected)^62^. We integrated samples across many studies, as no single dataset generated to date is sufficiently large to address this question. We assembled 22 datasets (**Supplementary Table 2, Supplementary Data File D3**), comprised of 1,176,683 cells from 164 individuals, spanning 282 healthy nasal, lung, and airway samples profiled by scRNA-seq or snRNA-seq from either biopsies, resections, entire lungs that could not be used for transplant, or post mortem examinations, allowing us to study a diversity of respiratory regions and cell types (**Fig. 3a**). These included 6 published datasets^63–68^ and 16 datasets that are not yet published^69–73^. In the case of unpublished data, we only obtained single-cell expression counts for the three genes, as well as the total UMI counts per cell, cell identity annotations, and the relevant anonymous clinical variables (age and sex, as well as smoking status when ascertained). Cell identity annotations were manually harmonized using an ontology with three levels of annotation specificity (**Fig. 3b, Supplementary Table 2**); focusing on levels 2 and 3 allowed us to include a large number of datasets, while retaining relatively high cell subset specificity (**Fig. 3a,b**). To facilitate rapid data sharing, we analyzed data pre-processed by each data-generating team at the level of gene counts, using total counts as a size factor. We used Poisson regression (diffxpy package; **Methods**) to model the association between the expression counts of the three genes and age, sex, and smoking status, and their possible pair-wise interactions (**Fig. 3c**), using total counts as an offset, and dataset as a technical covariate to capture sampling and processing differences. It should be noted that modeling interaction terms was crucial as their omission resulted in reversed effects for age and sex for particular cell types (**Discussion**). This model was fitted to non-fetal lung data (761,693 cells, 165 samples, 77 donors, 10 datasets) within each cell type to assess cell-type specific association of these covariates with the three genes. To further validate sex and age associations, we fit a simplified version of the model without smoking status covariates to the full non-fetal lung data (986,342 cells, 225 samples, 125 donors, 16 datasets).

Uncertainty is challenging to model in our single-cell meta-analysis as variability exists on the levels of both donors and cells. For simplicity, we modeled the overall variance with both contributions covered implicitly by treating each cell as an independent observation. As cells from the same donor cannot be typically regarded as independent observations, this can result in inflated p-values, especially when there are few donors for a particular cell type. To counteract this limitation, we employed three approaches: (**1**) We used a simple noise model (Poisson) to reduce the chance of overfitting donor variability to obtain spurious associations; (**2**) We confirmed significant associations from the single-cell model in a pseudo-bulk analysis to ensure effect directions are consistent when modeling only donor variation (**Methods, Fig. 3d, Extended Data Fig. 6, 7, 8, Supplementary Data D1**); and, (**3**) We investigated whether significant associations change direction when holding out any one dataset to ensure that the effect is not dominated by the inclusion of many cells from only one source (**Methods, Fig. 3f, Supplementary Data D1**). We regarded an association that passes all of these validations as a *robust trend*, while associations that appear dominated by a single dataset (often because this dataset is a major contributor of a given cell type) were denoted as *indications*.

**Figure 6.**
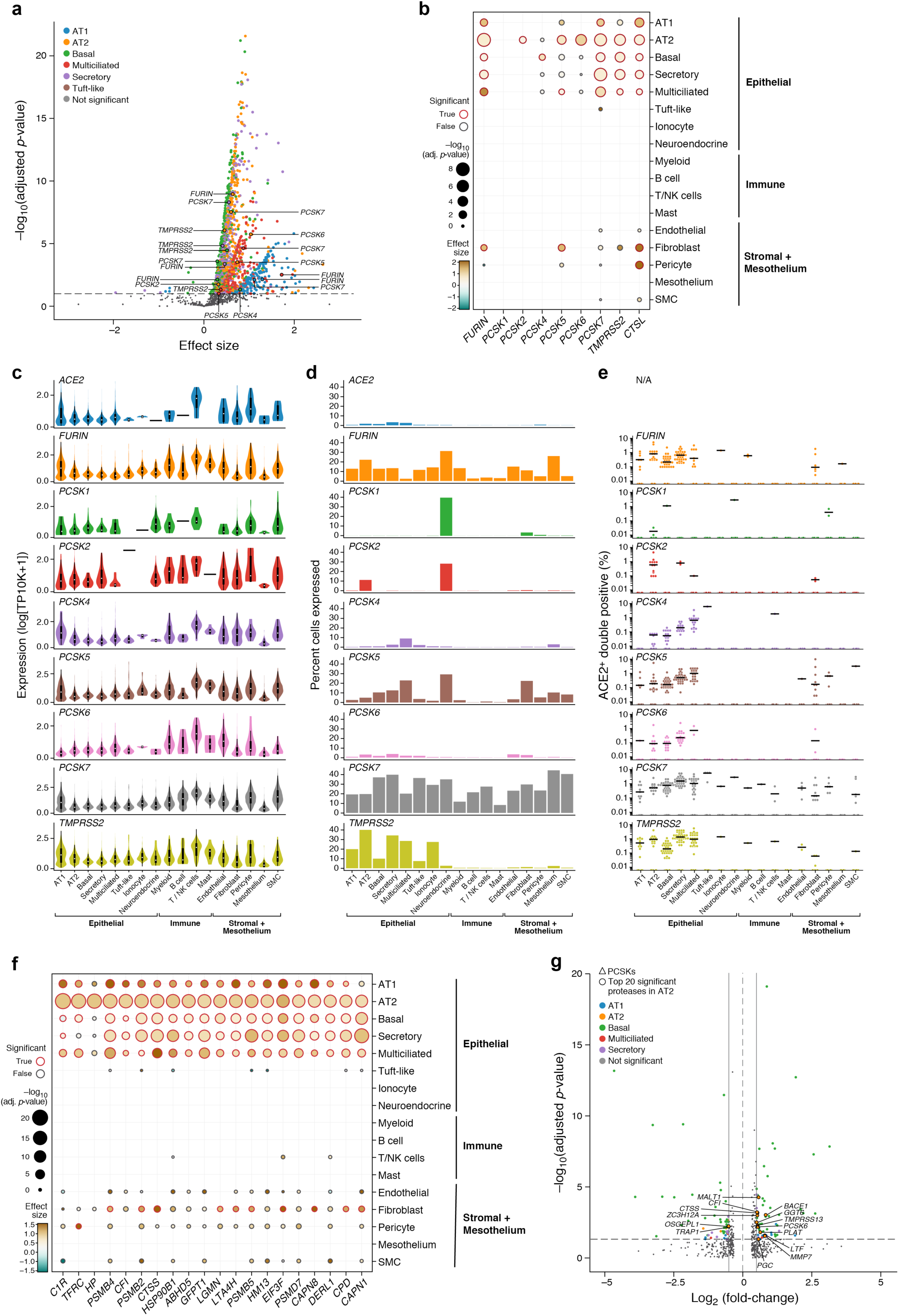
*ACE2*-protease co-expression and SARS-CoV-2 S-protein cleavage sites suggest a possible role for additional proteases in infection. **(a)** Multiple proteases are co-expressed with *ACE2* in human lung scRNA-seq. Scatter plot of significance (y axis, -log10(adjusted P value)) and effect size (x axis) of co-expression of each protease gene (dot) with *ACE* within each indicated epithelial cell type (color). Dashed line: significance threshold. *TMPRSS2* and *PCSK*s that significantly co-expressed with *ACE2* are marked. (**b**) *ACE2-protease* co-expression with *PCSKs*, *TMPRSS2* and *CTSL* across lung cell types. Significance (dot size, -log10(adjusted P value)) and effect size (color) for co-expression of ACE2 with selected proteases (columns) across cell types (rows). (**c,d**) Multiple proteases are expressed across lung cell types. (**c**) Distribution of expression (y axis) for *ACE2*, *PCSKs* and *TMPRSS2* across lung cell types (x axis). White dot: median expression. **(d)** Proportion of cells (y axis) expressing *ACE2*, *PCSK* family or *TMPRSS2* across lung cell types (x axis), ordered by compartment. **(e)** *ACE2*^+^*PCSK*^+^ dual positive cells across lung cell types. Fraction (y axis) of different *ACE2*^+^*PCSK*^+^ or *ACE2*^+^*TMPRSS2*^+^ dual positive cells across lung cell types (x axis). Dots: different individuals, line: median. **(f)** *ACE2-protease* co-expression analysis for the 20 most significant human proteases in AT2 cells. Significance (dot size, -log10(adjusted P value)) and effect size (color) for co-expression of *ACE2* with different proteases (columns) across cell types (rows). **(g)** Additional protease expression in *ACE2*^+^*TMPRSS2*^+^ dual positive cells. Significance (y axis) and fold change (x axis) of differential expression for each human protease between *ACE2*^+^*TMPRSS2*^+^ dual positive vs dual negative cells within each indicated epithelial cell types (color). The top 20 most significantly differentially expressed proteases within AT2 cells and *PCSKs* across all epithelial cell types are highlighted.

We focused on trends or indications in those cell types where both *TMPRSS2* and *ACE2* are predominantly expressed in the lung: airway epithelial cells (basal, multiciliated, and secretory cells), alveolar AT2 cells, and submucosal gland secretory cells (**Fig. 3e**). Strikingly, we find robust trends of *ACE2* expression with age, sex, and smoking status in these cell types (**Fig. 3d, Extended Data Fig. 6 and MALTE4**): *ACE2* expression increases with age in basal and multiciliated cells. *ACE2* expression is elevated in males in airway secretory cells and alveolar AT2 cells. Furthermore, we find strongly elevated levels of *ACE2* in past or current smokers in multiciliated cells (log fold change (log FC): 1.52, **Fig. 3d**). Significant associations of ACE2 expression indicate increased expression with age in AT2 (largest age effect: slope of log expression per year of 0.020) and secretory cells, and increased expression of *ACE2* in males in multiciliated cells. Further indications associate past or current smoking with decreased *ACE2* expression in AT2 cells, and increased ACE2 expression in basal cells. However, these last five indications are not robust trends and depend on the inclusion of a single dataset (**Fig. 3f**), often because that dataset contributes a large number of cells of a particular type. Specifically, when we held out the largest declined donor transplant dataset (**Supplementary Table 2**, “Regev-Rajagopal”, most cells and most samples), a declined donor tracheal epithelium dataset (“Seibold”, **Supplementary Table 2**, most donors in the smoking analysis), or a further declined donor lung dataset (“Kropski-Banovich”, **Supplementary Table 2**) respectively, the effect is no longer present (**Methods, Fig. 3f, Supplemenatry Data D1**).

The above trends and indications for sex and age were further validated in the simplified model on the full non-fetal lung dataset (**Extended Data Fig. 8, Supplementary Data D2**). With the exception of the age association in basal and secretory cells, all associations were found to be significant at a false discovery rate (FDR) threshold of 5%, confirmed by pseudo-bulk analysis, and were supported at least at the level of *indication*. Indeed, all robust trends were also supported as robust trends in the simplified model. Fitting the simplified model on the smoking data subset shows that modeling smoking status is crucial to detect the basal and secretory age association. Without modeling the effect of smoking status on basal and secretory cell *ACE2* expression, this variance is captured as uncertainty against which the age effect is evaluated as not significant.

Taking into account smoking status is of particular importance as the effect sizes associated with smoking tend to be much larger than age effects, and tend to be larger than sex effects. For example, in multiciliated cells the effect sizes assigned to smoking status, sex, and age associations with *ACE2* expression are β=1.52, β=0.51, and β=0.0107 respectively, where β represents the log FC and the slope of log expression per year.

Examining joint trends of *ACE2* and the protease genes within the same cell type, there are indications of up-regulation of both *ACE2* and *TMPRSS2* in multiciliated cells (*ACE2* indication dependent on “Seibold” dataset) in males and with age (both indications dependent on “Regev-Rajagopal” dataset). In AT2 cells, there is an indication of joint up-regulation of *ACE2* and *TMPRSS2* with age (dependent on “Regev-Rajagopal” dataset), and an indication of *ACE2* and *CTSL* down-regulation in smokers (dependent on “Kropski-Banovich” dataset). All above joint trends for age and sex covariates were confirmed on the full non-fetal lung data using the simple model without smoking covariates. In aggregate, elevated levels of cell-type specific *ACE2* and associated proteases are correlated to increasing age, in smokers, and in males.

### Distinctive changes in *ACE2^+^TMPRSS2^+^* expression patterns during human development and very low expression in infants and young children

The age associations highlighted particularly low expression in samples from very young children (newborn to 3 years old). As reports suggest that most infants and young children cases do not display severe disease^11, 74^, we inspected the subsets of studies of human development and pediatric samples from our integrated analysis **(Supplementary Table 2)**. These included cells from 12 first trimester samples (8 donors) of fetal lung (8.5 weeks post conception; wpc), 29 fetal lung samples (26 donors) from the second trimester (13-24 weeks)^68^, 13 lung samples (10 donors) spanning from third trimester premature births (n=4), full term newborns (n=1), ∼3-month old (n=4), 3-year-old (n=3), and 10-year-old (n=1) children. Because the number of samples here is small, all observations must be interpreted with caution.

The extent of *ACE2* expression in lung cells changes during development (**Fig. 3g, Supplementary Table 5**). There are dual-positive cells present in the very early first trimester lungs by scRNA-seq, and some *ACE2* expression in epithelial cells in the second trimester samples (**Extended Data Fig. 3a,b**). Notably, spatial transcriptomics of 8.5 pcw fetal lung did not capture any *ACE2* expression (data not shown). In lungs from third trimester pre-term births, *ACE2* expression is high, with *ACE2^+^TMPRSS2^+^* cells observed in alveolar AT2 cell populations (**Extended Data Fig. 3b**).

Strikingly, *ACE2* expression is very low in normal lungs of newborns, the one ∼3 months’ lung sample, and the 3-year-old lungs (**Fig. 3g**). This is further supported by single-cell chromatin accessibility by transposome hypersensitive sites sequencing (scTHS-Seq^75^) from human pediatric samples (full gestation, no known lung disease) collected at day 1 of life, 14 months, 3 years, and 9 years (n=1 at each time point) (**Extended Data Fig. 9a**). *ACE2* gene activity scores (**Methods**), when present, were in the AT1/AT2 population, but no signal was present at birth, it was low in the 9-year-old and 3-year-old sample, and higher in the 14 month-old (**Extended Data Fig. 9b-d**). Notably, immunohistochemistry (IHC) of the ∼3 month-old infant lung also showed fewer ACE2-immunoreactive AT2 cells (**Extended Data Fig. 5d**).

We also assessed whether *ACE2*, *TMPRSS2*, and *CTSL* are expressed in the human placenta during pregnancy, using data from three published scRNA-seq studies: two from the first trimester (64,000 cells and 14,000 cells) and one from full-term placenta (20,518 cells)^76–78^. *ACE2* was expressed (1.4%) in maternal decidual/stromal cells, maternal pericytes, and fetal extravillous trophoblasts, cytotrophoblasts, and syncytiotrophoblast in both first-trimester and term placenta (**Fig. 1d**). Note that there was little expression of *TMPRSS2* (0.2%) in the placenta and accordingly few *ACE2^+^TMPRSS2^+^* dual-positive cells (as we previously reported^45^). However, *CTSL* is expressed in most cells (56%) in the maternal-fetal interface, and there are *ACE2^+^CTSL^+^* dual-positive cells (1.3%) among maternal decidual/stromal cells, pericytes, and fetal trophoblasts (HLA-A, B negative) in both first-trimester and term placentae. Overall, these patterns may be important in understanding why children are more resistant to COVID-19 and and in considering risk during pregnancy.

### An immune gene program associated with *ACE2^+^TMPRSS2^+^* expression in airway, lung and gut cells

Our Human Lung Cell Atlas analyses have revealed immune signaling genes that co-vary with *ACE2* and *TMPRSS2* in airway and lung cells^45, 46^. These analyses identified antiviral response genes that are enriched in *ACE2^+^TMPRSS2^+^* cells (*e.g., IDO1, IRAK3, NOS2, TNFSF10, OAS1, MX1*), and suggested that *ACE2* itself is interferon regulated^45, 46^.

To explore such gene programs in a broader context, we identified signatures for dual-positive *ACE2^+^TMPRSS2^+^* cells compared to dual-negative *ACE2^-^TMPRSS2^-^* cells in the nasal epithelium, lung, and gut (**Supplementary Tables 6, 7 8**) with two complementary approaches. The first aimed to find features that characterize programs of dual positive cells that are shared by different cell types in one tissue (“tissue programs”). The second aimed to find features that are associated with dual positive cells compared to other cells of the same type, and may or may not be shared with other types (“cell programs”) (**Methods**). To infer tissue programs, we trained a random forest classifier to discriminate between dual-positive and dual-negative cells (excluding *ACE2* and *TMPRSS2*; 75:25 class balanced test-train split), generalizing across multiple cell types in one tissue, and ranked genes according to their importance scores in the classifier (**Methods**). To infer cell programs, we performed differential expression analysis between dual-positive and dual-negative cells within each cell subset. We note that *ACE2*^+^TMPRSS2^+^ cells have more unique transcripts detected (**Extended Data Fig. 10b**): this can reflect a technical confounder, biological features, or both. We conservatively controlled for these differences (by sampling dual positive and dual negative cells from matched gene complexity bins; **Methods; Extended Data Fig. 10b, Extended Data Fig. 11**). Importantly, these methods do not assume that *ACE2*^+^*TMPRSS2*^+^ cells form a distinct subset within each cell type. Rather, our goal is to leverage the variation among single cells within a single type to identify gene programs that are co-regulated with *ACE2* and *TMPRSS2* within each expressing cell subset.

Tissue programs (**Fig. 4a, Extended Data Fig. 10a, Supplementary Table 7, 8**) were enriched in several pathways related to viral infection and immune response (see **Fig. 4b, Extended Data Fig. 10a** for visualization of selected genes, and **Supplementary Tables 6** for the full list). These include phagosome structure, antigen processing and presentation, and apoptosis. Among the tissue program genes we highlight: *CEACAM5* (lung, nasal, gut programs) and *CEACAM6*^79^ (lung), surface attachment factors for coronavirus spike protein; *SLPI* (lung, nasal), a secreted protease inhibitor that is associated with virus resistance^80^; *PIGR* (lung, gut), the polymeric immunoglobulin receptor that may promote antibody-dependent enhancement via IgA^81^; and, *CXCL17* (lung, nasal), a mucosal chemokine that attracts dendritic cells and monocytes to the lungs^82^. In addition, the tissue programs yielded multiple genes that are associated with cholesterol and lipid metabolic pathways and endocytosis (*DHCR24*, *LCN2*, *FASN*); high expression of both MHC I (*B2M*, *HLA-B*) and MHC II (*HLA-DRA, DRB1*, *CTSS*, *CD74*) pathways, expanding on prior evidence that AT2 cells can serve as antigen presenting cells in infection^83^; genes indicating preparation against cellular injury through ROS inhibition, preventing proteases, or through antiviral responses (interferons, extracellular RNAse, *etc*: *PLAC8*, *TXNIP*); complement genes (*C3*, *C4BPA*); genes involved in immune modulation (*BTG1*); and, tight junction genes (*DST*, *CLDN3*, *CLDN4*).

The cell programs (**Fig. 4c, Extended Data Fig. 11a,b, Supplementary Table 6**) were enriched in many of the same genes and pathways as tissue-specific programs (**Fig. 4d, Supplementary Table 6,7,8,9,10**), and highlight a potential role for TNF signaling in ACE2 regulation. We first confirmed that the cell programs were not merely associated with the number of transcripts per cell (**Extended Data Fig. 11c**). While some genes were shared between the tissue and cell programs (*e.g.*, many virus-related genes, such as *CEACAM5*, *CXCL17*, *SLPI,* and *HLA-DRA*), the cell programs further captured unique biological functions and activities. For example, dual positive lung secretory cells differentially expressed genes involved in TNF signaling including *RIPK3*, a key regulator of inflammatory cell death via necroptosis, previously implicated in SARS-CoV pathogenesis^84^. Both lung dual positive secretory and multiciliated cells differentially expressed lysosomal genes (*MFSD8*, *CTSS*, *CTNS, CTSH*), potentially relevant for endo-lysosomal entry of coronaviruses^85^. Dual positive AT1 cell programs included genes involved in immunoproteasome (*PSMB8*, *PSMB9*, **Fig. 4c**), class I and II antigen presentation (*HLA-DMA*, *HLA-DRB5*, *HLA-DPB1*, *HLA-DRA*, *HLA-DPA1*), and phagocytosis. Dual-positive nasal goblet cells differentially expressed several cytokines and chemokines, including granulocyte-colony stimulating factor (*CSF3*), which may impact hematopoiesis, the recruitment of neutrophils, and inflammatory pathology; *CXCL1* and *CXCL3*, chemoattractants for neutrophils; interleukin-19 (*IL19*), which induces the production of IL-6 and TNF^86^; and CCL20, which is upregulated by TNF ^87^. The AT2 cell program included the surfactant proteins, *SFTPA1* and *SFTPA2*; the IL-1 receptor (*IL1R1*), which may promote antiviral immune responses (below); and, multiple components of MHC-II (*e.g., HLA-DPA1*, *HLA-DPB1*), congruent with a role in antigen presentation.

Cell programs from multiple tissues (**Fig. 4c,d**) included genes related to TNF signaling (*e.g., BIRC3*, *CCL20*, *CXCL1*, *CXCL2*, *JUN*, *NFKB1*), raising the possibility that anti-TNF therapy may impact the expression of *ACE2* and/or *TMPRSS2*. Consistent with this hypothesis, *ACE2* expression in enterocytes was significantly lower in ulcerative colitis patients treated with anti-TNF compared to untreated patients (mean = 0.22 and 0.13 log_2_(transcripts per 10,000 (TP10K)+1) in treated *vs.* untreated; adjusted *P* < 1e-3). However, we could not control for many important features, including disease severity, which is strongly associated with anti-TNF treatment, raising the need for future work. Some of the genes are targets of known drugs^88^. For example, dual-positive lung secretory cells expressed, in addition to ACE2 (targeted by ACE inhibitors), other drug targets, including C3, HDAC9, IL23A, PIK3CA, RAMP1, and SLC7A11. Other program genes were shown to interact with SARS-CoV-2 proteins via affinity purification mass spectrometry ^89^. Among those was GDF15, which was identified as a putative interaction partner for the SARS-CoV-2 protein Orf8^89^, is a central regulator of inflammation^90^, and was a member of the dual-positive cell programs of both lung basal cells and nasal multiciliated cells.

Some program genes may be particularly related to COVID19 pathological features and may indicate putative therapeutic targets. For example, *MUC1* is especially highly induced in dual-positive cells (in tissue and specific cell programs), which may be associated with respiratory secretions^91^. Importantly, the lung tissue and gut enterocyte programs include the gene encoding the IL6 co-receptor (*IL6ST*), and the AT2 cell program includes *IL6*. IL6 signaling has been implicated in uncontrolled immune responses in the lungs of COVID19 patients, elevated serum IL6 levels are associated with the need for mechanical ventilation^92^, and anti-IL6R antibodies (tocilizumab) are being tested for clinical efficacy in COVID-19 patients. Indeed, *IL6ST* and *IL6* are higher in dual positive *vs*. dual negative AT2 cells (**Extended Data Fig. 11d**), although *IL6* expression is relatively low in these cells from healthy tissue. Additional cell types, such as heart pericytes, are enriched for cells with co-expression of *ACE2* with *IL6R* or *IL6ST* (**Extended Data Fig. 12**).

The immune-like features of *ACE2^+^* epithelial cells are also reflected in the regulatory features of the *ACE2* locus by scATAC-Seq (**Fig. 4f**). Epithelial cells (AT1, AT2, secretory, multiciliated, ionocytes, and neuroendocrine cells) with open chromatin at the ACE2 locus show high motif activity scores^93^ for interferon-regulatory factors such as *IRF1/7* (MA0050.1, adj p-val=0.02, MA0772.1, adj p-val=0.08), *STAT1*::*STAT2* (MA0517.1, adj p-val=0.15), *FOXA1* (MA0148.2, adj p-val=0.24) and *FOXD2* (MA0847.2, adj p-val=0.27), compared to epithelial cells with no sign of open chromatin at the *ACE2* locus, though the difference is statistically significant (adj p-val <0.05) only for *IRF1*. Interestingly, Forkhead Box Transcription Factors have been implicated in regulation of *ACE2* transcription^94^. Note that because epithelial cells with an accessible *ACE2* locus tend to have a higher number of fragments in peaks than cells with inaccessible *ACE2* (**Extended Data Fig. 11f**), consistent also with higher UMIs in scRNA-seq, some of the cells with inaccessible *ACE2* could be false negatives, reducing our power.

Previous studies in the healthy lung predicted that interactions between AT2 cells and myeloid-lineage macrophages may be important for immune regulation and surfactant homeostasis^95^. To explore this possibility, we predicted interactions between AT2 cells (in general, or *ACE2^+^TMPRSS2^+^* dual-positives) and myeloid cells (**Methods**^96^), using our large declined donor transplant dataset (“Regev/Rajagopal”; 41 samples, 10 patients, 2-6 locations each). AT2 cells and myeloid cells were present in lung lobes samples from all 10 patients, whereas samples from 5 patients contained both *ACE2^+^TMPRSS2^+^* dual-positive AT2 cells and myeloid cells. We identified significant predicted interactions involving Oncostatin M (*OSM*), an IL6-type cytokine expressed by myeloid cells^97^, with the Oncostatin M Receptor (*OSMR*) and its paralog receptor LIF Receptor Subunit Alpha (LIFR) expressed in both for AT2 cells in general, and in double positive ones). Interactions involving the complement pathway were also predicted (for all AT2 and dual positives) between Complement C3 and *C5* expressed by AT2 cells and their cognate receptor expressed in myeloid cells. Three samples had interactions between the IL1 receptor on AT2 cells and IL1B or IL1RN in myeloid cells. The IL1-receptor interactions were identified mostly (in 2 out of 3 samples) involving only dual-positive AT2 cells, suggesting a possible role of *ACE2^+^TMPRSS2^+^* dual-positive cells in IL1-mediated processes. Finally, we identified interactions between CSF1, 2, or 3 expressed in AT2 cells (including double positives) and their receptors expressed in myeloid cells. These predicted interactions further support the previously identified roles for cross-talk between AT2 and myeloid cells, such as macrophages, in immune regulation (OSM, complement, IL1) and surfactant homeostasis (CSF), as previously highlighted^95^.

### Conserved expression patterns in mouse models

We next asked whether human cell types of interest were present in animal models. While such analyses cannot address molecular compatibility (due to sequence variation in ACE2 across species, as shown for lower compatibility of SARS-CoV and mouse ACE2^98^), they can help determine if dual-positive cells are present in commonly employed models, and if their characteristics, proportions, and programs are similar to those of their human counterparts. In a separate study^46^, our lung network showed strong similarities to the human data in a macaque model. Here, we focused on the more distant, but commonly used, mouse model.

*Ace2^+^Tmprss2^+^* and *Ace2^+^Ctsl^+^* dual-positive cells were present primarily in club and multiciliated cells in the airway epithelia of healthy mice^99^ (*Ace2^+^Tmprss2^+^* club 5.1% [4.7%, 5.4%] and multiciliated 2.8% [2.3%, 3.5%], *Ace2^+^Ctsl^+^* club 11.1% [10.6%, 11.6%] and multiciliated 3.5% [2.9%, 4.3%]), consistent with the expression patterns found in human airways (**Fig. 5a**). Furthermore, *Ace2* expression increased over a 2-month time-course of healthy mouse aging in both club (*p*=1.16e-06) and goblet (*p*=0.01911) cells (**Fig. 5a**). The proportion of *Ace2^+^Tmprss2^+^* dual-positive cells did not significantly increase with age during this time course (data not shown), but the proportion of *Ace2^+^Ctsl^+^* dual-positive cells significantly increased in club cells during this time-course (**Fig. 5b**). Interestingly, the mice were aged between 2-4 months, a 2-month period that is reported to reflect the maturation period from early to mature adults^100^. Examining bulk RNA-Seq profiles of sorted populations of alveolar AT2 cells (SFTPC^+^), airway basal cells (KRT5^+^), alveolar endothelial cells (CD45-CD31^+^), alveolar epithelial cells (Epcam^+^), whole lung and whole trachea from a KRT5-CreER/LSL-TdTomato/SFTPC-eGFP transgenic mouse model, and across tissues from ENCODE, showed that *Ace2, Tmprss2* and *Ctsl* are expressed in sorted AT2 cells, whole trachea and whole lung, as well as in stomach, intestine, kidney and bladder.

In human smokers, statistical modeling uncovered a robust trend of increased *ACE2* expression in airway epithelial cells, while expression in AT2 cells was reduced (**Fig. 3d, Extended Data Fig. 6**). To experimentally confirm these findings, we examined cell profiles from mice exposed daily to cigarette smoke for two months, followed by scRNA-seq of whole lungs (**Fig. 5c**). Epithelial specific expression patterns of mouse *Ace2* and the *Ace2^+^Tmprss2^+^* and *Ace2^+^Ctsl^+^* dual-positive cells were largely consistent with the human data (**Fig. 5d**). Upon smoke exposure, there was a significant increase in *Ace2^+^* airway secretory cell numbers, while the fraction of *Ace2^+^* AT2 cells was unaltered (**Fig. 5e**). Moreover, the expression levels of *Ace2* were significantly increased in airway secretory cells (**Fig. 5f**), but not in AT2 cells (**Fig. 5g**). This was in agreement with bulk RNA-seq of mouse lungs exposed to different doses of cigarette smoke^101^, in which *Ace2* levels increased in a dose-dependent manner by daily cigarette smoke over 5 months (**Fig. 5h**). Notably, the COVID-19 relevant proteases *Tmprss2* and *Ctsl* were also significantly increased by smoke exposure in mice (**Fig. 5i,j**). Thus, mouse smoking data shows similar trends as observed in humans and experimentally confirms the association of *ACE2* levels with smoking.

We also compared the patterns between the mouse and human placenta, analyzing *Ace2*, *Tmprss2*, and *Ctsl* expression across 100,000 cells from scRNA-seq data during mouse placenta development from embryonic days 9.5 to 18 (Shu et al., unpublished). We find *Ace2^+^Tmprss2^+^* dual-positive cells (1.4%) in a large fraction of fetal trophoblasts with strong epithelial signatures. *Ace2^+^Tmprss2^+^* dual-positive cells express signatures of AT2 cells and hepatocytes, and many also express *Ctsl*. *Ace2*^+^*Ctsl*^+^ dual-positive cells (3.4%) are also present among fibroblasts, stromal cells, and fetal trophoblasts in both mice and humans (**Fig. 5k, Extended Data Fig. 13**). Notably, while *ACE2*^+^*CTSL*^+^ dual-positive fibroblasts and stromal cells in humans are of maternal origin, *Ace2*^+^*Ctsl*^+^ dual-positive fibroblasts and stromal cells are of fetal origin in mice.

### Expression patterns of additional proteases that may be relevant to infection

TMPRSS2 has been demonstrated to mediate SARS-CoV-2 infection *in vitro*^37, 44^, but SARS-CoV-2 also infects cells in the absence of TMPRSS2^37^. Thus, additional proteases likely play roles in proteolytic cleavage events of spike and other viral proteins that underlie entry (fusion) and egress. To systematically predict proteases potentially involved in SARS-CoV-2 pathogenesis, we tested for co-expression of each of 625 annotated human protease genes^102^ with *ACE2* in the large declined donor transplant dataset (“Regev/Rajagopal”) from 10 patients. The analysis recovered *TMPRSS2* as one of the significantly co-expressed in multiple lung epithelial cell types (**Fig. 6a, Supplementary Table 11, 12**). In addition, multiple members of the proprotein convertase subtilisin kexin (*PCSK*) family were also significantly co-expressed with *ACE*2 in both proximal and distal airway epithelial cells (**Fig. 6a,b**), including *FURIN, PCSK2, PCSK5, PCSK6 and PCSK7* in AT2 cells. Proprotein convertases have known roles in coronavirus S-protein priming^43, 103, 104^. We obtained similar results in an independent dataset of 182,952 cells from 40 donors (**Extended Data Fig. 14a,b, “aggregated lung”**).

To further investigate the role of proprotein convertases as candidates for SARS-CoV-2 S-protein processing we analyzed the SARS-CoV-2 spike protein sequence. Multiple sequence alignment of S-protein sequences of SARS-CoV-2 and other beta-coronaviruses revealed a polybasic insert at the S1/S2 junction present only in SARS-CoV-2 spike (**Extended Data Fig. 14c**). While polybasic sites are found in multiple members of betacoronavirus lineages A and C (*e.g.*, MERS-CoV), SARS-CoV-2 is the only known member of lineage B harboring a polybasic motif in the S1/S2 region (**Extended Data Fig. 14c**). As previously reported, this polybasic sequence corresponds well to cleavage motifs of multiple PCSK family proteases (**Extended Data Fig. 14d**)^105–107^, and has a high probability for its PCSK-mediated cleavage (at amino acid 685) (by ProP and PROSPERous^108, 109^) as well as additional sites including the S2’ position (at amino acid 815), which would release predicted fusion-mediating peptides (**Extended Data Fig. 14e**)^110^.

We next examined PCSKs expression and co-expression with *ACE2* across lung cell subsets (**Fig. 6c, Extended Data Fig. 14f**). *FURIN*, *PCSK5* and *PCSK7* were broadly expressed across multiple lung cell types, and *PCSK1* and *PCSK2* were largely restricted to neuroendocrine cells, as previously reported^107^, with *PCSK2* further detected in 11.2% of AT2 cells (**Fig. 6d, Extended Data Fig. 14g**). In many cell subsets we observed dual expression with ACE2 at fractions comparable to or higher than those of *ACE2*^+^*TMPRSS2*^+^ cells (**Fig. 6e, Extended Data Fig. 14h**). These include AT2 cells (*ACE2^+^TMPRSS2^+^*, *ACE2*^+^*FURIN*^+^ and *ACE2*^+^*PCSK2*^+^ at 0.90%, 0.78%, and 0.56%, **Fig. 6e**); multiciliated cells in the proximal airway (*ACE2^+^TMPRSS2^+^*, *ACE2*^+^*FURIN*^+^, *ACE2*^+^*PCSK7*^+^, and *ACE2*^+^*PCSK5*^+^ at 0.91%, 0.37%, 1.02%, and 0.91%), and basal cells (*ACE2^+^TMPRSS2^+^*, *ACE2*^+^*FURIN*^+^, and *ACE2*^+^*PCSK7*^+^ at 0.19%, 0.21%, and 0.74%). Co-expression is present across tissues in addition to the lung (**Extended Data Fig. 14i,j**), including the liver, ileum, kidney and nasal airways, with the highest percentages of *ACE2*^+^*PCSK*^+^ dual positive cells in nasal airways (*ACE2*^+^*PCSK7*^+^ 1.36%, *ACE2*^+^*FURIN*^+^ 0.67%), bladder (*ACE2*^+^*PCSK5*^+^ 0.45%) and testis (*ACE2*^+^*PCSK7*^+^ 0.41%).

Because different host proteases may contribute to different stages of the viral life cycle^107, 111^, we also examined the prevalence of *ACE2^+^TMPRSS2^+^PCSK*^+^ triple-positive cells (TPs) in the lung dataset. *ACE2^+^TMPRSS2^+^PCSK7*^+^ were the main triple positive cells in multiciliated (0.75%) and secretory cells (0.72%) of proximal airways, and *ACE2^+^TMPRSS2^+^FURIN*^+^ TPs were the most common within AT2 cells (0.36%) (**Extended Data Fig. 14k**). Finally, when we examined all known human proteases for co-expression with *ACE2* in major lung epithelial cell types (**Fig. 6f**), we recovered cathepsins (*CTSB*, *CTSC*, *CTSD*, *CTSL*, *CTSS*), proteasome subunits (e.g. *PSMB2, PSMB4, PSMB5*), and complement proteases (*C1R*, *C2*, *CFI*) (**Fig. 6f, Extended Data Fig. 15**), the latter also captured in our programs above (**Fig. 4**).

## DISCUSSION

We performed integrative analyses of single-cell atlases in the lung and airways and across tissues to identify cell types and tissues that have the key molecular machinery required for SARS-CoV-2 infection. We then examined the relationship between specific cell types and three key covariates -- age, sex, and smoking status -- that have been related to disease severity. We further used the scale of these integrated atlases to identify gene programs in major epithelial cell subsets that may be infected by the virus, and search for other potential accessory proteases. Our hope is that this extensive analysis and resource will help with hypothesis generation (and refutation) towards better understanding of the molecular and cellular basis of COVID-19 infection, and the identification of putative therapeutic avenues.

Our cross-tissue analysis substantially expands on our ^45, 46, 58, 112^ and others’ ^113–115^ earlier efforts, allowing us to identify cell subsets across diverse tissues that may be implicated in virus transmission, pathogenesis, or both. Focusing on pathogenesis, in addition to key subsets in the lung, airways and gut, we identified *ACE2^+^* cells that co-express either *TMPRSS2* or *CTSL* in diverse organs, many of which have been associated with severe disease. These include epithelial cells in the liver, kidney, pancreas, and olfactory epithelium, cardiomyocytes, pericytes and fibroblasts in the heart, and oligodendrocytes in the brain. For example, the presence of double positive cardiomyocytes and cardiac pericytes and fibroblasts may provide a pathological basis for the cardiac abnormalities noted in COVID-19 patients including elevated troponin, a signature of cardiomyocyte injury, myocarditis, and sudden cardiac death^5^. As the co-expression of genes involved in SARS-CoV2 infection are highest in cardiac pericytes in healthy heart, damage to vascular beds may trigger troponin release in otherwise normal hearts. Moreover, as myocardial ACE expression is increased in patients with existing cardiovascular diseases (Tucker et al. companion manuscript^58^), SARS-CoV2 infection may result in greater damage to cardiomyocytes, and account for greater disease acuity and poorer survival in these patients.

One intriguing clinical observation is that some COVID-19 patients display an array of neurologic symptoms^116–118^, reported as seizures and acute necrotizing encephalopathy, similar to that previously observed following other infections such as influenza^119, 120^. Neuroinflammation could result from direct viral infection of the brain, or a systemic cytokine storm ^121–123, 124–126^. Direct viral invasion of SARS-CoV and MERS-CoV is observed in multiple brain regions in human patients and mouse models^124, 125^, consistent with widespread *ACE2* expression in numerous brain cell types. Furthermore, SARS-CoV has been shown to infiltrate the brain via the olfactory epithelium–olfactory bulb axis^127^; olfactory transmission for SARS-CoV-2 has been recently proposed^128^. Other possible transmission routes could be through the infection of *ACE2^+^TMPRSS2^+^* enteric neurons synapsing with vagal afferents, or entry through blood-CNS interfaces such as the choroid plexus or meninges^121, 129–131^. Profiling immune cells at these sites after infection is an important future step to better understand how the viral response may lead to encephalitis. One intriguing possibility is that encephalitis might arise as an autoimmune response to myelin antigens expressed by infected cells. Antibodies against peptides of myelin proteins have been clinically shown to be associated with autoimmune encephalitis and seizures^132–134^, and myelin peptides are targets of T cells^135^ in demyelinating inflammatory neurological diseases such as acute demyelinating encephalomyelitis, Guillain-Barre Syndrome, and multiple sclerosis^135–142^. Oligodendrocytes, the myelin-producing cells of the CNS, are the main *ACE2^+^TMPRSS2^+^* cell type in the brain, the myelin transcriptional regulator *MYRF* was enriched in certain *ACE2^+^TMPRSS2^+^* cell types as noted above, and myelin proteins *MOG* and *MBP* were co-expressed in numerous *ACE2+* clusters across organs. *MYRF* and *MBP* were significantly differentially expressed in *ACE2^+^TMPRSS2^+^* subsets of the lung and gut (**Supplementary Table 10**); myelin targeting Th17 cells trained in the gut are able to infiltrate the CNS in a mouse model of experimental autoimmune encephalomyelitis^143^. Taken together, the expression of myelin proteins across multiple *ACE2^+^TMPRSS2^+^* cells may hypothetically contribute to antigen presentation and autoimmune response in the context of viral infection. A small number of COVID-19 patients were reported to have Guillain-Barre Syndrome^144^, with a demyelinating process in some; this is consistent with observations of Guillain-Barre Syndrome following other viral infections such as Zika, influenza, and Epstein-Barr virus^139, 145, 146^. Whether anti myelin specific immunity can be induced by virus infected cells expressing myelin proteins remains an area for future study.

Our meta--analysis of scRNA-seq across studies provided the required statistical power to uncover population-level signals at a molecular level and at single-cell resolution. We found that the SARS-CoV-2 receptor and associated proteases were up-regulated in airway epithelial and AT2 cells with age and in males, an association that may shed light on the marked increase in mortality with age. Furthermore, *ACE2* was up-regulated in airway epithelial cells (basal and multiciliated cells) in past or present smokers, but down-regulated in their AT2 cells; we have also confirmed this in an experimental system in a mouse model. These contrasting smoking associations show the importance of the single-cell resolution, as the down-regulation in AT2 cells will be masked by the airway epithelial signal, leading to loss of association or misinterpretation of seemingly consistent ACE2 signals in bulk RNA-Seq^147^. Importantly, *ACE2* is particularly lowly expressed in young pediatric samples, also mirrored by lack of chromatin accessibility in the ACE2 locus. *ACE2* expression is known to be regulated in complex ways across different tissues and may be affected by both common therapies (ACEi/ARB) (Tucker et al., companion manuscript^58^), and during infection^46^. Moreover, both higher *ACE2* expression per cell and a higher fraction of *ACE2*^+^ cells can in principle have implications for infection, but may have conflicting effects on pathogenesis, as *ACE2* knockout mice show more severe ARDS upon lung injury because of its role in the renin-angiotensin pathway, which seems to protect from consequences of lung injury and inflammation^148^ (for potential roles of this pathway on CoV infection (pre-COVID) see the review in ^149^). As SARS-CoV2 binding will lead to internalization and therefore downregulation of ACE2 on the cell surface, the protective function of ACE2 via its proteolytic processing of Angiotensin-II may be lost. Thus, the smoking mediated downregulation of ACE2 in AT2 cells may not protect cells from being infected but rather may increase ARDS due to more severe loss of ACE2 from the cell surface upon infection. Other confounders, including ACEi which are not available to our meta-analysis may further impact our results.

To the best of our knowledge, this study is the first single-cell meta-analysis (in any setting). To perform this meta-analysis, we used a model that included both the tested covariates (age, sex, and smoking status), technical covariates (dataset and the number of UMIs per cell), and several interaction terms. Including these interaction terms was crucial, as omission resulted in increased background variation and reversed effect estimates. Likewise, modeling the smoking status of a donor was important to reduce background variation and account for the unbalanced distribution of covariates in the dataset. For example, while we have similar numbers of male ascertained smokers and non-smokers (21 and 20 donors), there are three times as many female ascertained non-smokers as female smokers (27 and 9 donors), which is reflective of this bias in the population^150^. The addition of these terms increases the complexity of the model. Indeed, only one dataset (“Seibold”) had sufficient numbers of donors of various ages, sex, and smoking status to fit the full model. Thus, performing the meta-analysis was only possible due to the aggregation of a large number of healthy single-cell datasets enabled by the HCA Lung Biological Network and a community-wide effort.

A limitation of our expression model is that each cell is treated as an independent observation. Thus, the significance of association with traits such as sex, age, and smoking status may show inflated p-values, especially where the associations are determined from few donors. In this case, the variation between cells from a single donor dominates the variation between donors, background variation is underestimated and effect significance can be overestimated. Aggregating many datasets allows us to counteract this effect, yet p-value inflation may occur in cell types that are not as commonly shared across datasets. Our main conclusions are drawn on airway epithelial and AT2 cells, which are distributed widely across datasets and are modeled on the basis of many donors. Furthermore, we have confirmed significant associations by pseudo-bulk analysis and by holding out datasets. This confirmation ensures that associations are consistent when only considering donor variation, and we are aware if these associations are dataset dependent (often when one dataset is a particularly major source of a given cell type). Models that account for both single-cell count distributions, and population structure in the data have the potential to improve future meta-analyses across single-cell atlases.

Having a cell type annotation with consistent resolution across datasets was instrumental for analyzing the association with clinical covariates up to the resolution of basal, secretory, or multiciliated cells. These cell type labels still aggregate over considerable diversity, which is the subject of ongoing scientific research. Importantly, the labeled subtypes of these cell clusters differ between datasets. Thus, in those cases where individual associations depend on a particular dataset not being held out, it may be the case that these associations become more robust at a higher level of cell type annotation. Future high-resolution cell annotation efforts have the potential to further consolidate our single-cell meta-analysis results.

In addition to modeling associations between gene expression and clinical co-variates, we also examined whether the proportion of *ACE2^+^TMPRSS2^+^* cells per sample is associated with age or sex. While we can observe a trend of double positive cell proportions increasing with age (**Extended Data Fig. 3a**), the high compositional diversity across samples and studies (**Fig. 3a, Extended Data Fig. 16**), the potential confounders (total counts, dataset), and limited sample numbers are prohibitive to modeling these associations. Further metadata that describe the sample diversity such as harmonized annotations on anatomical location, sampling methods, and sample processing can help to capture this heterogeneity in *ACE2^+^TMPRSS2^+^* cell proportion models.

The expression of *ACE2* and *TMPRSS2* in lung, nasal and gut epithelial cells is associated with expression programs with many shared features, involving key immunological genes and genes related to viral infection, raising many hypotheses for future studies, especially as more patient tissue samples are analyzed in the coming months. In the lung, epithelial cells express *IL6*, *IL6R* and *IL6ST*, which raises the hypothesis that infection may trigger cytokine expression from these cells and contribute to uncontrolled immunological responses. The immune-like programs in these cells are further reinforced by the accessibility of STAT and IRF binding sites in scATAC-Seq data, consistent with another study from our network showing the role of interferon in regulating *ACE2* expression in epithelial cells^46^. Notably, scRNA-Seq analysis of immune cells from bronchoalveolar lavage fluid of COVID-19 patients identified high activity of transcription factors such as STAT1/2 and IRF1/2/5/7/8/9 in macrophage states increased in severe COVID-19 patients^151^. Other hypotheses for future studies include lysosomal genes in dual positive lung secretory and multiciliated cells, which may be consistent with putative “viral entry” cells, and *RIPK3* expression in the cell programs of airway cells, which opens the hypothesis of necroptosis initiating a pro-inflammatory response. Interestingly, we observed relatively high enrichment of *ACE2* in secretory cell types (mucous cells and AT2 cells). We speculate that viruses may take advantage of the rich secretory pathway components in these cells for their efficient dispersal. Additionally, SMGs of the airways are recently shown to serve as reservoirs of reserve stem cells ^152, 153^. Therefore, we also speculate that SMGs similarly may serve as reservoirs for viruses where they can escape from muco-ciliary transport and mechanical expulsion associated with severe cough in the airway luminal surface.

The gene programs of AT2 cells can also contribute to cross talk with alveolar macrophages. Our cell-cell interaction analysis suggests that AT2 cells engage with alveolar macrophages through Oncostatin M, CSF, IL1 and complement pathways, suggesting therapeutic hypotheses. The complement pathway is particularly intriguing in the context of COVID-19. First, viral protein glycosylation is a known trigger for the lectin pathway (LP) of the proteolytic complement cascade, such that in addition to classical complement activation via antibody complexes, other coronavirus glycoproteins can be recognized by LP-inducing host collectin proteins^154, 155^. Moreover, excessive complement activation resulting in acute lung injury and cytokine storms were also implicated in the pathogenesis of SARS^156^, and complement inhibition using the anti-C5 antibody eculizumab^157^ is currently evaluated as anti-inflammatory experimental emergency treatment for severe COVID-19 in clinical trials (ClinicalTrials.gov identifier NCT04288713). In our analysis, multiple complement pathway proteases (e.g. *C1R*, C2, *CFI*) are co-expressed with *ACE2* across different lung cell subsets (**Fig. 6f,g, Extended Data Fig. 15**) and complement inhibitory factor *CD55* and complement protease *C3* were preferentially expressed by *ACE2^+^TMPRSS2^+^* DPs within lung tissue, in multiciliated and secretory cells, respectively (**Fig. 4a,c, Supplementary Table 6, Supplementary Tables 7,8**). Moreover, cell-cell interaction analysis predicted cross-talk between AT2 cells expressing complement proteins C3 and C5 and macrophages expressing cognate receptors. AT2 cell expression of negative complement regulators *CFI* and *CD55* might represent a strategy for SARS-CoV-2 to at least partially escape complement surveillance.

Finally, to explore therapeutic hypotheses related to disruption of viral processing via protease inhibition, we explored the expression of other proteases across our integrated atlases. Although a multitude of different SARS-CoV-2 features likely account for its high pathogenicity and transmissivity, it has been speculated that the PRRAR loop might contribute to increased COVID-19 severity. Introduction of similar polybasic cleavage sites into avian influenza viruses and human coronaviruses was shown to render them more pathogenic, increasing mortality and viral spread^158, 159^. One hypothesis is that acquisition of a PCSK cleavage site would expand the number of cell types that can be directly infected by SARS-CoV-2. A recent report has started to address expression of *FURIN* in cells expressing SARS-CoV-2 host factors *ACE2* or *TMPRSS2*^47^, and FURIN activity is inhibited by Guanylate-binding proteins (GBPs), a group of interferon-stimulated genes, in order to restrict viral envelope processing^160^. However, the highly overlapping recognition sequence of PCSK family members (**Extended Data Fig. 14d**) suggests that multiple PCSKs in addition to FURIN could mediate cleavage at the S1/S2 PRRAR motif (**Extended Data Fig. 14e**). Our expression analysis confirms that PCSK family members, in particular *FURIN*, *PCSK5* and *PCSK7,* are more broadly expressed than *TMPRSS2* across lung cell types (**Fig. 6d**), as well as across tissues (**Extended Data Fig. 14i**). In the lung, we note the higher proportion of *ACE2*^+^*PCSK7^+^* basal cells and *ACE2*^+^*PCSK7*/*5^+^* fibroblasts (**Fig. 6e, Extended Data Fig. 14h**). Interestingly, the host interactome of SARS-CoV-2 further suggests interaction of viral proteins with PCSK6^89^, which also showed significant *ACE2* co-expression in AT2 cells (**Fig. 6b, Extended Data Fig. 14b**). Moreover, because PCSK localization is detected in different membrane compartments along the secretory and endocytic pathways^107^, it is conceivable that PCSKs could process SARS-CoV-2 S-proteins at different stages of the viral life cycle. Moreover, further analysis is required to assess the extent to which SARS-CoV-2 relies on proteolytic activity provided in trans either by neighboring cells or extracellularly localized proteases^111^. Altogether, this could provide SARS-CoV-2 with an immense flexibility in different entry and egress pathways.

Taken together, our analyses provide a rich molecular and cellular map as context for the transmission, pathogenesis, clinical associations, and therapeutic hypotheses for COVID-19. As new single cell atlases will be generated from COVID-19 tissues and experimental models, they will help further advance our understanding of this disease.

## METHODS

### Patient samples

Sample collection underwent IRB review and approval at the institutions where the samples were originally collected. “Adipose_Healthy_Manton_unpublished” was collected under IRB 2007P002165/1(ORSP-3877). Tissue samples from breast, esophagus muscularis, esophagus mucosa, heart, lung, prostate, skeletal muscle and skin referred to as “Tissue_Healthy_Regev_snRNA-seq_unpublished” were collected under ORSP-3635. Samples referred to as “Eye_Sanes_unpublished” were collected under Dana Farber / Harvard Cancer Center Protocol Number 13-416 and Massachusetts Eye and Ear Protocol Number 18-034H. Samples referred to as “Kidney_Healthy_Greka_unpublished” were collected under Massachusetts General Hospital IRB number 2011P002692. Samples referred to as “Liver_Healthy_Manton_unpublished” were collected under IRB 02-240; ORSP 1702 as well as and ORSP-2630 under ORSP-2169. Lung samples from smokers and non-smokers with suffix “Regev/Rajagopal_unpublished” were collected under Massachusetts General Hospital IRB 2012P001079 / (ORSP-3900) under ORSP-3490. Healthy and fibrotic lung samples with suffix “Xavier_snRNA-seq_unpublished” were collected under Massachusetts General Hospital IRB number 2003P000555 (CG-5242) under ORSP-3490, Medoff, 2015P000319 (CG-5145) under ORSP-3490. Pancreas PDAC samples were collected under Fernandez-del Castillo, 2003P001289 (CG-4692) under ORSP-3490 Massachusetts General Hospital IRB number Fernandez-del Castillo, 2003P001289 (CG-4692) under ORSP-3490. Samples in the dataset “Barbry” were derived from a study that was approved by the Comité de Protection des Personnes Sud Est IV (approval number: 17/081) and informed written consent was obtained from all participants involved. All experiments were performed during 8 months, in accordance with relevant guidelines and French and European regulations. No deviations were made from our approved protocol named 3Asc (An Atlas of Airways at a single cell level - ClinicalTrials.gov identifier: NCT03437122). IPF and COPD lungs in the “Kaminski” dataset were obtained from patients undergoing transplant while healthy lungs were from rejected donor lung organs that underwent lung transplantation at the Brigham and Women’s Hospital or donor organs provided by the National Disease Research Interchange (NDRI). Patient tissues relating to the dataset “Krasnow” were obtained under a protocol approved by Stanford University’s Human Subjects Research Compliance Office (IRB 15166) and informed consent was obtained from each patient prior to surgery. The study protocol was approved by the Partners Healthcare Institutional Board Review (IRB Protocol # 2011P002419). Samples in the dataset “Kropski_Banovich” were collected under Vanderbilt IRB # 060165, 171657, and Western IRB#20181836. Ethics approval number 2018/769-31. “Meyer_b” were collected under CBTM (Cambridge Biorepository for Translational Medicine), research ethics approval number: UK NHS REC approval reference number 15/EE/0152. Samples in the dataset “Linnarsson” are covered by (2018/769-31) approved by the Swedish Ethical Review Authority. Samples in the “Misharin” dataset were collected under (STU00056197, STU00201137, and STU00202458) approved by the Northwestern University Institutional Review Board. Samples in the “Rawlins” dataset were obtained from terminations of pregnancy from Cambridge University Hospitals NHS Foundation Trust under permission from NHS Research Ethical Committee (96/085) and the Joint MRC/Wellcome Trust Human Developmental Biology Resource (grant R/R006237/1, www.hdbr.org, HDBR London: REC approval 18/LO/0822; HDBR Newcastle: REC approval 18/NE/0290). The studies relating to datasets “Schultze” and “Schultze_Falk” were approved by the ethics committees of the University of Bonn and University hospital Bonn (local ethics vote 076/16) and the Medizinische Hochschule Hannover (local ethics vote 7414/2017). Fifteen human tracheal airway epithelia in the “Schultze” dataset were isolated from de-identified donors whose lungs were not suitable for transplantation. Lung specimens were obtained from the International Institute for the Advancement of Medicine (Edison, NJ) and the Donor Alliance of Colorado. The National Jewish Health Institutional Review Board (IRB) approved the research under IRB protocols HS-3209 and HS-2240. Donor lungs samples in the “Xu/Whittset” dataset were provided through the federal United Network of Organ Sharing via the National Disease Research Interchange (NDRI) and International Institute for Advancement of Medicine (IIAM) and entered into the NHLBI LungMAP Biorepository for Investigations of Diseases of the Lung (BRINDL) at the University of Rochester Medical Center, overseen by the IRB as RSRB00047606. Samples in the “Xu/Whitsett” dataset were provided through the federal United Network of Organ Sharing via the National Disease Research Interchange (NDRI) and International Institute for Advancement of Medicine (IIAM) and entered into the NHLBI LungMAP Biorepository for Investigations of Diseases of the Lung (BRINDL) at the University of Rochester Medical Center, overseen by the IRB as RSRB00047606. (**Supplementary Table 1, 2**)

### Integrated analysis of published datasets

Publicly available single-cell RNA-seq datasets were downloaded from Gene Expression Omnibus (GEO)^161^. We searched GEO for datasets that met all of the following three criteria: (1) provided unnormalized count data; (2) was generated using the 10X Genomics’s Chromium platform^162^; and (3) profiled human samples. These tissue samples spanned a wide range, including primary tissues, cultured cell lines, and chemically or genetically perturbed samples. Applying these filters increases standardization of sample as the vast majority were prepared using the same 10X Chromium instrument and Cell Ranger pipelines.

Datasets comprise of one or more samples (individual gene expression matrices), which often correspond to individual experiments or patient samples. In total, this yielded 2,333,199 cells from 469 samples from 64 distinct datasets (**Supplementary Table 1**). To allow comparison across samples and datasets, we mapped through a common dictionary of gene symbols and excluded unrecognized symbols. If a gene from an aggregated master list was not found in a sample, the expression was considered to be zero for every cell in that sample.

After all datasets were collected, we quantified the percentage of cells with >0 UMIs for both ACE2 and TMPRSS2 or ACE2 and CTSL. For further analyses with broad cell classes, we only used datasets with more than 15 double positive cells yielding 252,871 cells from 40 samples.

For integration across datasets, we used two levels of annotations. When possible, every sample was annotated with its tissue of origin based on the available metadata from GEO. We excluded any sample for which tissue was not specified. For the smaller subset of 252,871 cells we then manually annotated cell clusters with broad cell type classes using marker genes. These clusters were generated using the harmony-pytorch Python implementation (version 0.1.1 (https://github.com/lilab-bcb/harmony-pytorch ^163^) of the Harmony scRNA-seq integration method^164^ for batch correction and leiden clustering from the Scanpy package (version 1.4.5)^165^. Clusters without clear markers distinguishing types were excluded from further analysis.

Data was processed using Scanpy. Individual datasets were normalized log (UMIs/10,000 +1) by column sum and the log1p function (ln(10,000 * g_ij_ + 1) where a gene’s expression profile, g, is the result of the UMI count for each gene, *i*, for cell *j*, normalized by the sum of all UMI counts for cell *j*. This data normalization step was only used for generating the clusters and cell type annotations.

All other statistical tests for the integrated analysis were performed on the cell’s binary classification as a double positive or not. For example, for a cell to be considered ACE2+, it has >0 ACE2 transcripts. Double positive cells have >0 transcripts for both genes of interest. We used Fisher’s exact test to test for statistical dependence between the expression of ACE2 and TMPRSS2 or CTSL and corrected for multiple testing via Benjamini-Hochberg over all tests for each gene pair.

### Integrated co-expression analysis of high resolution cell annotations across tissues

We compiled a compendium of published and unpublished datasets consisting of 2,433,890 cells from 21 tissues and/or organs including adipose, bone marrow, brain, breast, colon, cord blood, enteric nervous system, esophagus mucosa, esophagus muscularis, anterior eye, heart, kidney, liver, lung, nasal, olfactory epithelium, pancreas, placenta, prostate, skeletal muscle and skin. After the harmonization of cell type annotations, *ACE2*-*TMPRSS2* and *ACE2*-*CTSL* coexpression were assessed using a logistic mixed effect model:

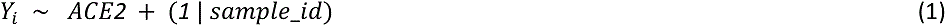

where *Y_i_* was the binarized expression level of either TMPRSS2 or CTSL, and covariates were binarized ACE2 expression in cell *i* and a sample-level random intercept.

Models were fit separately for each cell type in each dataset. In order to avoid spurious associations in cell types with very few ACE2^+^ cells and due to very low expression of ACE2, we subsampled ACE2^-^ cells to the number of ACE2^+^ cells within each cell type and discarded cell types containing fewer than 5 cells expressing either ACE2 or fewer than 5 cells expressing the other gene being tested after the subsampling procedure. The significance of the association between ACE2 and TMPRSS2/CTSL is controlled for 10% FDR using the statsmodels Python package (version 0.11.1)^166^. Data processing was performed using Scanpy Python package (version 1.4.6)^165^ and logistic models were fit using lme4 R package (version 1.1.21)^167^.

### Single-cell ATAC-Seq analysis

#### Library Generation and Sequencing

Libraries were generated using the 10x Chromium Controller and the Chromium Single Cell ATAC Library & Gel Bead Kit (#1000111) according to the manufacturer’s instructions (CG000169-Rev C; CG000168-Rev B) with unpublished modifications relating to cell handling and processing. Briefly, human lung derived primary cells were processed in 1.5ml DNA LoBind tubes (Eppendorf), washed in PBS via centrifugation at 400g, 5 min, 4C, lysed for 3 min on ice before washing via centrifugation at 500g, 5 min, 4C. The supernatant was discarded and lysed cells were diluted in 1x Diluted Nuclei buffer (10x Genomics) before counting using Trypan Blue and a Countess II FL Automated Cell Counter to validate lysis. If large cell clumps were observed, a 40µm Flowmi cell strainer was used prior to the tagmentation reaction, followed by Gel Bead-In-Emulsions (GEMs) generation and linear PCR as described in the protocol. After breaking the GEMs, the barcoded tagmented DNA was purified and further amplified to enable sample indexing and enrichment of scATAC-seq libraries. The final libraries were quantified using a Qubit dsDNA HS Assay kit (Invitrogen) and a High Sensitivity DNA chip run on a Bioanalyzer 2100 system (Agilent). All libraries were sequenced using Nextseq High Output Cartridge kits and a Nextseq 500 sequencer (Illumina). 10x scATAC-seq libraries were sequenced paired end (2 x 72 cycles).

#### Initial data processing and QC

Fastq files were demultiplexed using 10x Genomics CellRanger ATAC mkfastq (version 1.1.0). We obtained peak-barcode matrices by aligning reads to GRCh38 (CR v1.2.0 pre-built reference) using CellRanger ATAC count. Peak-barcode matrices from six channels were normalized per sequencing depth and pooled using CellRanger ATAC aggr.

The aggregated, depth-normalized, filtered dataset was analyzed with Signac (v0.1.6, https://github.com/timoast/signac), a Seurat^168^ extension developed for the analysis of scATAC-seq data. All the analyses in Signac were run with a random number generator seed set as 1234. Cells that appeared as outliers in QC metrics (peak_region_fragments ≤ 750 or peak_region_fragments ≥ 20,000 or blacklist_ratio ≥ 0.025 or nucleosome_signal ≥ 10 or TSS.enrichment ≤ 2) were excluded from the analysis.

#### Normalization and dimensionality reduction

The aggregated dataset was processed with Latent Semantic Indexing^169^, i.e. datasets were normalized using term frequency-inverse document frequency (TF-IDF), then singular value decomposition (SVD), ran on all binary features, was used to embed cells in low-dimensional space. Uniform Manifold Approximation and Projection (UMAP)^170^ was then applied for visualization, using the first 30 dimensions of the SVD space.

#### Gene activity matrix and differential motif activity analysis

A gene activity matrix was calculated as the chromatin accessibility associated with each gene locus (extended to include 2kb upstream of the transcription start site, as described in the vignette ‘Analyzing PBMC scATAC-seq’ (version: March 13, 2020, https://satijalab.org/signac/articles/pbmc_vignette.html), using as gene annotation the genes.gtf file provided together with Cellranger’s atac GRCh38-1.2.0 reference genome.

Clusters were annotated using label transfer from matching scRNA samples or by literature / expert search of marker “active” (i.e. accessible) genes. Differential motif activity analysis was performed using Signac’s implementation of ChromVAR^93^, with motif position frequency matrices from JASPAR2020^171^ (http://jaspar.genereg.net/) selecting transcription factors motifs from human (species=9606), broadly following the vignette ‘Motif analysis with Signac’ (https://satijalab.org/signac/articles/motif_vignette.html). Cells were identified as positive for *ACE2* and/or *TMPRSS2* (i.e. with the loci accessible) if at least one fragment was overlapping with the gene locus or 2kb upstream. Differential activity scores between epithelial cells positive for *ACE2* (with the above-mentioned definition of ‘positive’) and non-expressing *ACE2* was performed with the *FindMarkers* function of Seurat (version 3.1.1), using as test ‘LR’ (i.e. logistic regression) and as latent variable the number of counts in peak.

### Analysis of ENCODE bulk-RNAseq datasets

The following publically available bulk-RNAseq datasets were obtained from the ENCODE database: LID20728; LID20729; LID47030; LID47031; LID47036; LID47037; LID21040; LID21041; LID47032; LID47033; LID20870; LID20871; LID20872; LID20873; LID21183; LID21184; LID21042; LID21043; LID20920; LID20921; LID20924; LID20925; LID20821; LID20822; LID46983; LID46984; LID20819; LID20820; LID21038; LID21039; LID20732; LID20733; LID20868; LID20869; LID20922; LID20923, generated by the Gingeras lab^172^. These FASTQ datasets were aligned to the MM9 annotation build using STAR and processed using the standard Tuxedo Suite to yield a normalized FPKM matrix.

### Immunohistochemistry

Immunohistochemistry analysis was performed on 4% PFA fixed, OCT embedded tissue sections from human explant donors. Briefly, cells were permeabilized with 5% Triton-X in PBS. Slides were either not treated with antigen retrieval or antigen retrieval was performed with the Tris-Citrate buffer as needed (**Supplementary Table 4**). Slides were incubated overnight with primary antibodies at indicated concentrations (**Supplementary Table 4**) in donkey serum with 5% Triton-X in PBS. Slides were treated with Alexa-Fluor secondary antibodies mixed with DAPI at 1:500 concentration in donkey serum with 5% Triton-X in PBS for 1 hour at room temperature. Slides were mounted and imaged on a confocal microscope.

### Proximity ligation in situ hybridization (PLISH)

Proximity ligation in situ hybridization (PLISH) was performed as described previously^173^. Briefly, frozen human trachea and distal lung sections were fixed with 4.0% paraformaldehyde for 20 min, treated with protease (20 µg/mL proteinase K for lung or Pepsin for trachea for 9 min) at 37°C, and dehydrated with up-series of ethanol. The sections were incubated with gene-specific oligos (**Supplementary Table 11**) in hybridization buffer (1 M sodium trichloroacetate, 50 mM Tris [pH 7.4], 5 mM EDTA, 0.2 mg/mL heparin) for 2 h at 37°C. Common bridge and circle probes were added to the section and incubated for 1 h followed by T4 ligase reaction for 2 h. Rolling circle amplification was performed by using phi29 polymerase (#30221, Lucigen) for 12 hours at 37°C. Fluorophore-conjugated detection probe was applied and incubated for 30 min at 37°C followed by mounting in medium containing DAPI.

### Integrated analysis for association of *ACE2* and *TMPRSS2* expression with age, sex or smoking status in nasal, airways and lung cells

To assess the association of age, sex, and smoking status with the expression of ACE2, TMPRSS2, and CTSL, we aggregated 22 scRNA-seq datasets of healthy human nasal and lung cells, as well as fetal samples. Aggregation of these datasets was enabled by harmonizing the cell type labels of individual datasets within Scanpy^165^ (version 1.4.5.1). We harmonized annotations together with data contributors using a preliminary ontology generated on the basis of 5 published datasets^63–66, 174^ with 3 levels of annotations (level 1 - lowest resolution; **Supplementary Table 2**). We further harmonized metadata by collapsing the smoking covariate into “has smoked” and “has never smoked” and by taking mean ages where only age ranges were given. This endeavor produced a dataset of 1,176,683 cells in 282 samples from 164 donors (**Supplementary Data D3**). We divided the data into fetal (136,742 cells, 41 samples, 32 donors), adult nasal (32,881 cells, 12 samples, 12 donors), and adult lung (1,007,060 cells, 229 samples, 127 donors) datasets based on metadata provided.

To get an overview of sample diversity, we clustered the samples using the proportion of cells in level 2 cell types as features. Clustering was performed using louvain clustering (resolution 0.3; louvain package version 0.6.1) on a knn-graph (k=15) computed on Euclidean distances over the top 5 principal components of the cell type proportion data within Scanpy. This produced five clusters. Sample cluster labels were assigned based on metadata for anatomical location that was obtained from the published datasets and via input from the data generators.

Within adult datasets we modeled the association of age, sex, and smoking status with gene expression for ACE2, TMPRSS2, and CTSL using a generalized linear model with the log total counts per cell as offset and Poisson noise as implemented in statsmodels^166^ (version 0.11.1) and using a Wald test from Diffxpy (www.github.com/theislab/diffxpy; version 0.7.3, batchglm version 0.7.4). Specifically we used the model:

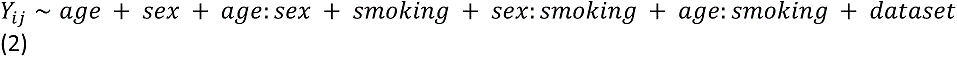

Here, *Y_i,j_* denotes the raw count expression of gene *i* in cell *j* and *age:sex*, *sex:smoking*, and *age:smoking* are interaction terms of the three modeled covariates. These terms model whether there is a difference in the smoking effect in men and women, and likewise whether the age effect is different for smokers and non-smokers. While we model these interaction terms, we only tested *age*, *sex*, and *smoking* effects individually to reduce the multiple testing burden. We included the *dataset* term to model the batch effects between the diverse datasets we obtained, and the log total counts per cell was used as an offset. Here, the total counts were scaled to have a mean of 1 across all cells before the log was taken. In order to fit this model we pruned the data to contain only datasets that have at least 2 donors and for which smoking status metadata was provided. This resulted in a dataset of 761,693 cells and 165 samples from 77 donors for adult lung data. Only a single dataset remained for adult nasal data after this filtering on which the model could not be fit. To obtain cell-type specific associations the above model was fit within each cell type for all cell types with at least 1,000 cells. We performed Wald tests over the age and sex covariates independently and corrected for multiple testing via Benjamini-Hochberg over all tests within a cell type.

As metadata on smoking status was only available for a subset of the data, we also fitted a simpler model on a larger dataset to confirm sex and age associations. The simplified model:

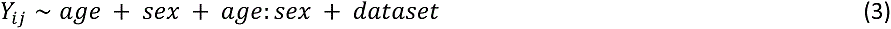

was fit on 986,342 cells in 225 samples from 125 donors of adult lung data. Again, log total counts (scaled) was used as an offset.

The above models treat cells as independent observations and thus model cellular and donor variation jointly. As donor variation tends to be larger than single-cell variation, when most cells come from few donors (either there are few donors, or few donors contribute most of the cells), this can lead to an inflation of p-values. To counteract this effect, we verified that significant associations are consistent when modeling only donor variation via pseudo-bulk analysis, and we tested whether effects are dependent on few donors by holding out datasets.

Pseudo-bulk data was generated by computing the mean for each gene expression value and *nUMI* covariate for cells in the same cell type and donor. After filtering as described above, models (2) and (3) were fit to the data. In contrast to the single-cell model, pseudo-bulk analysis underestimates certainty in modeled effects as uncertainty in the pseudo-bulk means are not taken into account. Thus, we used only effect directions from pseudo-bulk analysis to validate single-cell associations. We regarded only those associations as significant, where the FDR-corrected p-value in the single-cell model is below 0.05, and the sign of the estimated effect is consistent in both the single-cell and the pseudo-bulk analysis.

We further separated significant associations into robust trends and indications depending on the holdout analysis. A significant association was regarded as a *robust trend* if the effect direction is consistent when holding out any dataset when fitting the model irrespective of p-value. In the case that holding out one dataset caused the maximum likelihood estimate of the coefficient to be reversed, we denote this as the effect *no longer being present*, which characterized the association as an *indication*.

### Identification of gene programs using feature importances for a random forest trained to classify *ACE2^+^TMPRSS2^+^ vs ACE2^-^TMPRSS2^-^* cells

For each of the lung, nasal, and gut datasets, we labeled the cells with non-zero counts for both *ACE2* and *TMPRSS2* as dual-positive cells (DPs), and the cells with zero counts for both *ACE2* and *TMPRSS2* as dual-negative cells (DNs). Within each tissue, we identified cell types with greater than 10 DPs, and for each of these cell types, we selected the genes with increased expression (log fold change greater than 0) in DPs vs DNs (so that we focus on important “positive” features). We trained a classifier with 75:25 train:test split to classify the DPs from DNs within each of these cell types using the *sklearn* (version 0.21.3)^175^ *RandomForestClassifier* function with the following parameters: *n_estimators* set to *100*, the *criterion* as *gini*, and the *class_weight* parameter set to *balanced_subsample*. We first trained individual classifiers separately for each of the cell types, and pooled genes with positive feature importance values (using the *feature_importance*^176^ field in the trained *RandomForestClassifier* object) to train a final DP vs DN classifier across each tissue. We used the top 500 genes, as ranked by their feature importance scores, to define the signature for the gene expression program of DPs for the tissue. This procedure was carried out in lung, nasal, and gut datasets, yielding tissue-specific signatures for gene expression programs of DPs from each tissue.

For visualization purposes only, we generated network diagrams using the *networkx* (version 2.2)^177^ tool with the *ForceAtlas2* graph layout algorithm^178^. We scored genes that appeared in signatures for multiple tissues by their aggregated feature importance (using a plotting heuristic that used the sum of importance ranks for genes in individual tissues and by assigning a large valued rank (10000) to a gene that did not appear in a particular tissue) and selected the top 10 genes that were shared by each pair of tissues or shared by all tissues along with additional genes that included the ones unique to each tissue’s signature to plot in the network visualization. The GO terms enriched in the gene expression programs shared by DPs across tissues were found using *gprofiler* (version 1.0.0)^179^ using the *scanpy.queries.enrich* tool.

This analysis was performed in two ways: on the original data, as well as after accounting for differences in distribution of the number of UMIs (nUMI) per cell between DPs and DNs. This was done by binning the nUMI distribution in the DPs for each tissue into a 100 bins and then randomly sampling from the nUMI distribution for the DNs in each bin to match the distribution of the DPs in that bin. The nUMI distributions before and after the matching are shown in **Extended Data Fig. 10b.**

### Identification of gene programs enriched in DP vs. DN cells using regression

In parallel, we used a regression framework to recover gene modules enriched in DP *vs.* DN cells (**Fig. 4c,d, Extended Data Fig. 11a,b**) in the nasal, lung, and gut datasets. We first restricted our analysis to cell subsets derived from at least two donor individuals that each contained a mixture of DN and DP cells (Nawijn Nasal: multiciliated, Goblet; Regev/Rajagopal Lung: AT1, AT2, Basal, multiciliated, Secretory; Aggregated Lung: AT2, multiciliated, Secretory; Regev/Xavier Colon: *BEST4*^+^ Enterocytes, Cycling TA (Transit Amplifying), Enterocytes, Immature Enterocytes 2, TA-2). For each of these cell subsets, we then used MAST (version 1.8.2)^180^ to fit the following regression model to every gene with cells as observations:

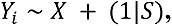

where *Y_i_* is the expression level of gene *i* in cells, measured in units of log2(TP10K+1), *X* is the binary co-expression state of each cell (*i.e.* DP *vs.* DN), and *S* is the donor that each cell was isolated from. To control for donor-specific effects (*i.e.* batch effects), we used a mixed model with a random intercept that varies for each donor. To fit this model, we subsampled cells from DP and DN groups to ensure that both the donor distribution and the cell complexity (*i.e.* the number of genes per cell) were evenly matched between the two groups, as follows. First, for each subset, we restricted our analysis to donors containing at least two DN and two DP cells. Using these samples, we partitioned the cells into 10 equally-sized bins based on cell complexity and subsampled DN cells from each bin to match the cell complexity distribution of the DP cells. Finally, we fit the mixed model (above), controlling for both donor and cell complexity.

To build gene modules for DP cells, we prioritized genes by requiring that they be expressed in at least 10% of DP cells, and to have a model coefficient greater than 0 with an FDR-adjusted p-value less than 0.05 (for the combined coefficient in the hurdle model). After this filtering step, genes were ranked by their model coefficient (*i.e.* estimated effect size). The top 12 genes were selected for network visualization within each cell type (**Fig. 4c,d, Extended Data Fig. 11a,b**). In three cases (gut Cycling TA, TA-2 and *BEST4*^+^ cells), *RP11-\** antisense genes were flagged and excluded from visualizations. To visualize overlap across each network, we indicated whether each gene was among the top 250 genes from each of the other cell types. Putative drug targets were identified by querying the Drugbank database^88^. Gene set enrichment analysis was performed using the R package EnrichR (version 1.0)^181^, selecting the top 25 genes from each cell type for the pan-tissue analysis (“All” category; **Fig. 4e**), and the top 50 genes from each cell type for the tissue-specific analyses (“Gut”, “Nasal”, and “Lung” categories; **Fig. 4e**). We note a few caveats/challenges/limitations that may influence our results, including non uniform sampling across donors; variation in cell compositions across regions (e.g., distal lung vs carina), and additional cellular heterogeneity that the current level of broad subset annotation may not have been captured.

### Cell-Cell interaction analysis

CellphoneDB^96^ v.2.0.0 was run with default parameters on the 10 human lung samples of the Regev/Rajagopal dataset, analyzing the cells from each dissected region separately. For each sample (patient/location combination), for each cell type we distinguished double positive cells (*ACE2* > 0 and *TMPRSS2* > 0) from all others. Only interactions highlighted as significant, i.e. present in the “significant means” output from CellphoneDB were considered.

### Co-expression patterns of additional proteases and *IL6*/*IL6R*/*IL6ST*

*ACE2*-protease co-expression (**Figure 6, Extended Data Fig. 14**) and *ACE2*-*IL6*/*IL6R*/*IL6ST* co-expression (**Extended Data Fig. 12**) were tested via the logistic mixed-effects model described in “Integrated co-expression analysis of high resolution cell annotations across tissues” (Equation 1, above).

### Code availability

Data and an interactive analysis examining the co-expression of genes across datasets can be accessed via the open-source data platform, Terra at https://app.terra.bio/#workspaces/kco-incubator/COVID-19_cross_tissue_analysis.

## Data availability

Interactive visualization and download of gene expression data can be accessed on the Single Cell Portal at https://singlecell.broadinstitute.org/single_cell?scpbr=hca-covid-19-integrated-analysis

## AUTHORS AND AFFILIATIONS

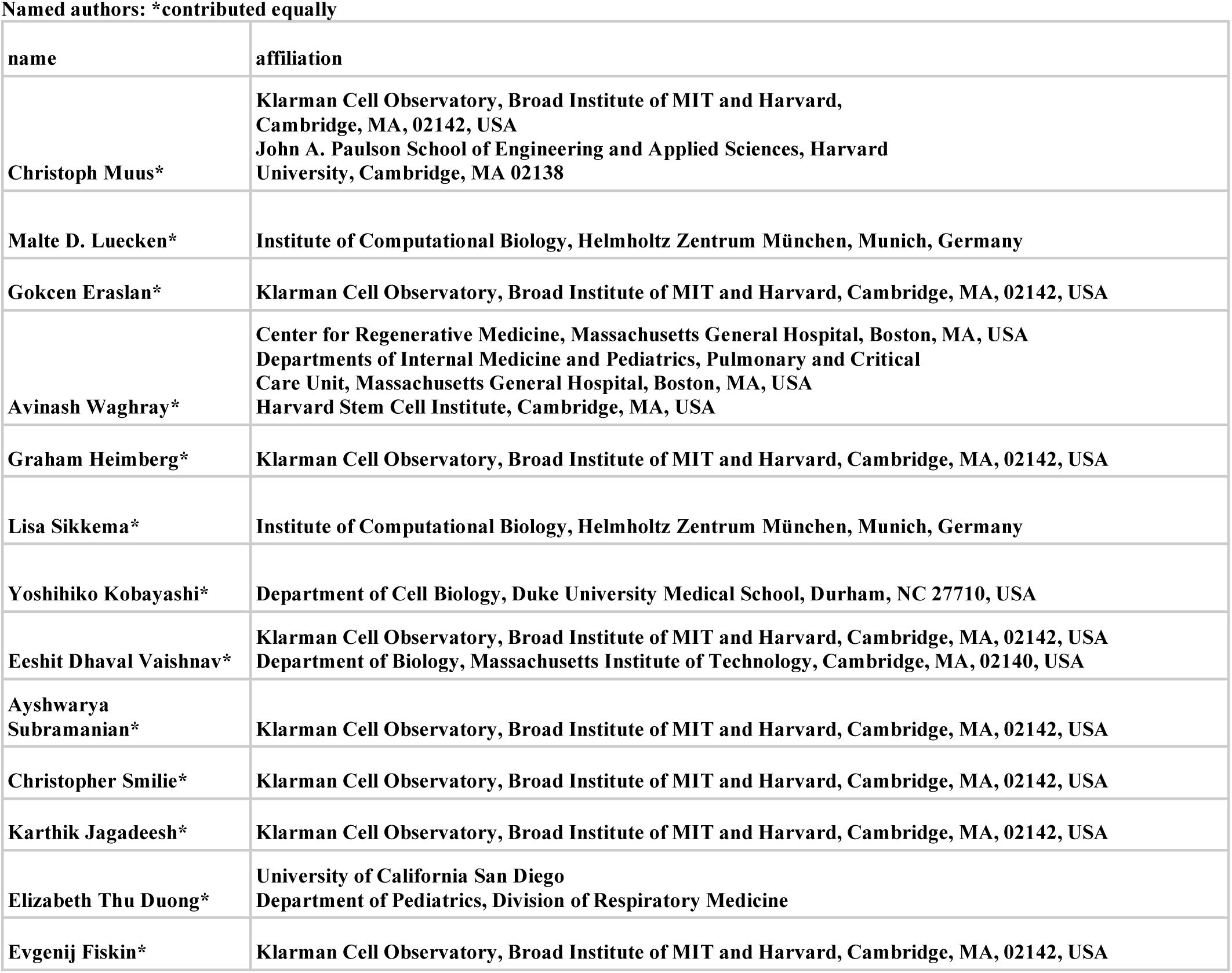
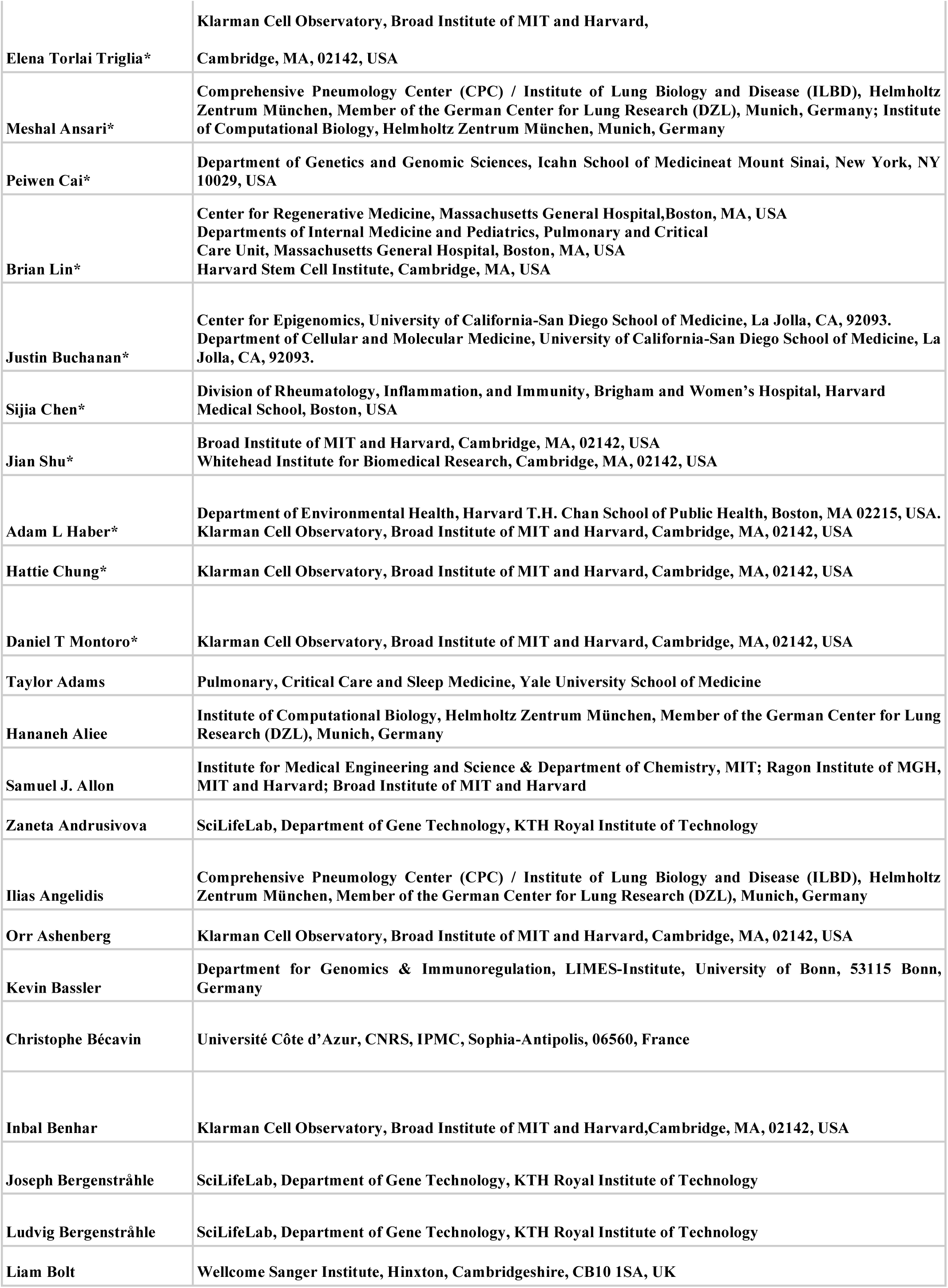
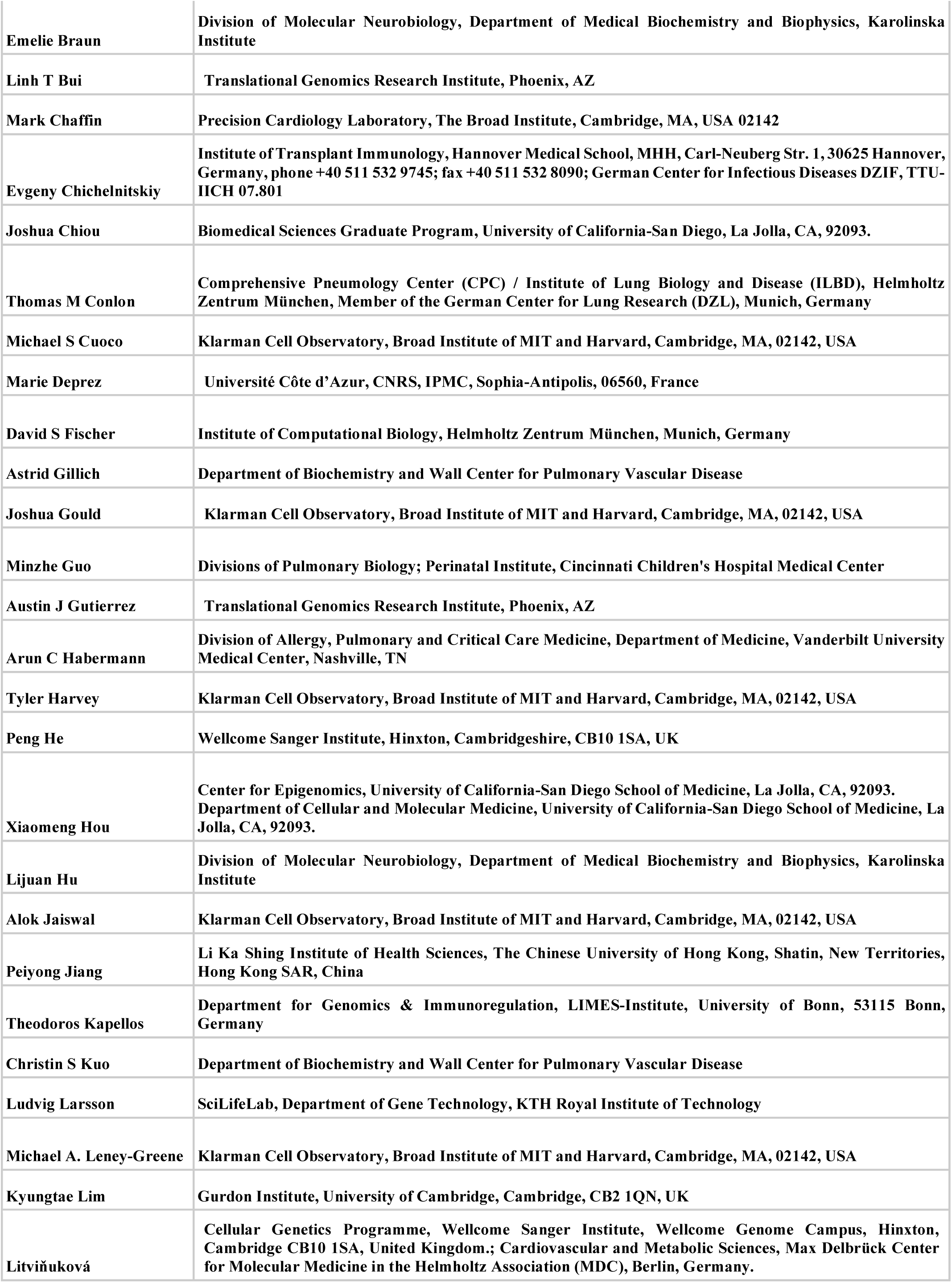
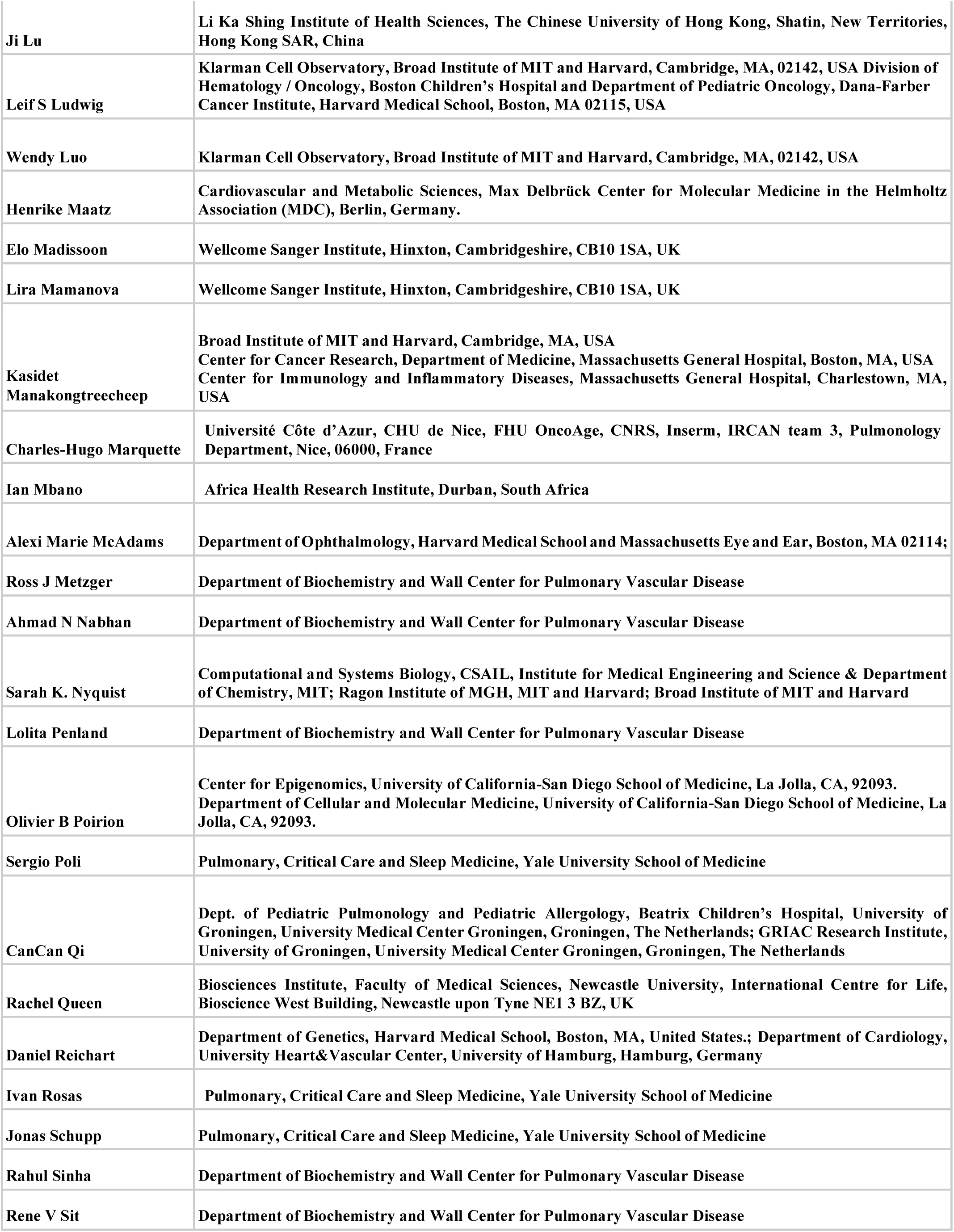
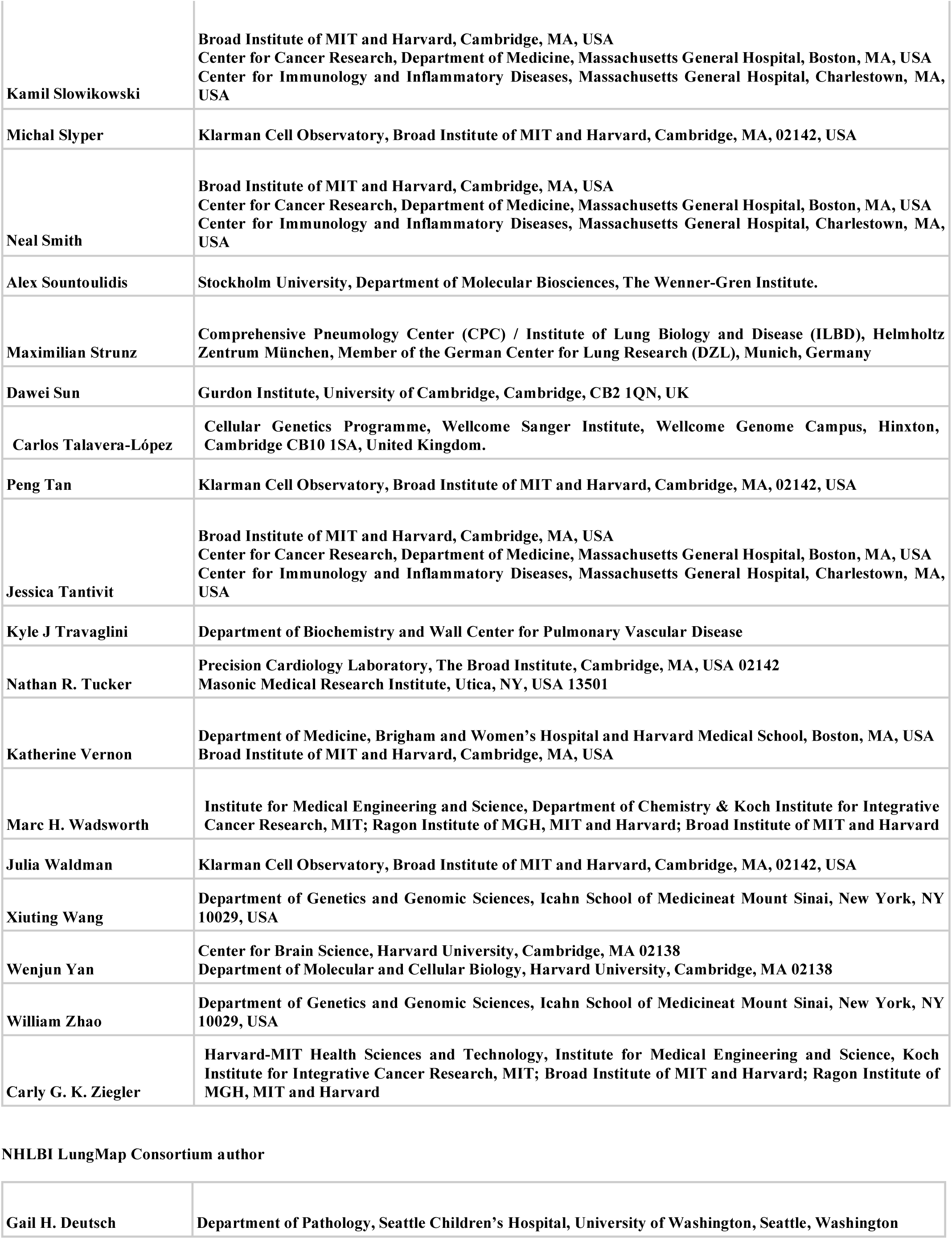
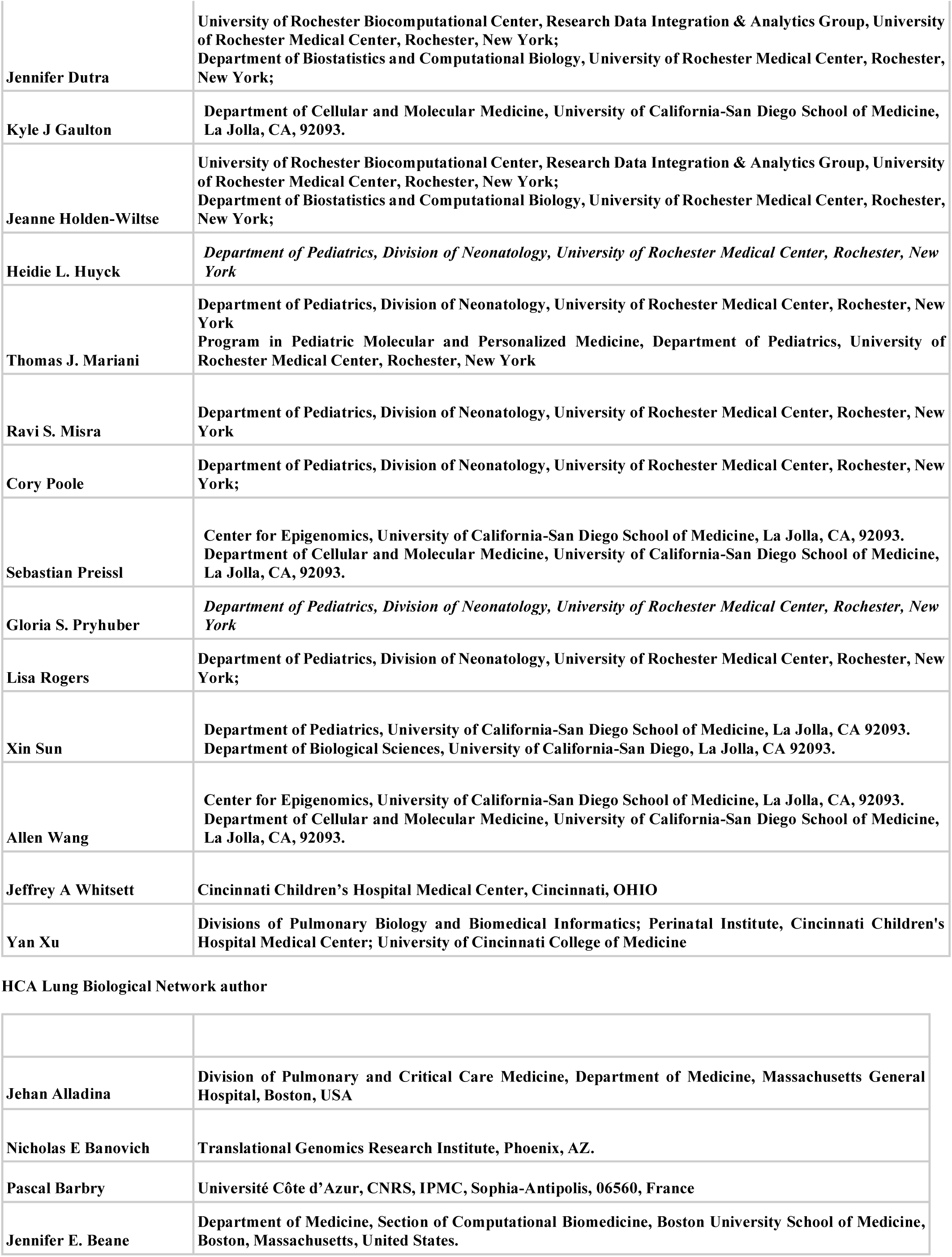
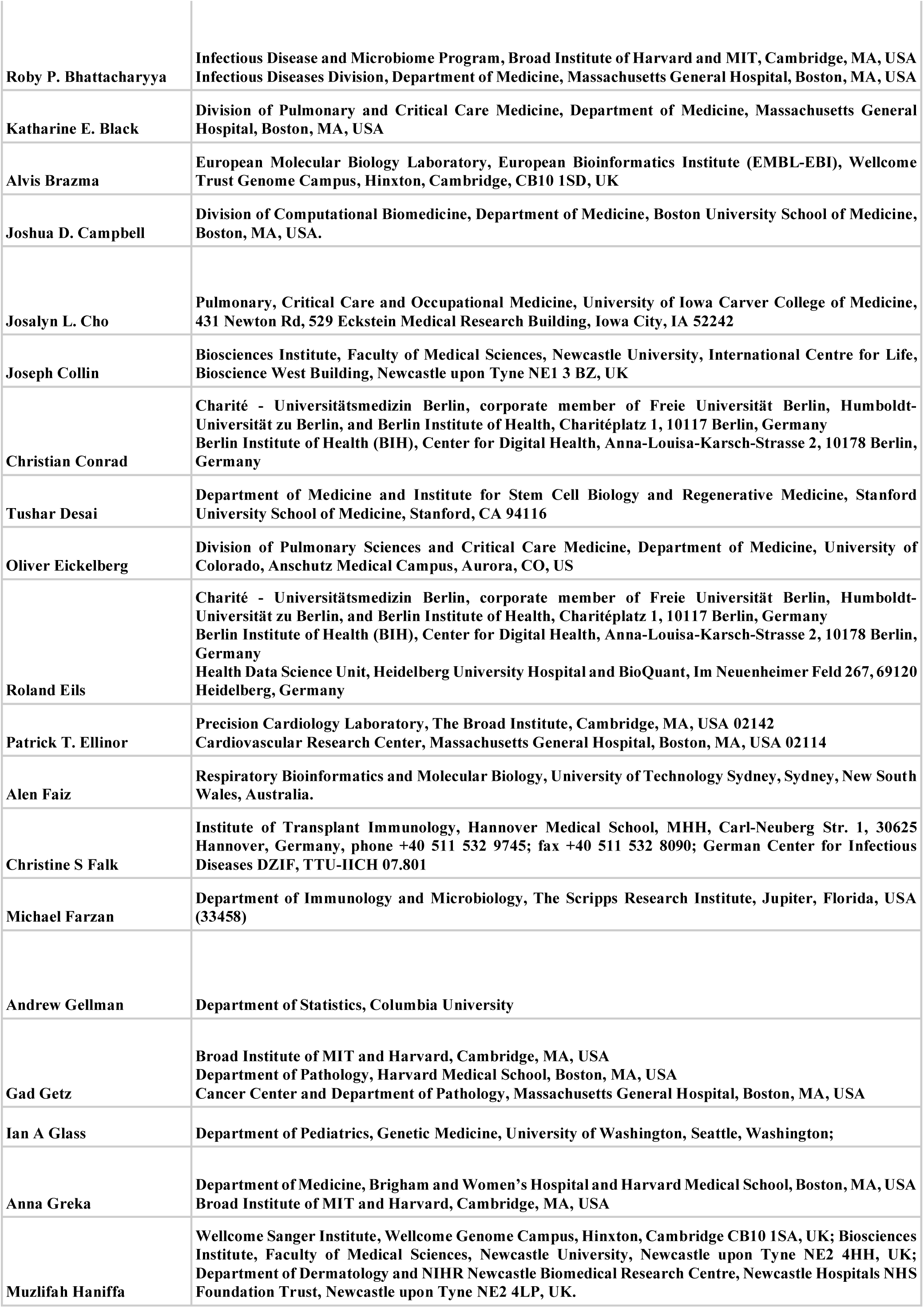
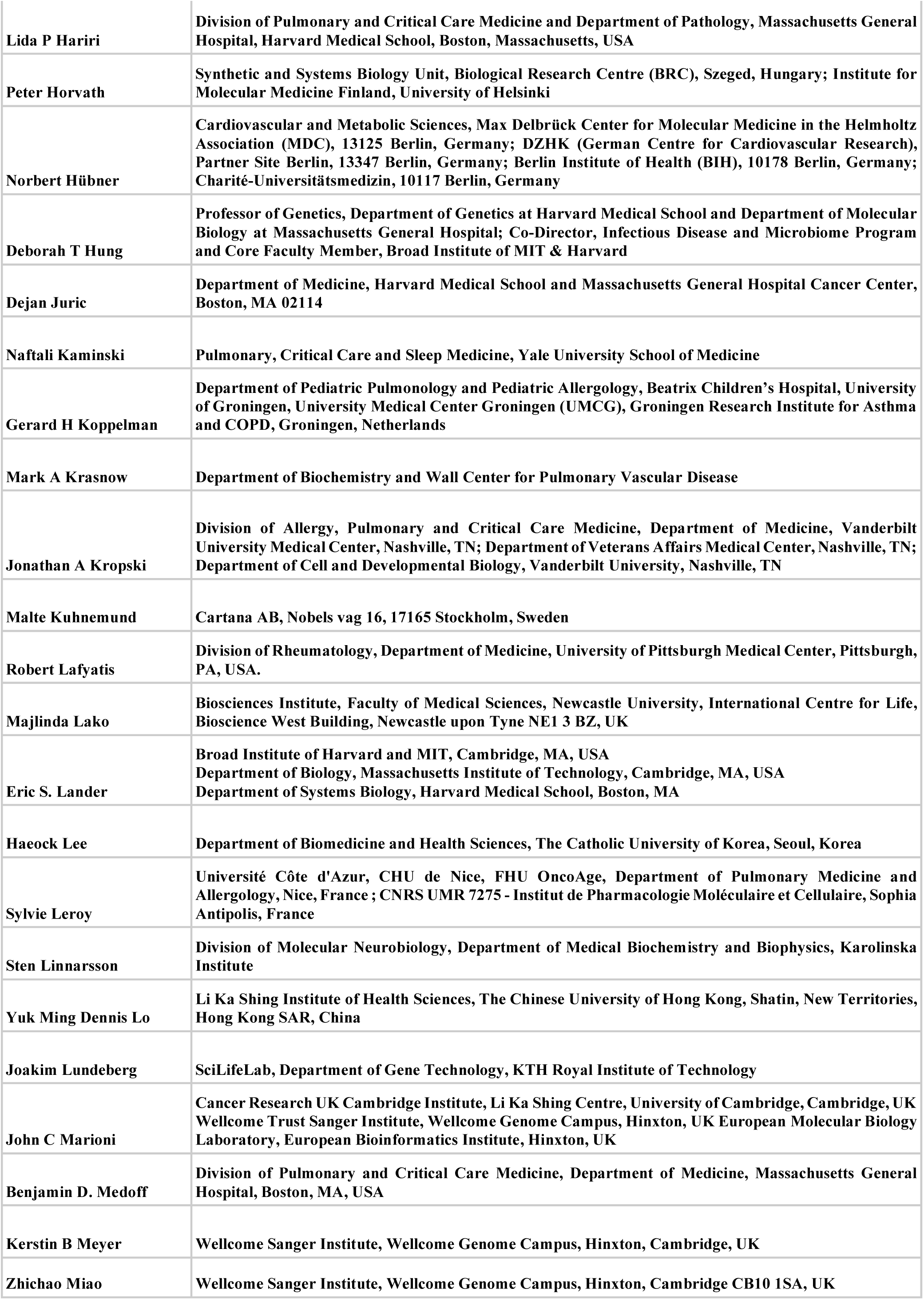
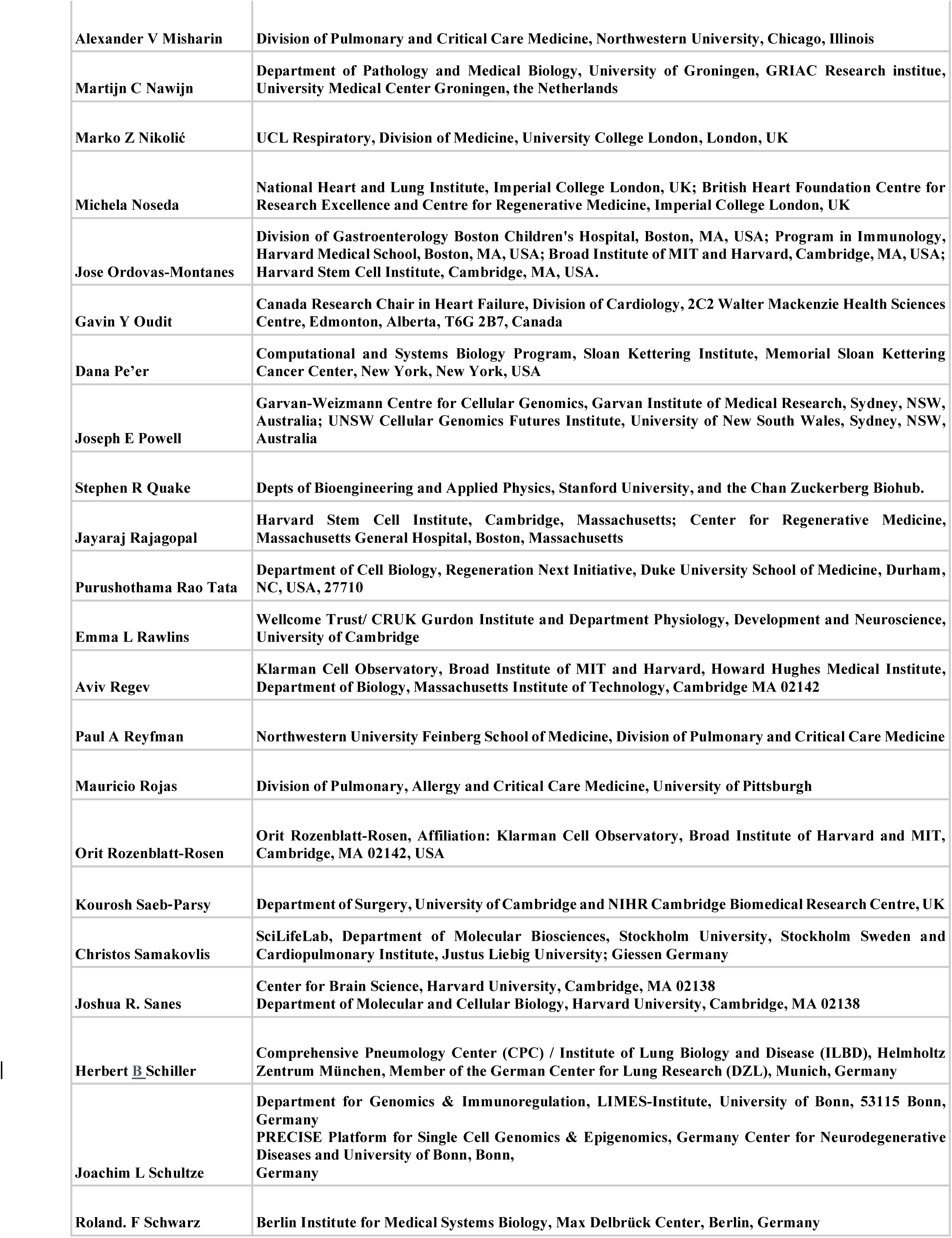
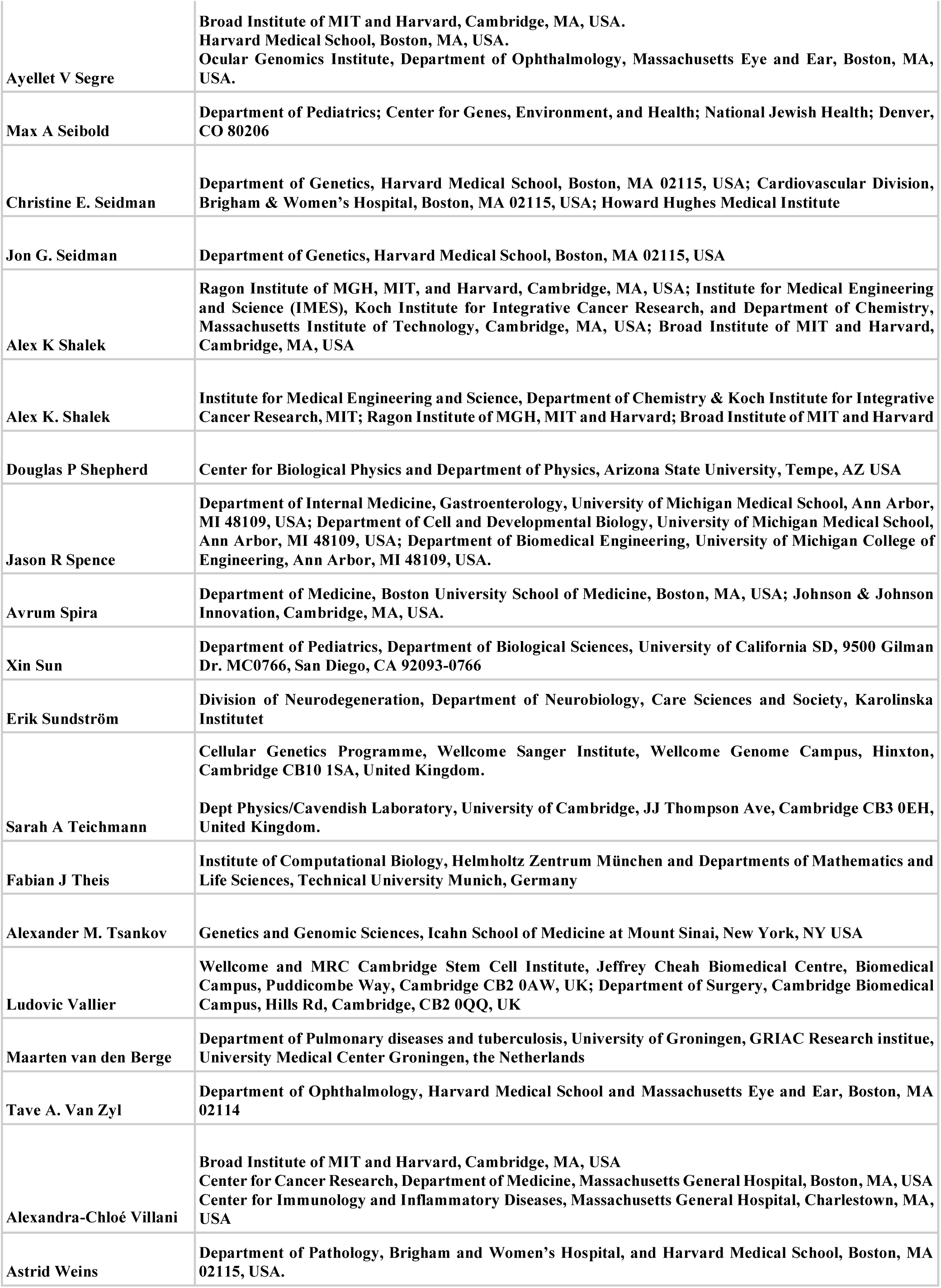
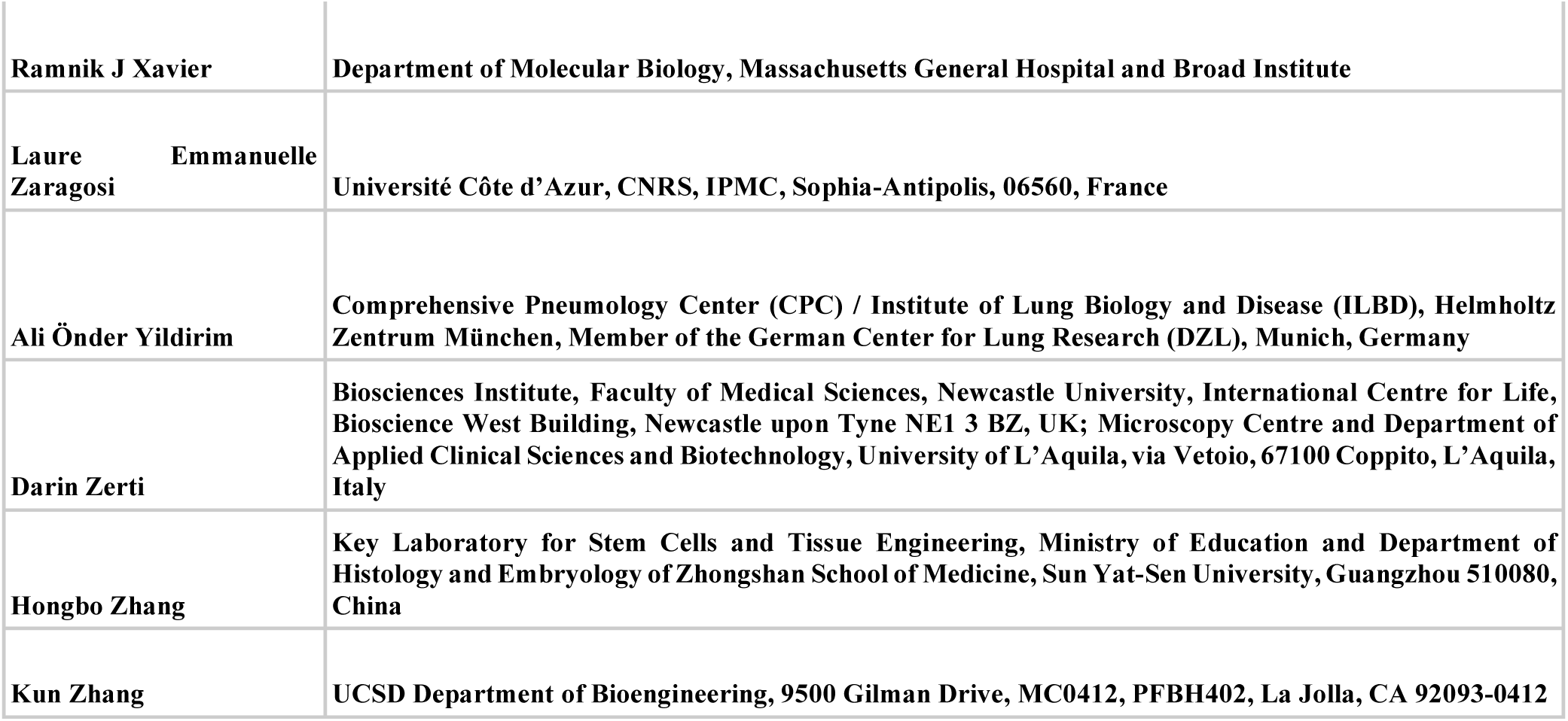

## Extended Data Figures

**Extended Data Figure 1.**
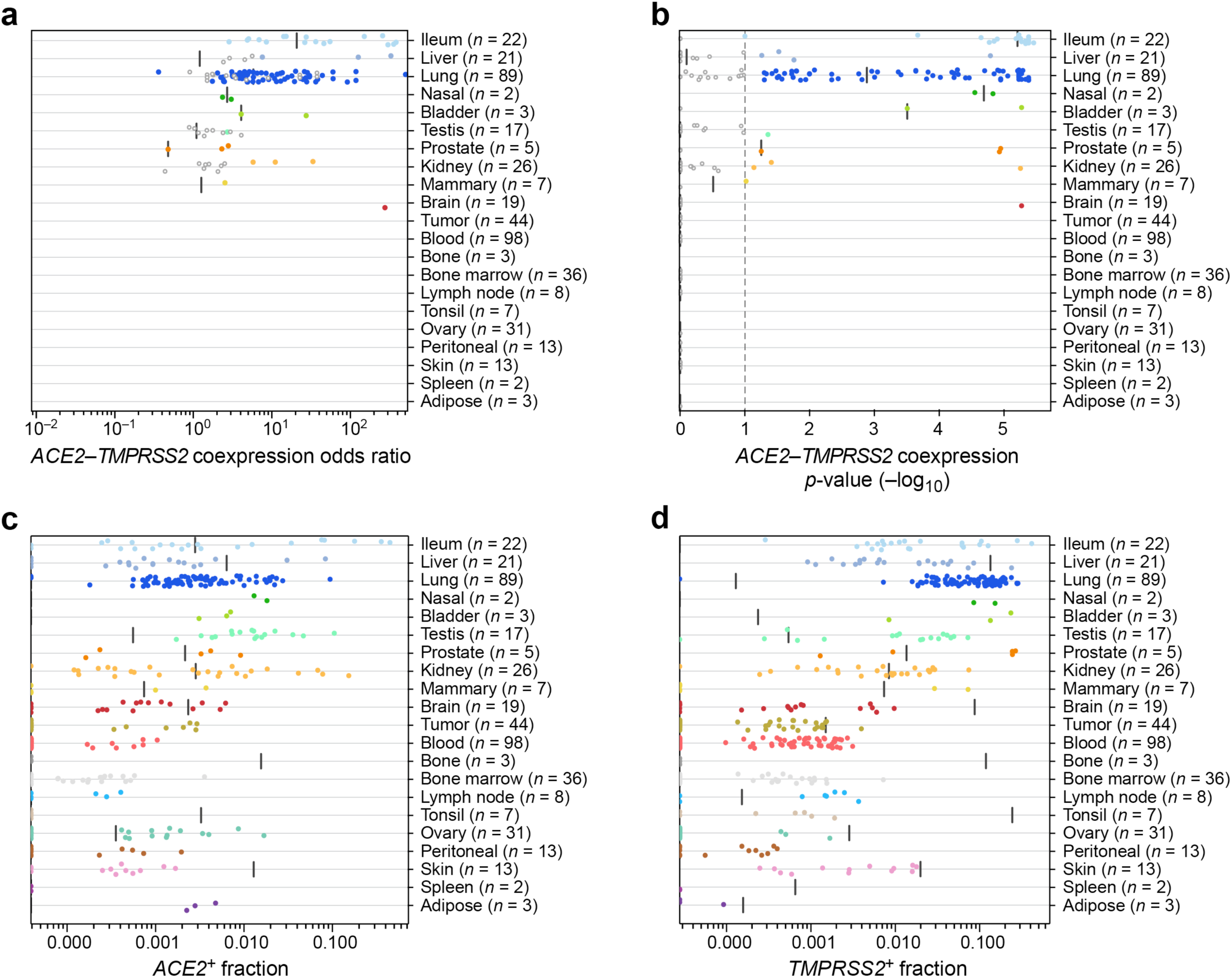
A cross-tissue survey of *ACE2^+^TMPRSS2^+^* cells in published single-cell datasets. **(a)** Odds ratio (x axis) of *ACE2^+^TMPRSS2^+^* co-expression in single-cell datasets (dots) from different tissues (y axis). (**b**) Significance (-log_10_(*p*-value), x axis) of co-expression of *ACE2^+^TMPRSS2^+^* in single-cell datasets (dots) from different tissues (y axis). (**c,d**) Proportion (x axis) of *ACE2^+^* cells per dataset (c) and *TMPRSS2^+^* cells per dataset (d) across different tissues (y axis).

**Extended Data Figure 2.**
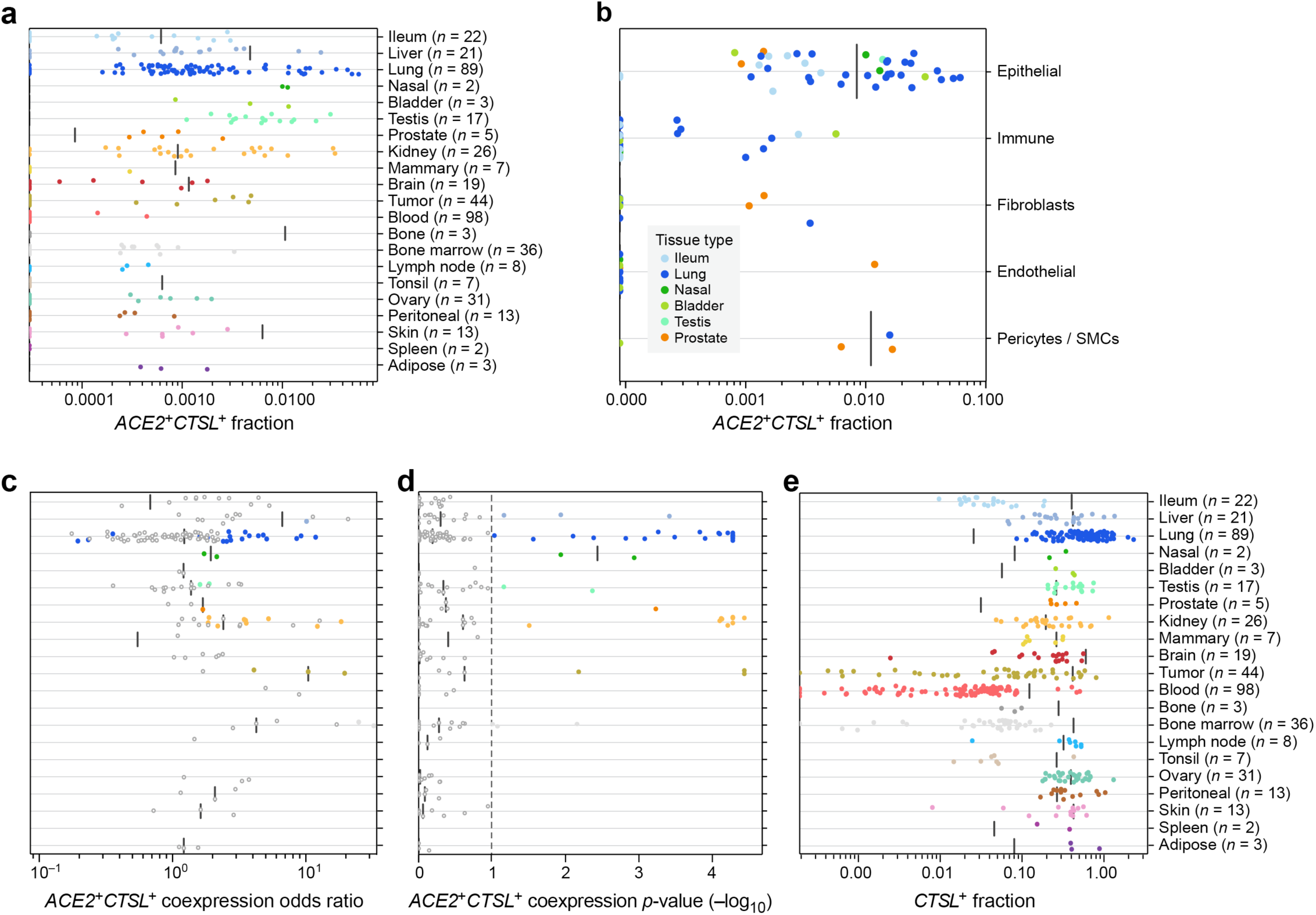
A cross-tissue survey of *ACE2^+^CTSL^+^* cells in published single-cell datasets. **(a)** Proportion (x axis) of *ACE2^+^CTSL^+^* cells per dataset (dots) across different tissues (y axis). (**b**) Proportion (x axis) of *ACE2^+^CTSL^+^* cells within clusters annotated by broad cell-type categories (dots) in each of the top 7 enriched datasets (y axis; color legend, inset). (**c**) Odds ratio (x axis) of *ACE2^+^CTSL^+^* co-expression in single-cell datasets (dots) from different tissues (y axis). (**d**) Significance (-log_10_(*p*-value), x axis) of co-expression of *ACE2* and *CTSL* in single-cell datasets (dots) from different tissues (y axis). (**e**) Proportion (x axis) of *CTSL^+^* cells per dataset across different tissues (y axis).

**Extended Data Figure 3.**
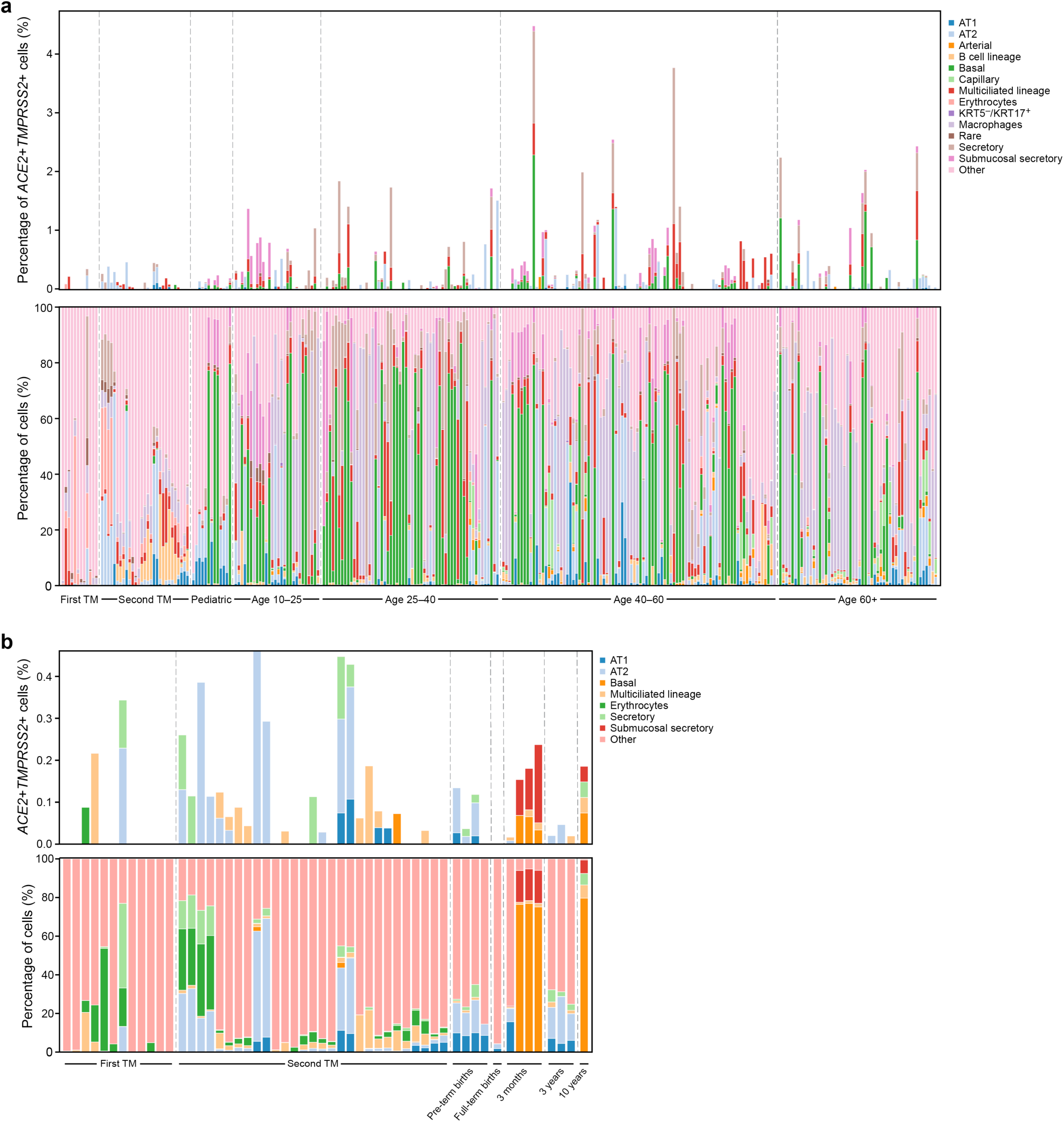
Cellular composition and fraction of *ACE2^+^TMPRSS2^+^* cells across the integrated lung dataset **(a)** Percentage of *ACE2*^+^*TMPRSS2*^+^ (double positive - DP) cells across samples with sample composition. Top: Percentage ACE2^+^TMPRSS2^+^ cells in each sample, from each level 2 cell type in which DP cells are found. Bottom: Sample composition. All level 2 cell type annotations where no DP cells are found are summarized as “Other” for ease of visualization. Samples are ordered by age with 31-week pre-term births and 39-week full-term births both set to age 0. (**b**) Zoom in on fetal and pediatric samples of plot (a). All level 2 cell types without DP cells in fetal and pediatric samples are labeled as “Other”. Samples are ordered and labeled by age. Fetal samples are partitioned into first and second trimester (TM) and pediatric samples are divided into 31-week pre-term births, 39-week full term births, 3 month, 3 year, and 10 year old children.

**Extended Data Figure 4.**
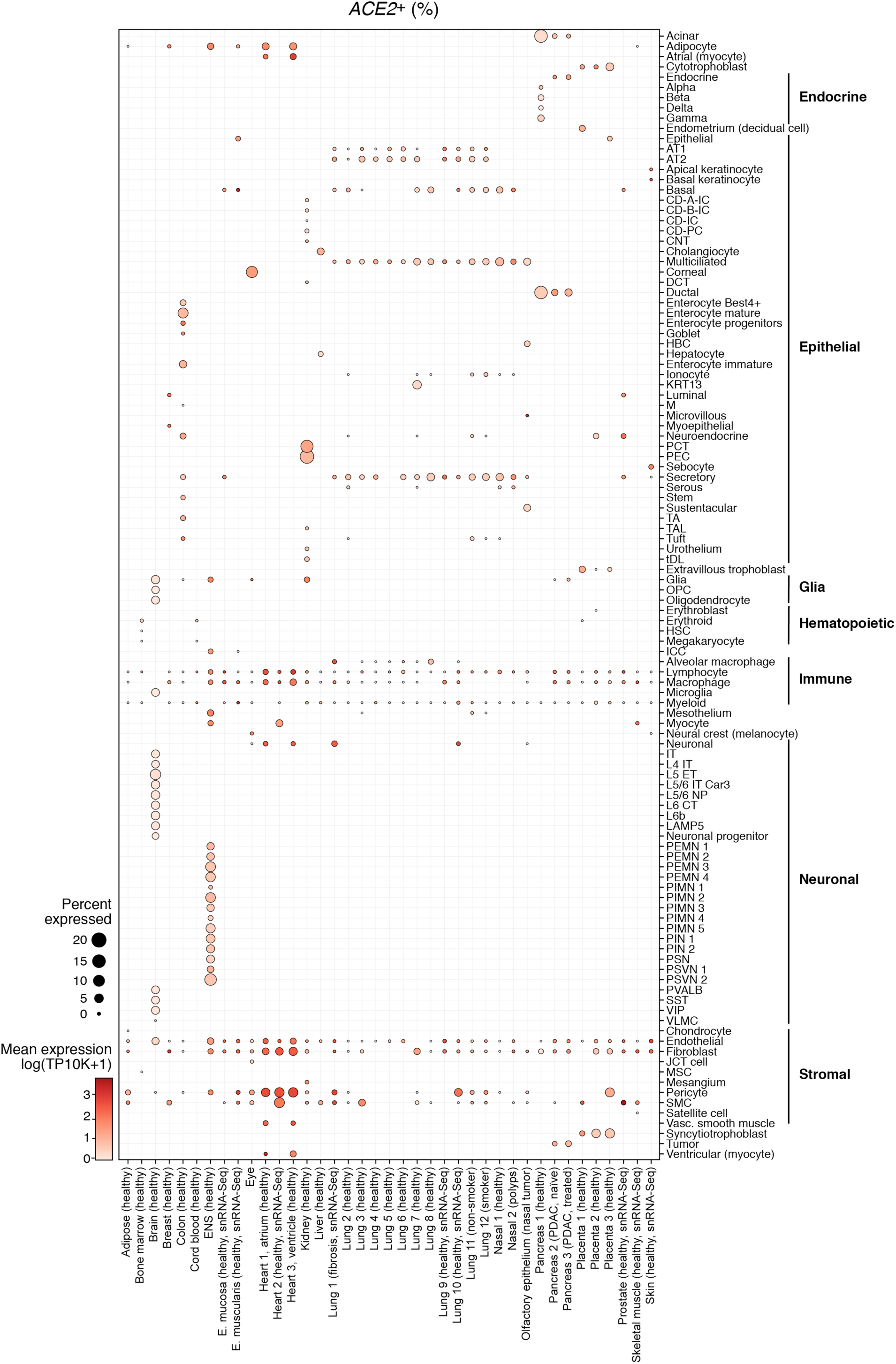
*ACE2* expression across tissues and celltypes. Shown are fractions of *ACE2* expressing cells (dot size) and mean ACE2 expression level in expressing cells (dot color) across datasets (rows) and cell types (columns).

**Extended Data Figure 5.**
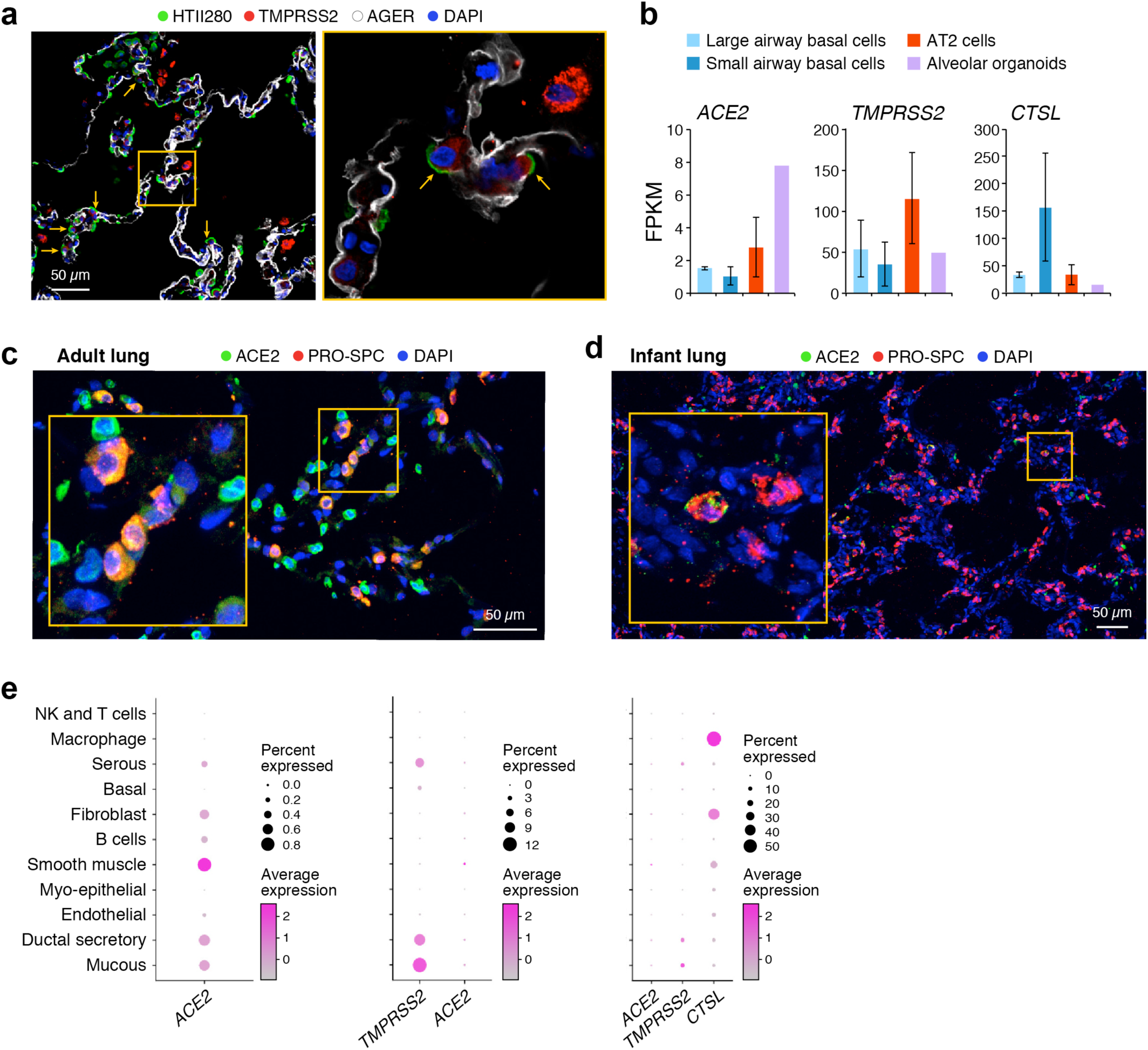
ACE2, TMPRSS2, CTSL Immunofluorescence and RNA profiling **(a)** Immunostaining in human adult lung alveoli. TMPRSS2 (red), HTII-280 (green) and AGER (white). Blue shows DAPI in nuclei. (**b**) Mean expression (y axis, FPKM, from bulk RNA-seq) of *ACE2, CTSL, TMPRSS2* in sorted cells from 3 different human explant donors using the following markers: large and small airway basal cells (NGFR+), AT2 cells (HT-II 280+) and alveolar organoids (HT-II 280+). (**c**) Human adult lung: alveolar section stained with ACE2 (green) and Pro-SFTPC (red) antibodies and DAPI (blue). (**d**) Lung from infant donor: alveolar section stained with ACE2 (green) and Pro-SFTPC (red) antibodies and DAPI (blue). (**e**) Expression in the submucosal gland. Mean expression (color) and proportion of expressing cells (dot size) of *ACE2*, *TMPRSS2* and *CTSL* in key cell types (rows), from scRNA-seq of human large airway submucosal glands.

**Extended Data Figure 6.**
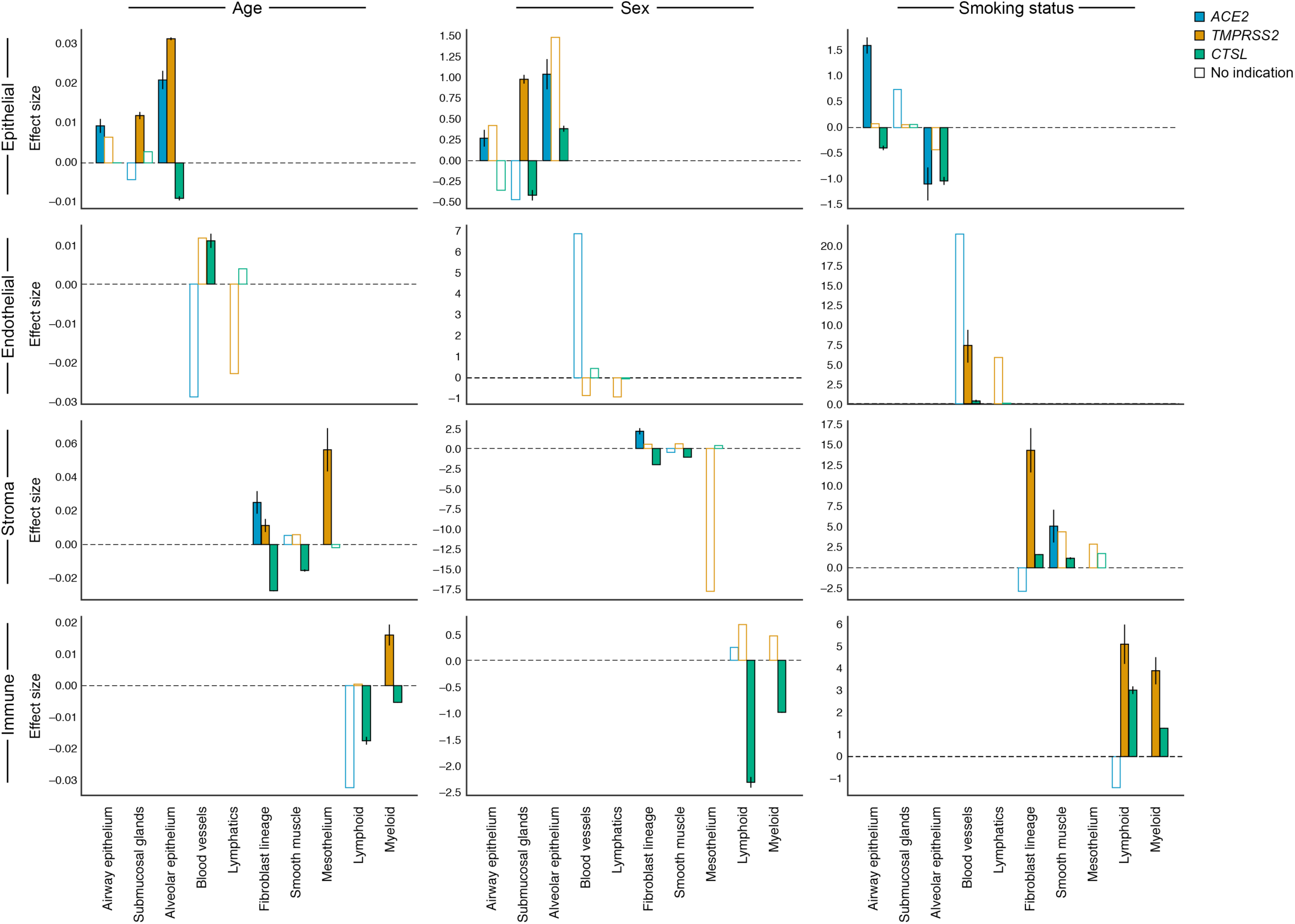
Age, sex, and smoking status associations with expression of *ACE2*, *TMPRSS2*, and *CTSL* across level 2 cell type annotations. Effect size (y axis) of association as log fold changes (sex, smoking status) and slope of log expression with age. Bars that are colored in indicate associations with an FDR-corrected p-value of < 0.05 where the pseudo-bulk analysis shows a consistent effect direction. Error bars represent model uncertainties.

**Extended Data Figure 7.**
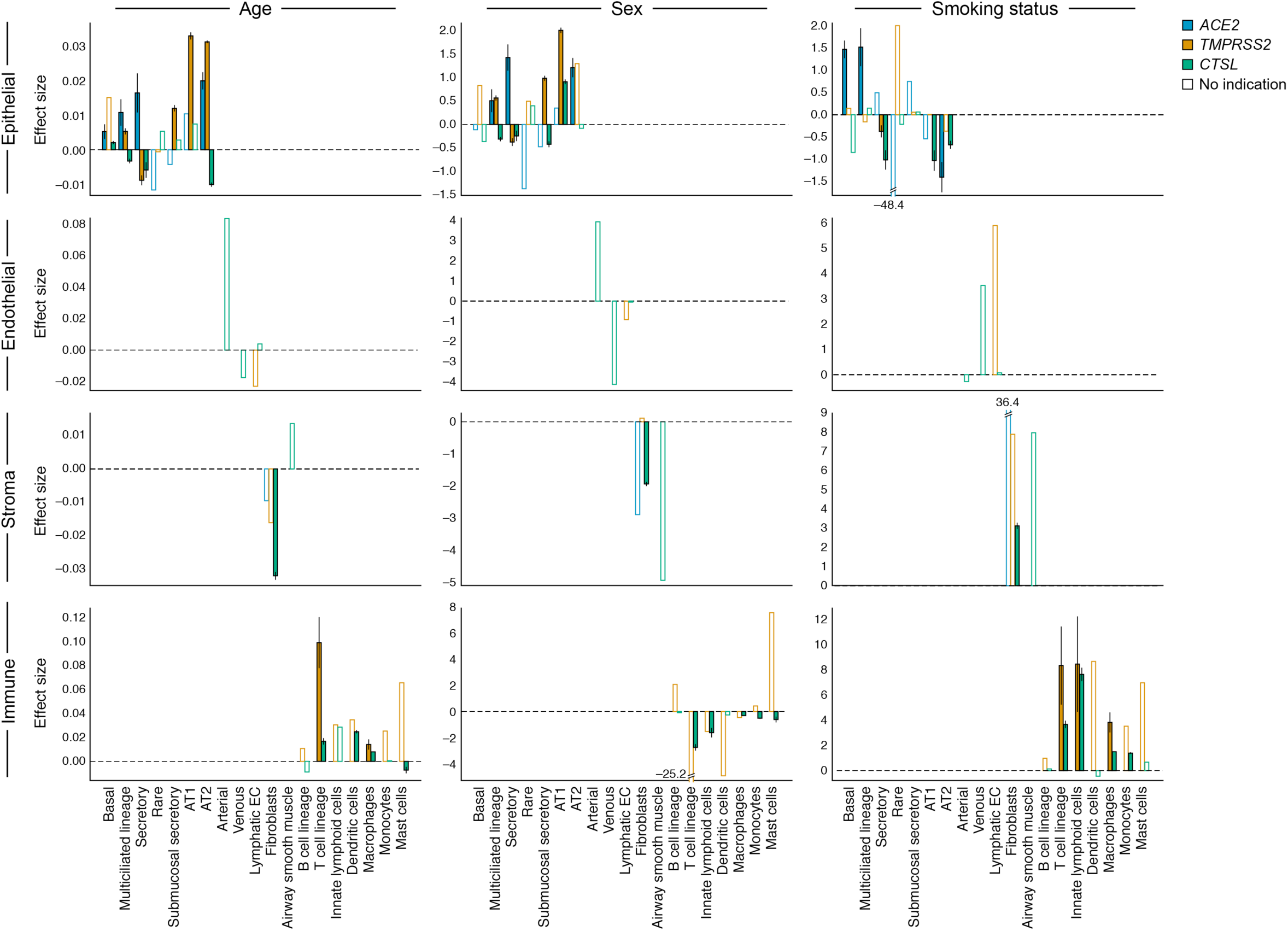
Age, sex, and smoking status associations with expression of *ACE2*, *TMPRSS2*, and *CTSL* across level 3 cell type annotations. Effect size (y axis) of the association as log fold changes (sex, smoking status) and slope of log expression with age. Bars that are colored in indicate associations with an FDR-corrected p-value of < 0.05 where the pseudo-bulk analysis shows a consistent effect direction. Error bars represent model uncertainties.

**Extended Data Figure 8.**
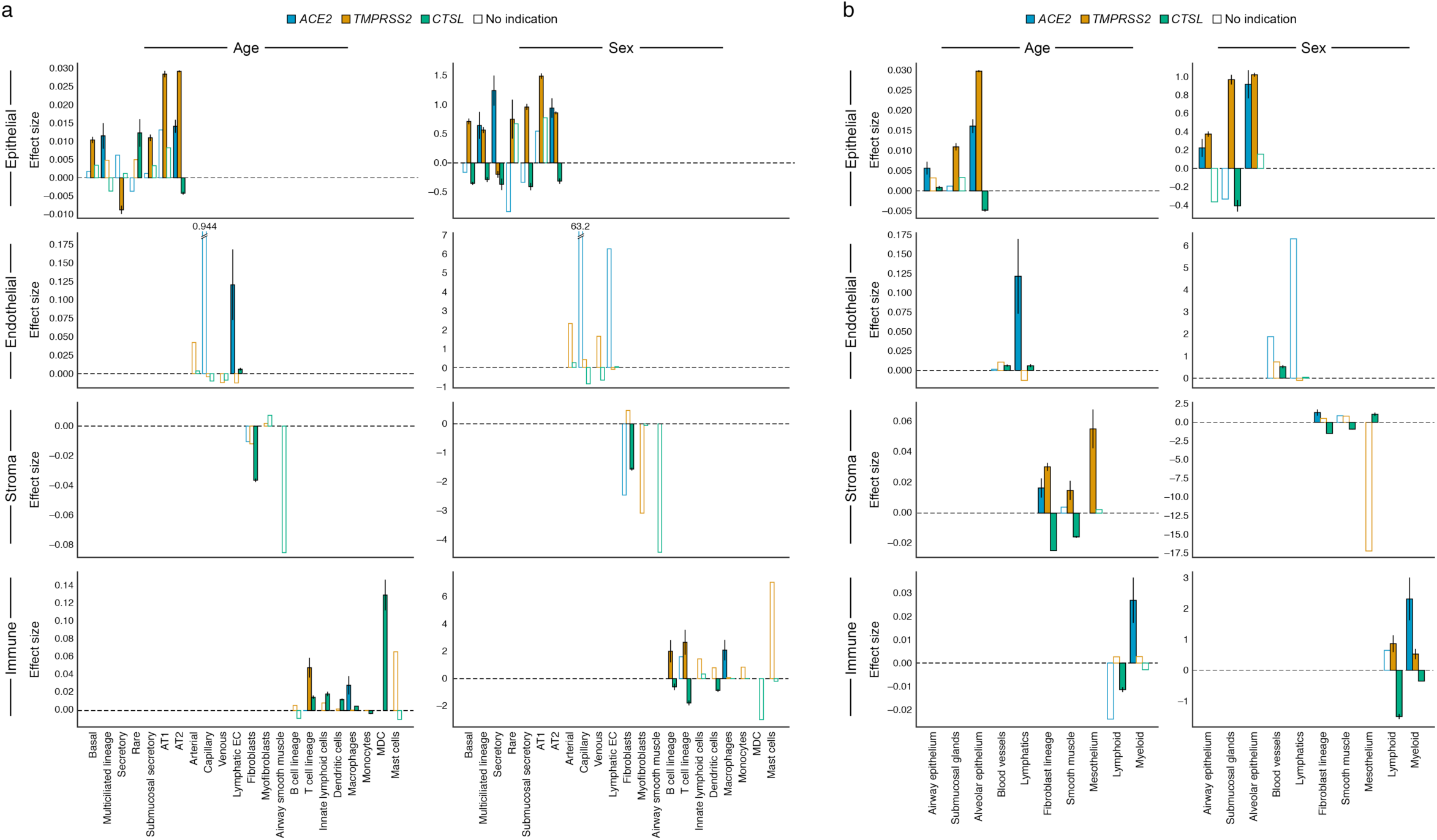
Age and sex associations with expression of *ACE2*, *TMPRSS2*, and ***CTSL* in level 3 and level 2 cell type annotations on the full, non-fetal, lung data** Effect size (y axis) of the association as log fold changes (sex, smoking status) and slope of log expression with age. Bars that are colored in indicate associations with an FDR-corrected p-value of < 0.05 where the pseudo-bulk analysis shows a consistent effect direction. Error bars represent model uncertainties. (**a**) Level 3 and (**b**) Level 2 cell type annotations.

**Extended Data Figure 9.**
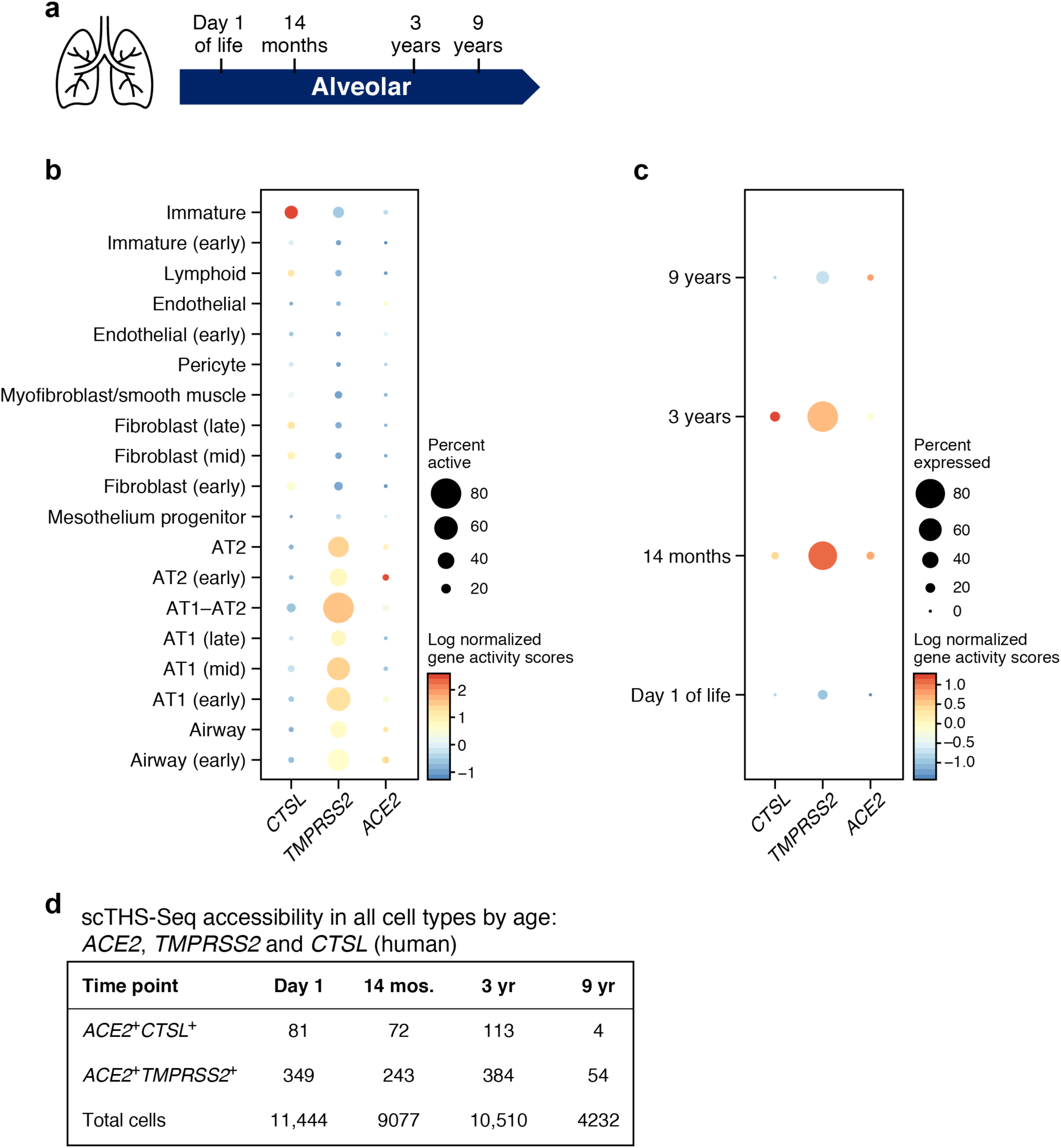
Chromatin accessibility at the *ACE2, TMPRSS* and *CTSL* loci across lung cells in early life **(a)** Schematic: single-cell chromatin accessibility by transposome hypersensitive sites sequencing (THS-Seq) from human pediatric samples (full gestation, no known lung disease) collected at day 1 of life, 14 months, 3 years, and 9 years (n=1 at each time point). (**b**) Accessibility (dot color log normalized gene activity scores), and % of cells with accessible loci (dot size) for the *ACE2, TMPRSS,* and *CTSL* loci (columns) across different cell types (rows) in scTHS-Seq with all time points aggregated. (**c**) Accessibility (dot color log normalized gene activity scores), and % of cells with accessible loci (dot size) of *ACE2, TMPRSS* and *CTSL* in AT1--AT2 cells in scTHS-Seq at day 1 of life, 14 months, 3 years, and 9 years (rows). (**d**) Number of *ACE2*+*CTSL*+ and *ACE2*+*TMPRSS2*+ cells per time point.

**Extended Data Fig. 10.**
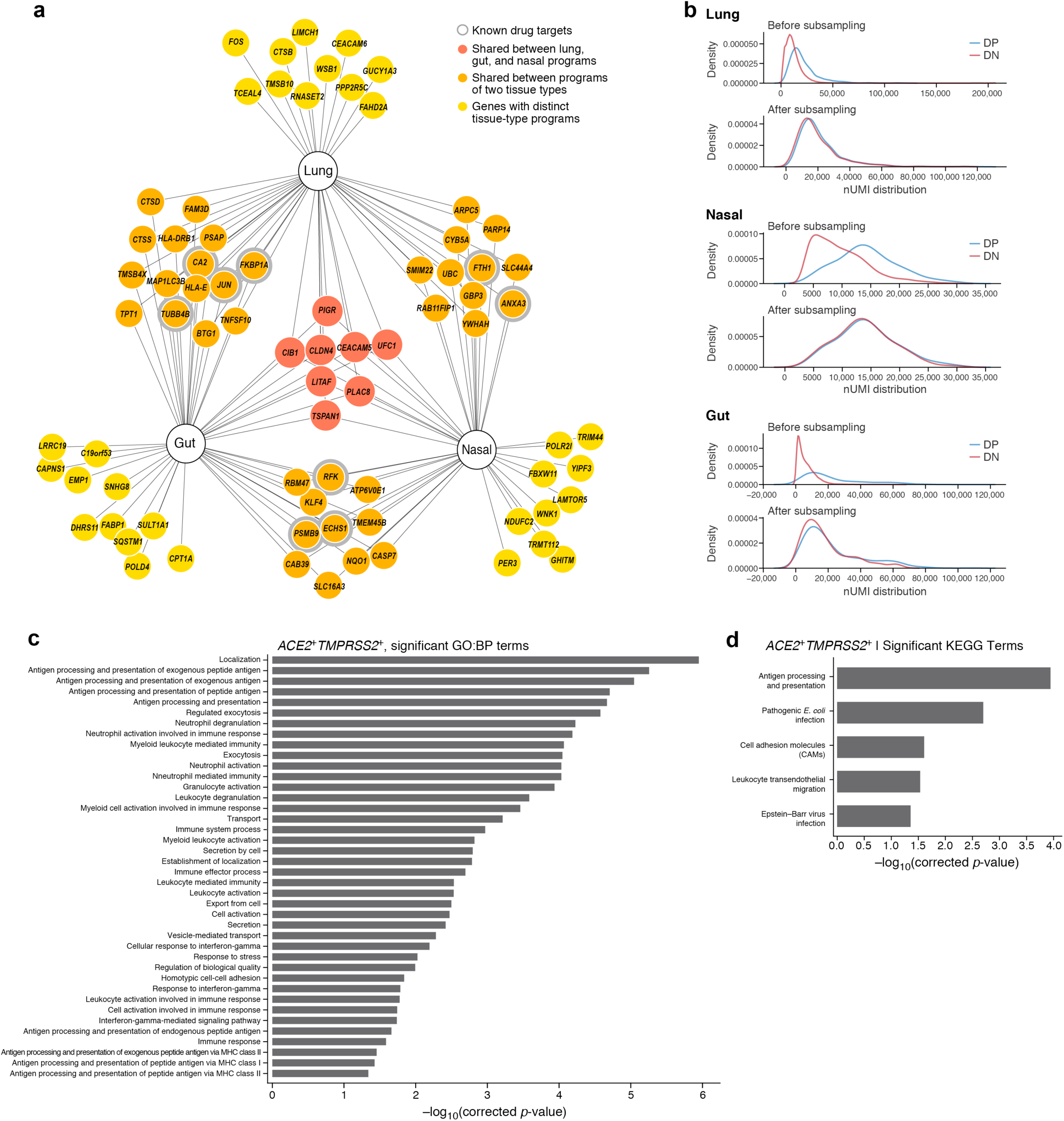
Tissue programs for dual positive cells **(a)** Selected tissue program genes. Node: gene; Edge: program membership. Genes are selected heuristically for visualization, derived from the positive feature importance values of a random forest classifier without nUMI distribution matching (Methods).). (**b**) Stratified subsampling to match nUMI distributions. (**c,d**) Enrichment (-log10(adj P-value), x axis) of GO Biological Process (c) and KEGG pathway (d) gene sets (y axis) in the full tissue programs without nUMI distribution matching.

**Extended Data Fig. 11.**
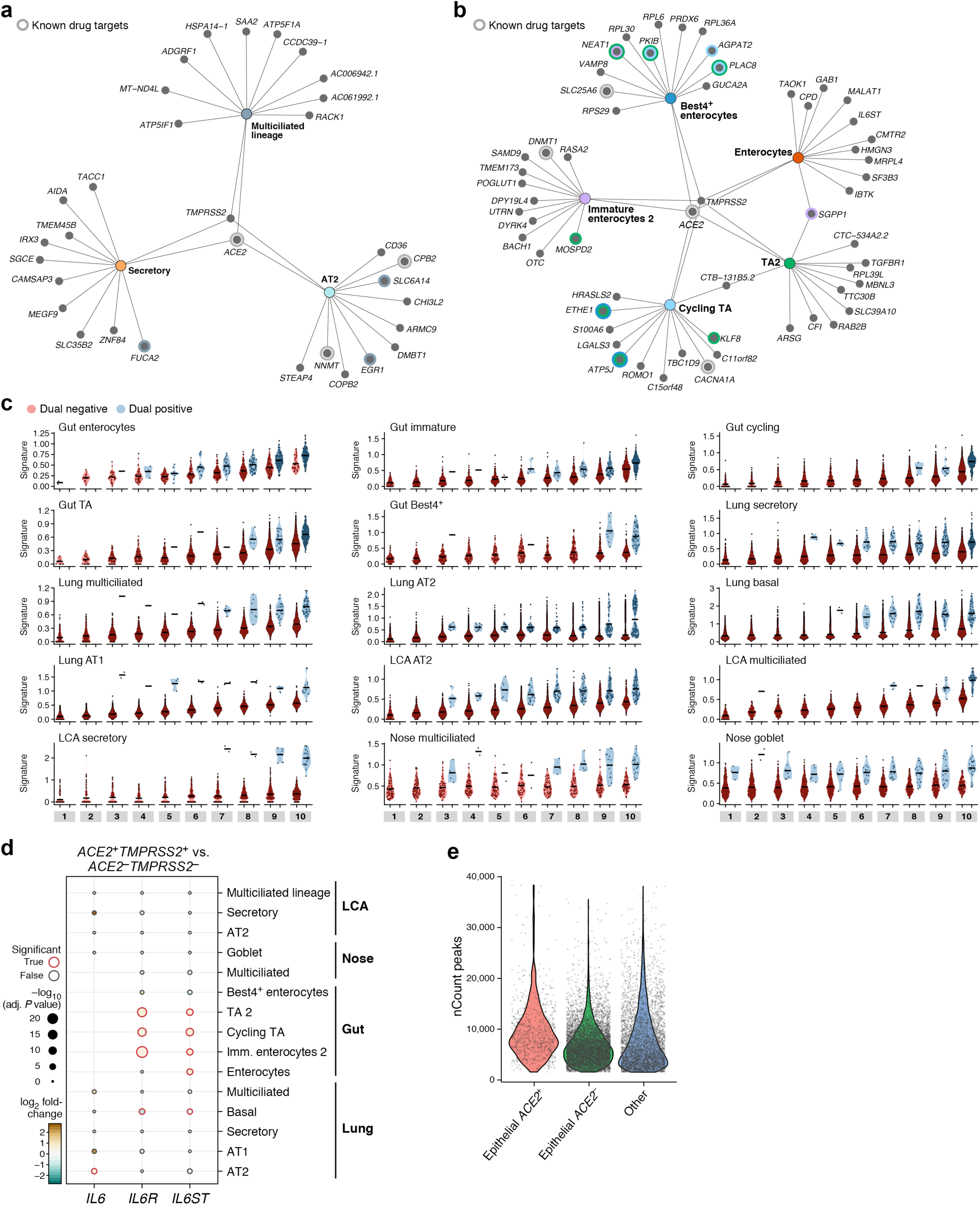
Cell programs for dual positive cells (**a,b**) Top 12 genes from each cell program recovered for different lung (**a**) or gut (**b**) epithelial cell-type (nodes, colors). Colored concentric circles: overlap with a gene in the top 250 significant genes in other cell types. ACE2 and TMPRSS2 are included even if not among the top 12. (**c**) Comparison of signature scores of cell programs between DP and DN cells for each cell type stratified by gene complexity bin. Cells were partitioned into 10 gene complexity bins for every cell type. (**d,e**) IL6 and its receptor’s expression in specific cell types in lung and heart. (**d**) Significance (dot size) and fold change (dot color) of differential expression between DP and DN cells within different types (rows) for *IL6* and its receptors *IL6R* and *IL6ST* (columns) across tissues. (**e**) Top: Significance (dot size) and fold change (dot color) of differential expression between DP and DN cells within different cell types in the heart (rows)for *IL6* and its receptors *IL6R* and *IL6ST* (columns). Bottom: Significance (dot size) and effect size (dot color) from a mixed effects model of co-expression of IL6, IL6R, or IL6ST (columns) coexpression with *ACE2*. (f) Distribution of number of counts in peaks (y axis) in *ACE2^+^* epithelial cells (having at least 1 fragment in the *ACE2* gene locus) and *ACE2^-^* cells.

**Extended Data Figure 12.**
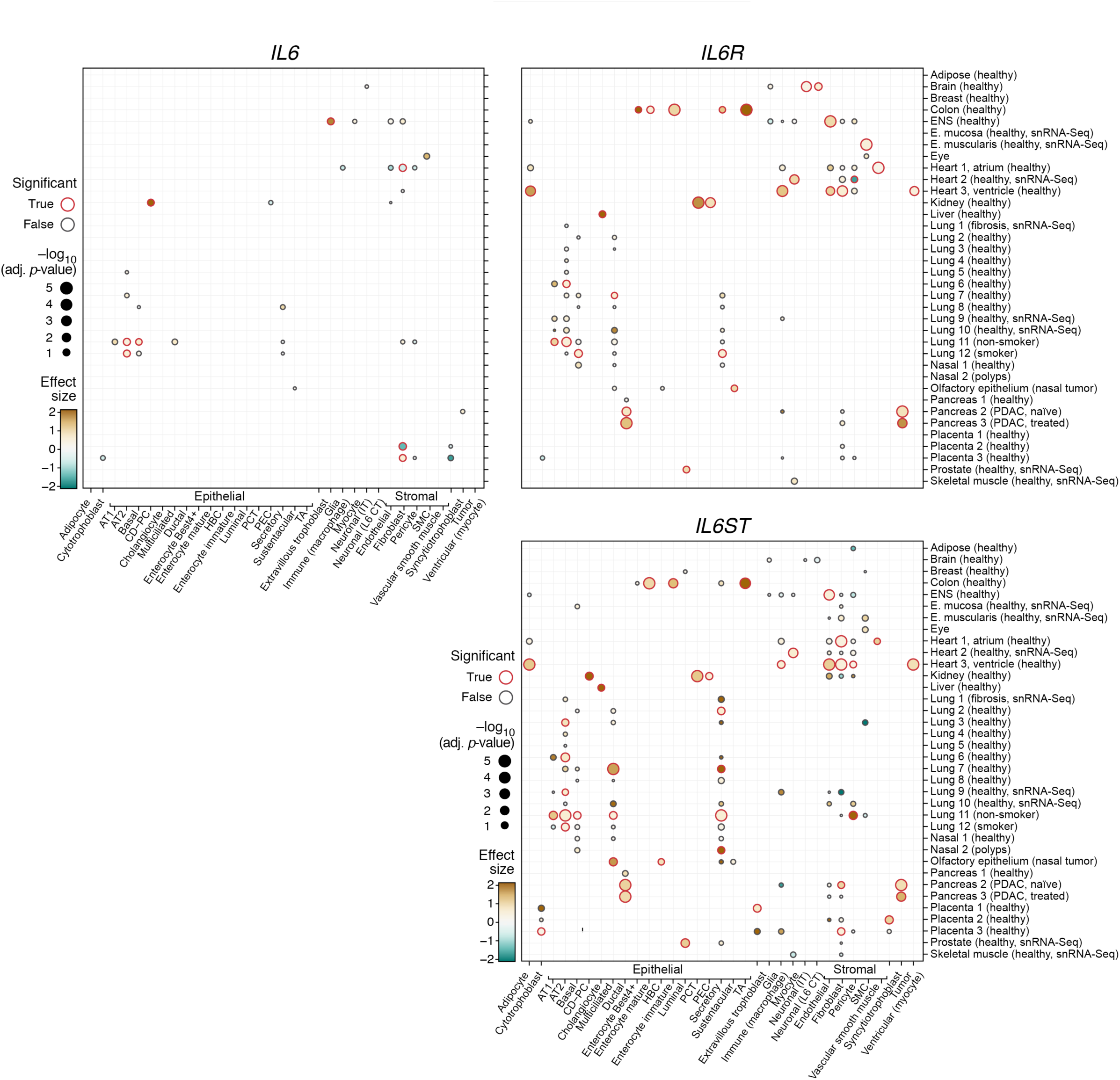
Co-expression of *ACE2* and *IL6*,*IL6R*,*IL6ST*. Co-expression of ACE2 and IL6,IL6R,IL6ST in select single-cell datasets. P-values and significance (FDR 10%) derived from the logistic mixed-effects model.

**Extended Data Figure 13.**
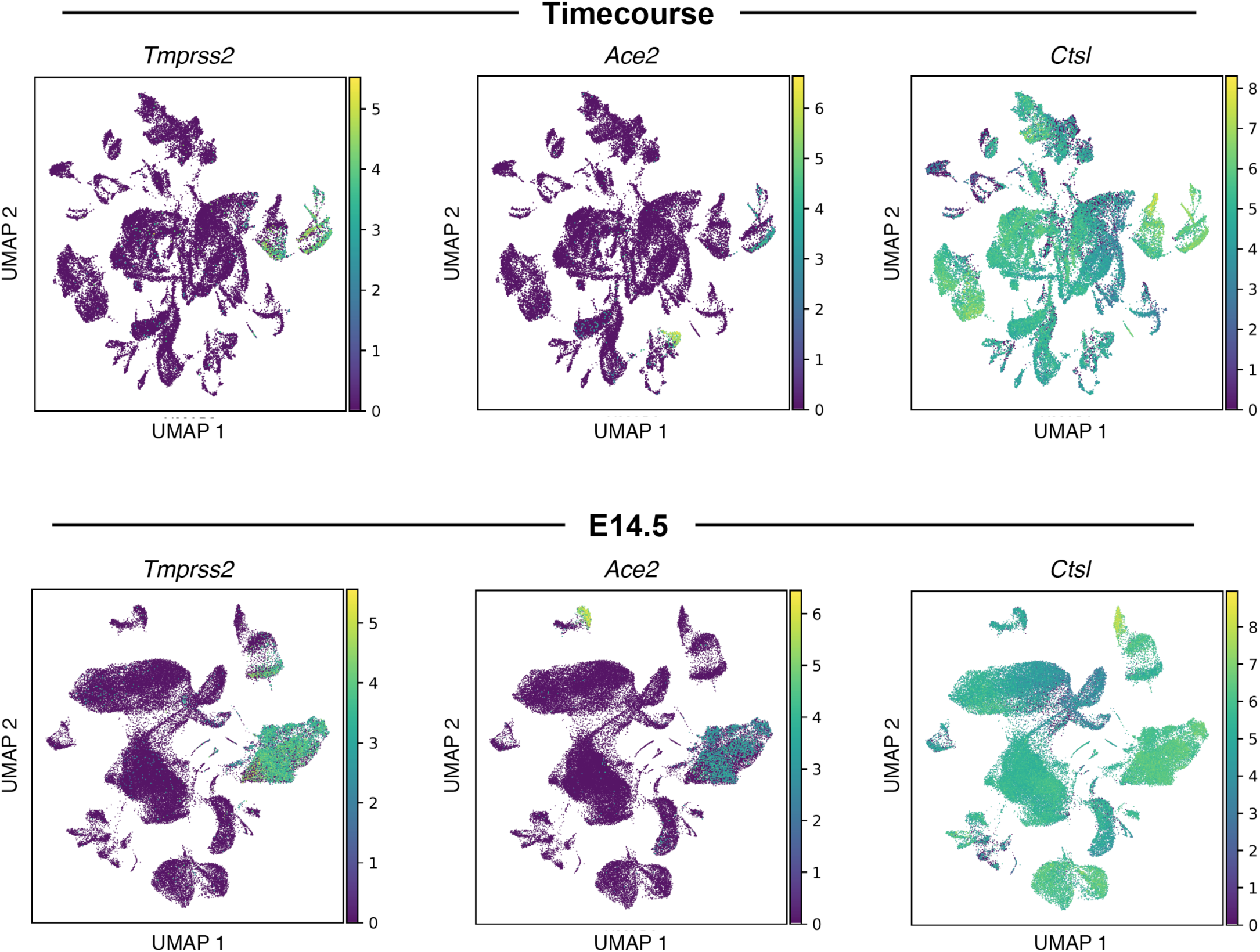
Expression of *Ace2*, *Tmprss2* and *Ctsl* in mouse placenta. UMAP embedding (as in Fig. 5k) of scRNA-seq profiles of placenta cells collected at E.14.5 (top) or along a time course (bottom), colored by expression level of*Ace2*, *Tmprss2*, and *Ctsl*.

**Extended Data Figure 14.**
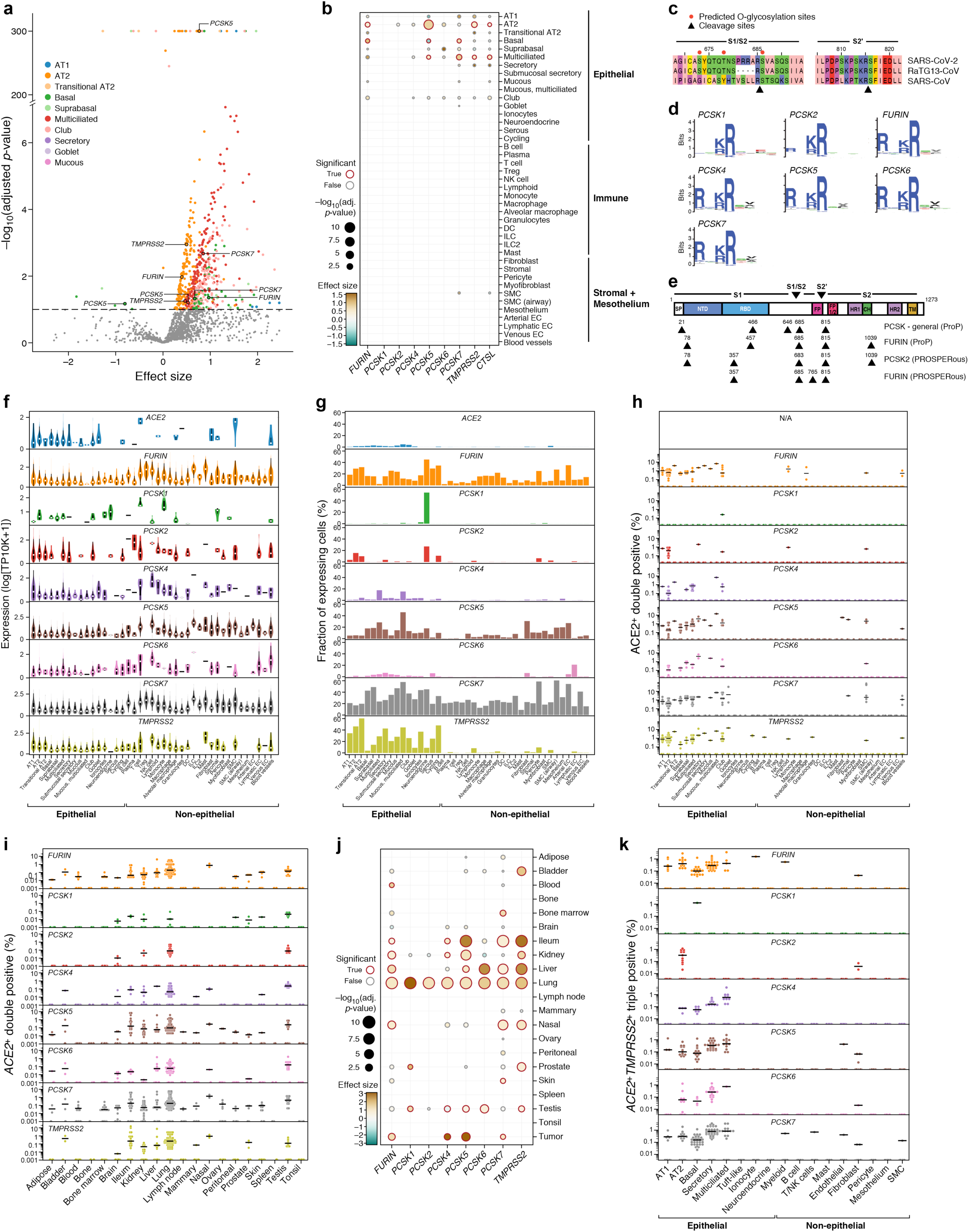
Additional analyses to identify other proteases that may have a role in infection. (a) Multiple proteases are co-expressed with *ACE2* in another human lung scRNA-seq (“aggregated lung”). Scatter plot of significance (y axis, -log10(adjusted p value)) and effect size (x axis) of co-expression of each protease gene (dot) with *ACE* within each indicated epithelial cell type (color). Dashed line: significance threshold. *TMPRSS2* and *PCSK*s that significantly co-expressed with *ACE2* are marked. (**b**) *ACE2-protease* co-expression with *PCSKs*, *TMPRSS2* and *CTSL* across lung cell types (“aggregated lung”). Significance (dot size, -log10(adjusted p value)) and effect size (color) for co-expression of ACE2 with selected proteases (columns) across cell types (rows). (c-d) Predicted cleavage sites in the SARS-CoV-2 S-protein S1/S2 region. (**c**) Multiple amino acid sequence alignment of SARS-CoV-2 S-protein S1/S2 region with orthologous sequences from other betacoronaviruses (top) and polybasic cleavage sites of other human pathogenic viruses (bottom). (**d**) Sequence logo plot showing cleavage site preference derived from MEROPS database for PCSK1, PCSK2, FURIN, PCSK4, PCSK5, PCSK6 and PCSK7. (**e**) Protease cleavage sites (triangles) predicted by ProP and PROSPERous in the SARS-CoV-2 spike protein. Top: Full-length SARS-CoV-2 S-protein sequence schematic with predicted functional protein domains and motifs. Numbers: amino acid residues after which cleavage occurs; SP: signal peptide; NTD: N-terminal domain; RBD: Receptor-binding domain; FP: Fusion peptide; FP1/2: Fusion peptide 1/2; HR1: Heptad repeat 1; CH: connecting helix; HR2: Heptad repeat 2; TM: Transmembrane domain. (**f,g**) Multiple proteases are expressed across lung cell types (“aggregated lung”). (f) Distribution of expression (y axis) for *ACE2*, *PCSKs* and *TMPRSS2* across lung cell types (x axis). White dot: median expression. (g) Proportion of cells (y axis) expressing *ACE2*, *PCSK* family or *TMPRSS2* across lung cell types (x axis), ordered by compartment. (**h**) *ACE2*^+^*PCSK*^+^ dual positive cells across lung cell types. Fraction (y axis) of different *ACE2*^+^*PCSK*^+^ or *ACE2*^+^*TMPRSS2*^+^ dual positive cells across lung cell types, ordered by compartment (x axis). Dots: different individuals, line: median. (**i,j**) *ACE2*^+^*PCSK*^+^ co-expression across human tissues (collection of published scRNA seq datasets). (i) Percent (y axis) of different *ACE2*^+^*PCSK*^+^ or *ACE2*^+^*TMPRSS2*^+^ dual positive cells across human tissues (x axis). Dots: different individuals, line: median. (j) *ACE2* co-expression with *PCSKs* or *TMPRSS2* across human tissues. Significance (dot size) and effect size (dot color) of co-expression. (**k**) Fraction of *ACE2*^+^*TMPRSS2^+^PCSK*^+^ cells across lung cell types (“Regev/Rajagopal dataset”). Dots: individuals, line: median.

**Extended Data Figure 15.**
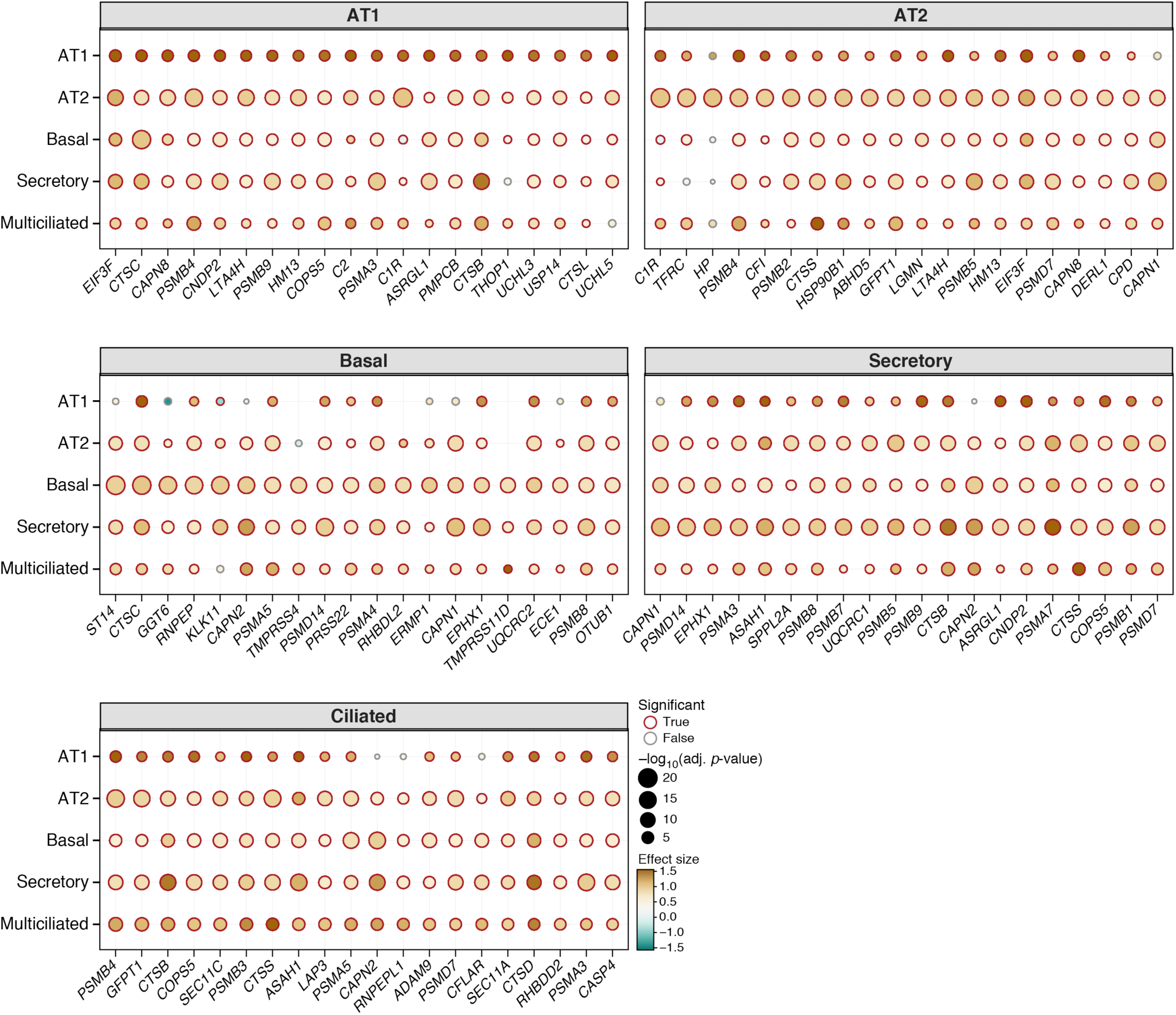
*ACE2-protease* co-expression of the top 20 most significantly co-expressed human proteases in key lung epithelial cell types. Significance (dot size) and effect size (dot color) of co-expression of each protease (columns) with ACE2 in each cell subset (rows).

**Extended Data Figure 16.**
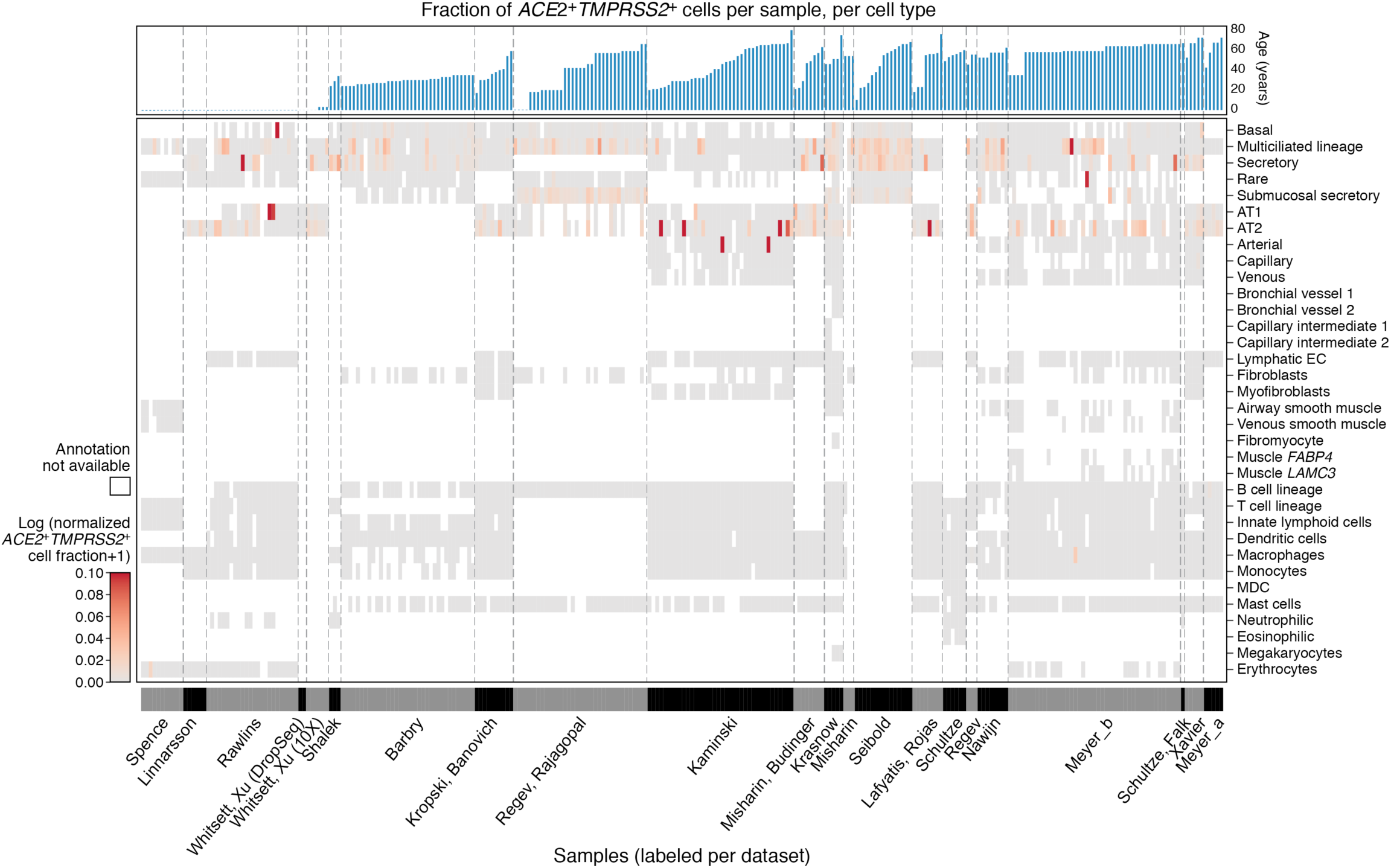
Variation in fraction of *ACE2*^+^*TMPRSS2*^+^ cells The normalized fraction of *ACE2*^+^*TMPRSS2*^+^ cells in 282 lung and nasal samples from 164 donors, subdivided by cell type. Normalization was performed by dividing the fraction by the median total counts per cell in the cell type of the sample. Samples are grouped by dataset and ordered by donor age within each dataset (blue bars at the top). Datasets are ordered by mean age of donors. White patches indicate the cell type annotation was not observed in the sample. This occurs either due to coarseness of annotation, or absence of cell type in the sample.

## REFERENCES

1. Xu, Z. et al. Pathological findings of COVID-19 associated with acute respiratory distress syndrome. Lancet Respir Med 8, 420–422 (2020).

2. Huang, C. et al. Clinical features of patients infected with 2019 novel coronavirus in Wuhan, China. Lancet 395, 497–506 (2020).

3. Fan, Z. et al. Clinical Features of COVID-19 Related Liver Damage. *medRxiv* (2020).

4. Guan, W.-J. et al. Clinical Characteristics of Coronavirus Disease 2019 in China. N. Engl. J. Med. (2020) doi:10.1056/NEJMoa2002032.

5. Zheng, Y.-Y., Ma, Y.-T., Zhang, J.-Y. & Xie, X. COVID-19 and the cardiovascular system. Nat. Rev. Cardiol. (2020) doi:10.1038/s41569-020-0360-5.

6. Ren, L.-L. et al. Identification of a novel coronavirus causing severe pneumonia in human: a descriptive study. Chin. Med. J. (2020) doi:10.1097/CM9.0000000000000722.

7. Wang, C., Horby, P. W., Hayden, F. G. & Gao, G. F. A novel coronavirus outbreak of global health concern. The Lancet vol. 395 470–473 (2020).

8. Rothan, H. A. & Byrareddy, S. N. The epidemiology and pathogenesis of coronavirus disease (COVID-19) outbreak. J. Autoimmun. 102433 (2020).

9. Cheng, Y. et al. Kidney impairment is associated with in-hospital death of COVID-19 patients. Nephrology (2020) doi:10.1101/2020.02.18.20023242.

10. Hui, H. et al. Clinical and radiographic features of cardiac injury in patients with 2019 novel coronavirus pneumonia. Cardiovascular Medicine (2020) doi:10.1101/2020.02.24.20027052.

11. Xu, Y. et al. Characteristics of pediatric SARS-CoV-2 infection and potential evidence for persistent fecal viral shedding. Nature Medicine (2020) doi:10.1038/s41591-020-0817-4.

12. CDC. Coronavirus Disease 2019 (COVID-19). Centers for Disease Control and Prevention https://www.cdc.gov/coronavirus/2019-ncov/hcp/clinical-guidance-management-patients.html (2020).

13. Wang, W. et al. Detection of SARS-CoV-2 in Different Types of Clinical Specimens. JAMA (2020) doi:10.1001/jama.2020.3786.

14. Shen, Q., Xiao, X., Aierken, A., Liao, M. & Hua, J. The ACE2 Expression in Sertoli cells and Germ cells may cause male reproductive disorder after SARS-CoV-2 Infection. (2020) doi:10.31219/osf.io/fs5hd.

15. Cai, X. An Insight of comparison between COVID-19 (2019-nCoV disease) and SARS in pathology and pathogenesis. Open Science Framework (2020) doi:10.31219/osf.io/hw34x.

16. Dong, L. et al. Possible Vertical Transmission of SARS-CoV-2 From an Infected Mother to Her Newborn. JAMA (2020) doi:10.1001/jama.2020.4621.

17. Chen, H. et al. Clinical characteristics and intrauterine vertical transmission potential of COVID-19 infection in nine pregnant women: a retrospective review of medical records. Lancet 395, 809–815 (2020).

18. Zeng, L. et al. Neonatal Early-Onset Infection With SARS-CoV-2 in 33 Neonates Born to Mothers With COVID-19 in Wuhan, China. JAMA Pediatr. (2020) doi:10.1001/jamapediatrics.2020.0878.

19. Zeng, H. et al. Antibodies in Infants Born to Mothers With COVID-19 Pneumonia. JAMA (2020) doi:10.1001/jama.2020.4861.

20. Schwartz, D. A. An Analysis of 38 Pregnant Women with COVID-19, Their Newborn Infants, and Maternal-Fetal Transmission of SARS-CoV-2: Maternal Coronavirus Infections and Pregnancy Outcomes. Arch. Pathol. Lab. Med. (2020) doi:10.5858/arpa.2020-0901-SA.

21. Chen, Y. et al. Infants Born to Mothers With a New Coronavirus (COVID-19). Frontiers in Pediatrics 8, 104 (2020).

22. Yang Li et al. Lack of Vertical Transmission of Severe Acute Respiratory Syndrome Coronavirus 2, China. Emerging Infectious Disease journal 26, (2020).

23. Li, R. et al. Substantial undocumented infection facilitates the rapid dissemination of novel coronavirus (SARS-CoV2). Science (2020) doi:10.1126/science.abb3221.

24. Riou, J., Hauser, A., Counotte, M. J. & Althaus, C. L. Adjusted age-specific case fatality ratio during the COVID-19 epidemic in Hubei, China, January and February 2020. Epidemiology (2020) doi:10.1101/2020.03.04.20031104.

25. Wu, J. T. et al. Estimating clinical severity of COVID-19 from the transmission dynamics in Wuhan, China. Nat. Med. (2020) doi:10.1038/s41591-020-0822-7.

26. Nick Wilson, Amanda Kvalsvig, Lucy Telfar Barnard & Michael G. Baker. Case-Fatality Risk Estimates for COVID-19 Calculated by Using a Lag Time for Fatality. Emerging Infectious Disease journal 26, (2020).

27. Kenji Mizumoto & Gerardo Chowell. Estimating Risk for Death from 2019 Novel Coronavirus Disease, China, January–February 2020. Emerging Infectious Disease journal 26, (2020).

28. Baud, D. et al. Real estimates of mortality following COVID-19 infection. Lancet Infect. Dis. (2020) doi:10.1016/S1473-3099(20)30195-X.

29. Wang, C. et al. Evolving Epidemiology and Impact of Non-pharmaceutical Interventions on the Outbreak of Coronavirus Disease 2019 in Wuhan, China. Epidemiology (2020) doi:10.1101/2020.03.03.20030593.

30. del Rio, C. & Malani, P. N. COVID-19—New Insights on a Rapidly Changing Epidemic. JAMA (2020) doi:10.1001/jama.2020.3072.

31. Lu, X. et al. SARS-CoV-2 Infection in Children. N. Engl. J. Med. (2020) doi:10.1056/NEJMc2005073.

32. Vardavas, C. I. & Nikitara, K. COVID-19 and smoking: A systematic review of the evidence. Tob. Induc. Dis. 18, 20 (2020).

33. Inciardi, R. M. et al. Cardiac Involvement in a Patient With Coronavirus Disease 2019 (COVID-19). JAMA Cardiol (2020) doi:10.1001/jamacardio.2020.1096.

34. Bonow, R. O., Fonarow, G. C., O’Gara, P. T. & Yancy, C. W. Association of Coronavirus Disease 2019 (COVID-19) With Myocardial Injury and Mortality. JAMA Cardiol (2020) doi:10.1001/jamacardio.2020.1105.

35. Fehr, A. R. & Perlman, S. Coronaviruses: an overview of their replication and pathogenesis. Methods Mol. Biol. 1282, 1–23 (2015).

36. Wrapp, D. et al. Cryo-EM structure of the 2019-nCoV spike in the prefusion conformation. Science (2020) doi:10.1126/science.abb2507.

37. Hoffmann, M. et al. SARS-CoV-2 Cell Entry Depends on ACE2 and TMPRSS2 and Is Blocked by a Clinically Proven Protease Inhibitor. Cell 0, (2020).

38. Wan, Y., Shang, J., Graham, R., Baric, R. S. & Li, F. Receptor recognition by novel coronavirus from Wuhan: An analysis based on decade-long structural studies of SARS. J. Virol. (2020) doi:10.1128/JVI.00127-20.

39. Bader, M. Tissue renin-angiotensin-aldosterone systems: Targets for pharmacological therapy. Annu. Rev. Pharmacol. Toxicol. 50, 439–465 (2010).

40. Li, W. et al. Angiotensin-converting enzyme 2 is a functional receptor for the SARS coronavirus. Nature 426, 450–454 (2003).

41. Walls, A. C., et al. Structure, Function, and Antigenicity of the SARS-CoV-2 Spike Glycoprotein. Cell (2020) doi:10.1016/j.cell.2020.02.058.

42. Wang, K. et al. SARS-CoV-2 invades host cells via a novel route: CD147-spike protein. bioRxiv 2020.03.14.988345 (2020) doi:10.1101/2020.03.14.988345.

43. Millet, J. K. & Whittaker, G. R. Host cell proteases: Critical determinants of coronavirus tropism and pathogenesis. Virus Res. 202, 120–134 (2015).

44. Matsuyama, S. et al. Enhanced isolation of SARS-CoV-2 by TMPRSS2-expressing cells. Proc. Natl. Acad. Sci. U. S. A. (2020) doi:10.1073/pnas.2002589117.

45. Sungnak, W., Huang, N., Bécavin, C., Berg, M. & HCA Lung Biological Network. SARS-CoV-2 Entry Genes Are Most Highly Expressed in Nasal Goblet and Ciliated Cells within Human Airways. arXiv [q-bio.CB] (2020).

46. Ziegler, C., et al. SARS-CoV-2 Receptor ACE2 is an Interferon-Stimulated Gene in Human Airway Epithelial Cells and Is Enriched in Specific Cell Subsets Across Tissues. (2020).

47. Lukassen, S. et al. SARS-CoV-2 receptor ACE2 and TMPRSS2 are predominantly expressed in a transient secretory cell type in subsegmental bronchial branches. bioRxiv 2020.03.13.991455 (2020) doi:10.1101/2020.03.13.991455.

48. Qi, F., Qian, S., Zhang, S. & Zhang, Z. Single cell RNA sequencing of 13 human tissues identify cell types and receptors of human coronaviruses. Biochem. Biophys. Res. Commun. (2020) doi:10.1016/j.bbrc.2020.03.044.

49. Wang, J. et al. ACE2 expression by colonic epithelial cells is associated with viral infection, immunity and energy metabolism. medRxiv (2020).

50. Huang, I.-C. et al. SARS coronavirus, but not human coronavirus NL63, utilizes cathepsin L to infect ACE2-expressing cells. J. Biol. Chem. 281, 3198–3203 (2006).

51. Chai, X. et al. Specific ACE2 Expression in Cholangiocytes May Cause Liver Damage After 2019-nCoV Infection. bioRxiv 2020.02.03.931766 (2020) doi:10.1101/2020.02.03.931766.

52. Gallagher, P. E., Ferrario, C. M. & Tallant, E. A. Regulation of ACE2 in cardiac myocytes and fibroblasts. Am. J. Physiol. Heart Circ. Physiol. 295, H2373–9 (2008).

53. Smillie, C. S. et al. Intra- and Inter-cellular Rewiring of the Human Colon during Ulcerative Colitis. Cell 178, 714–730.e22 (2019).

54. Emery, B. et al. Myelin gene regulatory factor is a critical transcriptional regulator required for CNS myelination. Cell 138, 172–185 (2009).

55. Zhou, F. et al. Clinical course and risk factors for mortality of adult inpatients with COVID-19 in Wuhan, China: a retrospective cohort study. Lancet (2020) doi:10.1016/S0140-6736(20)30566-3.

56. Rabin, R. C. Lost Sense of Smell May Be Peculiar Clue to Coronavirus Infection. The New York Times (2020).

57. Xia, J., Tong, J., Liu, M., Shen, Y. & Guo, D. Evaluation of coronavirus in tears and conjunctival secretions of patients with SARS-CoV-2 infection. J. Med. Virol. (2020) doi:10.1002/jmv.25725.

58. Tucker, N. R., et al. Myocyte Specific Upregulation of ACE2 in Cardiovascular Disease: Implications for SARS-CoV-2 mediated myocarditis. *medRxiv* 2020.04.09.20059204 (2020).

59. Henry, B. M. COVID-19, ECMO, and lymphopenia: a word of caution. Lancet Respir Med (2020) doi:10.1016/S2213-2600(20)30119-3.

60. Tan, L. et al. Lymphopenia predicts disease severity of COVID-19: a descriptive and predictive study. Signal Transduct Target Ther 5, 33 (2020).

61. Chen, Y. et al. The Novel Severe Acute Respiratory Syndrome Coronavirus 2 (SARS-CoV-2) Directly Decimates Human Spleens and Lymph Nodes. Infectious Diseases (except HIV/AIDS) (2020) doi:10.1101/2020.03.27.20045427.

62. Cai, H. Sex difference and smoking predisposition in patients with COVID-19. Lancet Respir Med (2020) doi:10.1016/S2213-2600(20)30117-X.

63. Vieira Braga, F. A. et al. A cellular census of human lungs identifies novel cell states in health and in asthma. Nat. Med. 25, 1153–1163 (2019).

64. Reyfman, P. A. et al. Single-Cell Transcriptomic Analysis of Human Lung Provides Insights into the Pathobiology of Pulmonary Fibrosis. Am. J. Respir. Crit. Care Med. 199, 1517–1536 (2019).

65. Morse, C. et al. Proliferating SPP1/MERTK-expressing macrophages in idiopathic pulmonary fibrosis. Eur. Respir. J. 54, (2019).

66. Madissoon, E. et al. scRNA-seq assessment of the human lung, spleen, and esophagus tissue stability after cold preservation. Genome Biol. 21, 1 (2019).

67. Ordovas-Montanes, J. et al. Allergic inflammatory memory in human respiratory epithelial progenitor cells. Nature 560, 649–654 (2018).

68. Miller, A. J. et al. In Vitro and In Vivo Development of the Human Airway at Single-Cell Resolution. Dev. Cell (2020) doi:10.1016/j.devcel.2020.01.033.

69. Deprez, M. et al. A single-cell atlas of the human healthy airways. bioRxiv 2019.12.21.884759 (2019) doi:10.1101/2019.12.21.884759.

70. Adams, T. S. et al. Single Cell RNA-seq reveals ectopic and aberrant lung resident cell populations in Idiopathic Pulmonary Fibrosis. bioRxiv 759902 (2019) doi:10.1101/759902.

71. Travaglini, K. J. et al. A molecular cell atlas of the human lung from single cell RNA sequencing. bioRxiv 742320 (2020) doi:10.1101/742320.

72. Habermann, A. C. et al. Single-cell RNA-sequencing reveals profibrotic roles of distinct epithelial and mesenchymal lineages in pulmonary fibrosis. bioRxiv 753806 (2019) doi:10.1101/753806.

73. Goldfarbmuren, K. C. et al. Dissecting the cellular specificity of smoking effects and reconstructing lineages in the human airway epithelium. bioRxiv 612747 (2019) doi:10.1101/612747.

74. Zimmermann, P. & Curtis, N. Coronavirus Infections in Children Including COVID-19. The Pediatric Infectious Disease Journal 1 (2020) doi:10.1097/inf.0000000000002660.

75. Sos, B. C. et al. Characterization of chromatin accessibility with a transposome hypersensitive sites sequencing (THS-seq) assay. Genome Biol. 17, 20 (2016).

76. Vento-Tormo, R. et al. Single-cell reconstruction of the early maternal-fetal interface in humans. Nature 563, 347–353 (2018).

77. Suryawanshi, H. et al. A single-cell survey of the human first-trimester placenta and decidua. Sci Adv 4, eaau4788 (2018).

78. Tsang, J. C. H. et al. Integrative single-cell and cell-free plasma RNA transcriptomics elucidates placental cellular dynamics. Proc. Natl. Acad. Sci. U. S. A. 114, E7786–E7795 (2017).

79. Chan, C.-M. et al. Carcinoembryonic Antigen-Related Cell Adhesion Molecule 5 Is an Important Surface Attachment Factor That Facilitates Entry of Middle East Respiratory Syndrome Coronavirus. J. Virol. 90, 9114–9127 (2016).

80. Wahl, S. M. et al. Secretory leukocyte protease inhibitor (SLPI) in mucosal fluids inhibits HIV-I. Oral Dis. 3 **Suppl 1**, S64–9 (1997).

81. Turula, H. & Wobus, C. E. The Role of the Polymeric Immunoglobulin Receptor and Secretory Immunoglobulins during Mucosal Infection and Immunity. Viruses 10, (2018).

82. Burkhardt, A. M. et al. CXCL17 is a mucosal chemokine elevated in idiopathic pulmonary fibrosis that exhibits broad antimicrobial activity. J. Immunol. 188, 6399–6406 (2012).

83. Debbabi, H. et al. Primary type II alveolar epithelial cells present microbial antigens to antigen-specific CD4+ T cells. Am. J. Physiol. Lung Cell. Mol. Physiol. 289, L274–9 (2005).

84. Yue, Y. et al. SARS-Coronavirus Open Reading Frame-3a drives multimodal necrotic cell death. Cell Death Dis. 9, 904 (2018).

85. Burkard, C. et al. Coronavirus cell entry occurs through the endo-/lysosomal pathway in a proteolysis-dependent manner. PLoS Pathog. 10, e1004502 (2014).

86. Liao, Y.-C. et al. IL-19 induces production of IL-6 and TNF-alpha and results in cell apoptosis through TNF-alpha. J. Immunol. 169, 4288–4297 (2002).

87. Starner, T. D., Barker, C. K., Jia, H. P., Kang, Y. & McCray, P. B., Jr. CCL20 is an inducible product of human airway epithelia with innate immune properties. Am. J. Respir. Cell Mol. Biol. 29, 627–633 (2003).

88. Wishart, D. S. et al. DrugBank 5.0: a major update to the DrugBank database for 2018. Nucleic Acids Res. 46, D1074–D1082 (2018).

89. Gordon, D. E. et al. A SARS-CoV-2-Human Protein-Protein Interaction Map Reveals Drug Targets and Potential Drug-Repurposing. bioRxiv 2020.03.22.002386 (2020) doi:10.1101/2020.03.22.002386.

90. Luan, H. H. et al. GDF15 Is an Inflammation-Induced Central Mediator of Tissue Tolerance. Cell 178, 1231–1244.e11 (2019).

91. Dhar, P. & McAuley, J. The Role of the Cell Surface Mucin MUC1 as a Barrier to Infection and Regulator of Inflammation. Front. Cell. Infect. Microbiol. 9, 117 (2019).

92. Herold, T., III et al. Level of IL-6 predicts respiratory failure in hospitalized symptomatic COVID-19 patients. Infectious Diseases (except HIV/AIDS) (2020) doi:10.1101/2020.04.01.20047381.

93. Schep, A. N., Wu, B., Buenrostro, J. D. & Greenleaf, W. J. chromVAR: inferring transcription-factor-associated accessibility from single-cell epigenomic data. Nat. Methods 14, 975–978 (2017).

94. Pedersen, K. B., Chodavarapu, H. & Lazartigues, E. Forkhead Box Transcription Factors of the FOXA Class Are Required for Basal Transcription of Angiotensin-Converting Enzyme 2. J Endocr Soc 1, 370–384 (2017).

95. Raredon, M. S. B. et al. Single-cell connectomic analysis of adult mammalian lungs. Sci Adv 5, eaaw3851 (2019).

96. Efremova, M., Vento-Tormo, M., Teichmann, S. A. & Vento-Tormo, R. CellPhoneDB: inferring cell-cell communication from combined expression of multi-subunit ligand-receptor complexes. Nat. Protoc. (2020) doi:10.1038/s41596-020-0292-x.

97. Bruce, A. G., Hoggatt, I. H. & Rose, T. M. Oncostatin M is a differentiation factor for myeloid leukemia cells. J. Immunol. 149, 1271–1275 (1992).

98. Li, W. et al. Efficient replication of severe acute respiratory syndrome coronavirus in mouse cells is limited by murine angiotensin-converting enzyme 2. J. Virol. 78, 11429–11433 (2004).

99. Montoro, D. T. et al. A revised airway epithelial hierarchy includes CFTR-expressing ionocytes. Nature 560, 319–324 (2018).

100. The Mouse in Biomedical Research. (Elsevier Science, 2006).

101. Gebel, S. et al. The transcriptome of Nrf2-/-mice provides evidence for impaired cell cycle progression in the development of cigarette smoke-induced emphysematous changes. Toxicol. Sci. 115, 238–252 (2010).

102. Pérez-Silva, J. G., Español, Y., Velasco, G. & Quesada, V. The Degradome database: expanding roles of mammalian proteases in life and disease. Nucleic Acids Res. 44, D351–5 (2016).

103. Izaguirre, G. The Proteolytic Regulation of Virus Cell Entry by Furin and Other Proprotein Convertases. Viruses 11, (2019).

104. Braun, E. & Sauter, D. Furin-mediated protein processing in infectious diseases and cancer. Clin Transl Immunol 8, 9298 (2019).

105. Coutard, B. et al. The spike glycoprotein of the new coronavirus 2019-nCoV contains a furin-like cleavage site absent in CoV of the same clade. Antiviral Res. 176, 104742 (2020).

106. Jaimes, J. A., André, N. M., Millet, J. K. & Whittaker, G. R. Structural modeling of 2019-novel coronavirus (nCoV) spike protein reveals a proteolytically-sensitive activation loop as a distinguishing feature compared to SARS-CoV and related SARS-like coronaviruses. bioRxiv 2020.02.10.942185 (2020) doi:10.1101/2020.02.10.942185.

107. Seidah, N. G. & Prat, A. The biology and therapeutic targeting of the proprotein convertases. Nat. Rev. Drug Discov. 11, 367–383 (2012).

108. Duckert, P., Brunak, S. & Blom, N. Prediction of proprotein convertase cleavage sites. Protein Eng. Des. Sel. 17, 107–112 (2004).

109. Song, J. et al. PROSPERous: high-throughput prediction of substrate cleavage sites for 90 proteases with improved accuracy. Bioinformatics 34, 684–687 (2018).

110. Millet, J. K. & Whittaker, G. R. Physiological and molecular triggers for SARS-CoV membrane fusion and entry into host cells. Virology 517, 3–8 (2018).

111. Glowacka, I. et al. Evidence that TMPRSS2 activates the severe acute respiratory syndrome coronavirus spike protein for membrane fusion and reduces viral control by the humoral immune response. J. Virol. 85, 4122–4134 (2011).

112. Lukassen, S. et al. SARS-CoV-2 receptor ACE2 and TMPRSS2 are primarily expressed in bronchial transient secretory cells. EMBO J. (2020) doi:10.15252/embj.20105114.

113. Hamming, I. et al. Tissue distribution of ACE2 protein, the functional receptor for SARS coronavirus. A first step in understanding SARS pathogenesis. J. Pathol. 203, 631– 637 (2004).

114. Zhao, Y. et al. Single-cell RNA expression profiling of ACE2, the putative receptor of Wuhan 2019-nCov. bioRxiv 2020.01.26.919985 (2020) doi:10.1101/2020.01.26.919985.

115. Venkatakrishnan, A. J. et al. Knowledge synthesis from 100 million biomedical documents augments the deep expression profiling of coronavirus receptors. doi:10.1101/2020.03.24.005702.

116. Mao, L. et al. Neurological Manifestations of Hospitalized Patients with COVID-19 in Wuhan, China: A Retrospective Case Series Study. Lancet (2020) doi:10.2139/ssrn.3544840.

117. Poyiadji, N. et al. COVID-19–associated Acute Hemorrhagic Necrotizing Encephalopathy: CT and MRI Features. Radiology 201187 (2020) doi:10.1148/radiol.2020201187.

118. Helms, J. et al. Neurologic Features in Severe SARS-CoV-2 Infection. N. Engl. J. Med. (2020) doi:10.1056/NEJMc2008597.

119. Morishima, T. et al. Encephalitis and encephalopathy associated with an influenza epidemic in Japan. Clin. Infect. Dis. 35, 512–517 (2002).

120. Okabe, N., Yamashita, K., Taniguchi, K. & Inouye, S. Influenza surveillance system of Japan and acute encephalitis and encephalopathy in the influenza season. Pediatr. Int. 42, 187–191 (2000).

121. McGavern, D. B. & Kang, S. S. Illuminating viral infections in the nervous system. Nat. Rev. Immunol. 11, 318–329 (2011).

122. Desforges, M., Le Coupanec, A., Stodola, J. K., Meessen-Pinard, M. & Talbot, P. J. Human coronaviruses: viral and cellular factors involved in neuroinvasiveness and neuropathogenesis. Virus Res. 194, 145–158 (2014).

123. Li, Y., Bai, W. & Hashikawa, T. The neuroinvasive potential of SARS-CoV2 may be at least partially responsible for the respiratory failure of COVID-19 patients. Journal of Medical Virology (2020) doi:10.1002/jmv.25728.

124. Bohmwald, K., Gálvez, N. M. S., Ríos, M. & Kalergis, A. M. Neurologic Alterations Due to Respiratory Virus Infections. Front. Cell. Neurosci. 12, 386 (2018).

125. Xu, J. et al. Detection of severe acute respiratory syndrome coronavirus in the brain: potential role of the chemokine mig in pathogenesis. Clin. Infect. Dis. 41, 1089–1096 (2005).

126. Kawada, J.-I. et al. Systemic cytokine responses in patients with influenza-associated encephalopathy. J. Infect. Dis. 188, 690–698 (2003).

127. Netland, J., Meyerholz, D. K., Moore, S., Cassell, M. & Perlman, S. Severe acute respiratory syndrome coronavirus infection causes neuronal death in the absence of encephalitis in mice transgenic for human ACE2. J. Virol. 82, 7264–7275 (2008).

128. Brann, D. H., Tsukahara, T., Weinreb, C., Logan, D. W. & Datta, S. R. Non-neural expression of SARS-CoV-2 entry genes in the olfactory epithelium suggests mechanisms underlying anosmia in COVID-19 patients. bioRxiv (2020) doi:10.1101/2020.03.25.009084.

129. Forrester, J. V., McMenamin, P. G. & Dando, S. J. CNS infection and immune privilege. Nat. Rev. Neurosci. 19, 655–671 (2018).

130. Matsuda, K. et al. The vagus nerve is one route of transneural invasion for intranasally inoculated influenza a virus in mice. Vet. Pathol. 41, 101–107 (2004).

131. Gesser, R. M. & Koo, S. C. Oral inoculation with herpes simplex virus type 1 infects enteric neuron and mucosal nerve fibers within the gastrointestinal tract in mice. J. Virol. 70, 4097–4102 (1996).

132. Hamid, S. H. M. et al. Seizures and Encephalitis in Myelin Oligodendrocyte Glycoprotein IgG Disease vs Aquaporin 4 IgG Disease. JAMA Neurol. 75, 65–71 (2018).

133. Wang, L. et al. Encephalitis is an important clinical component of myelin oligodendrocyte glycoprotein antibody associated demyelination: a single-center cohort study in Shanghai, China. Eur. J. Neurol. 26, 168–174 (2019).

134. Kakalacheva, K. et al. Infectious Mononucleosis Triggers Generation of IgG Auto-Antibodies against Native Myelin Oligodendrocyte Glycoprotein. Viruses 8, (2016).

135. Heink, S. et al. Trans-presentation of IL-6 by dendritic cells is required for the priming of pathogenic T17 cells. Nat. Immunol. 18, 74–85 (2017).

136. Chastain, E. M. L. & Miller, S. D. Molecular mimicry as an inducing trigger for CNS autoimmune demyelinating disease. Immunol. Rev. 245, 227–238 (2012).

137. Wagner, C. A., Roqué, P. J., Mileur, T. R., Liggitt, D. & Goverman, J. M. Myelin-specific CD8 T cells exacerbate brain inflammation in CNS autoimmunity. Journal of Clinical Investigation vol. 130 203–213 (2019).

138. Dos Santos, T. et al. Zika Virus and the Guillain-Barré Syndrome - Case Series from Seven Countries. N. Engl. J. Med. 375, 1598–1601 (2016).

139. Yuki, N. & Hartung, H.-P. Guillain-Barré syndrome. N. Engl. J. Med. 366, 2294–2304 (2012).

140. Bernard, C. C. A., Johns, T. G., de Rosbo, N. K. & Ichikawa, M. A peptide from myelin oligodendrocyte glycoprotein that induces demyelinating encephalomyelitis resembling multiple sclerosis. Journal of Neuroimmunology vol. 54 153 (1994).

141. Berger, T. et al. Antimyelin antibodies as a predictor of clinically definite multiple sclerosis after a first demyelinating event. N. Engl. J. Med. 349, 139–145 (2003).

142. Egg, R., Reindl, M., Deisenhammer, F., Linington, C. & Berger, T. Anti-MOG and anti-MBP antibody subclasses in multiple sclerosis. Mult. Scler. 7, 285–289 (2001).

143. Duc, D. et al. Disrupting Myelin-Specific Th17 Cell Gut Homing Confers Protection in an Adoptive Transfer Experimental Autoimmune Encephalomyelitis. Cell Rep. 29, 378–390.e4 (2019).

144. Toscano, G. et al. Guillain-Barré Syndrome Associated with SARS-CoV-2. N. Engl. J. Med. (2020) doi:10.1056/NEJMc2009191.

145. Simon, O. et al. Early Guillain–Barré Syndrome associated with acute dengue fever. Journal of Clinical Virology vol. 77 29–31 (2016).

146. Dirlikov, E. et al. Clinical Features of Guillain-Barré Syndrome With vs Without Zika Virus Infection, Puerto Rico, 2016. JAMA Neurol. 75, 1089–1097 (2018).

147. Smith, J. C. & Sheltzer, J. M. Cigarette smoke triggers the expansion of a subpopulation of respiratory epithelial cells that express the SARS-CoV-2 receptor ACE2. bioRxiv 2020.03.28.013672 (2020) doi:10.1101/2020.03.28.013672.

148. Imai, Y. et al. Angiotensin-converting enzyme 2 protects from severe acute lung failure. Nature 436, 112–116 (2005).

149. Wevers, B. A. & van der Hoek, L. Renin-angiotensin system in human coronavirus pathogenesis. Future Virol. 5, 145–161 (2010).

150. Smoking in men vs. women. Our World in Data https://ourworldindata.org/grapher/comparing-the-share-of-men-and-women-who-are-smoking.

151. Liao, M. et al. The landscape of lung bronchoalveolar immune cells in COVID-19 revealed by single-cell RNA sequencing. medRxiv (2020).

152. Tata, A. et al. Myoepithelial Cells of Submucosal Glands Can Function as Reserve Stem Cells to Regenerate Airways after Injury. Cell Stem Cell 22, 668–683.e6 (2018).

153. Lynch, T. J. et al. Submucosal Gland Myoepithelial Cells Are Reserve Stem Cells That Can Regenerate Mouse Tracheal Epithelium. Cell Stem Cell 22, 653–667.e5 (2018).

154. Funk, C. J. et al. Infection of human alveolar macrophages by human coronavirus strain 229E. J. Gen. Virol. 93, 494–503 (2012).

155. Zhou, Y. et al. A single asparagine-linked glycosylation site of the severe acute respiratory syndrome coronavirus spike glycoprotein facilitates inhibition by mannose-binding lectin through multiple mechanisms. J. Virol. 84, 8753–8764 (2010).

156. Wang, R., Xiao, H., Guo, R., Li, Y. & Shen, B. The role of C5a in acute lung injury induced by highly pathogenic viral infections. Emerg. Microbes Infect. 4, e28 (2015).

157. Gao, T. et al. Highly pathogenic coronavirus N protein aggravates lung injury by MASP-2-mediated complement over-activation. *medRxiv* (2020).

158. Cheng, Z., Brown, L. E. & Wathes, D. C. Bovine Viral Diarrhoea Virus Infection Disrupts Uterine Interferon Stimulated Gene Regulatory Pathways During Pregnancy Recognition in Cows. Viruses 12, (2019).

159. Coupanec, A. L. et al. Cleavage of a Neuroinvasive Human Respiratory Virus Spike Glycoprotein by Proprotein Convertases Modulates Neurovirulence and Virus Spread within the Central Nervous System. PLOS Pathogens vol. 11 e1005261 (2015).

160. Braun, E. et al. Guanylate-Binding Proteins 2 and 5 Exert Broad Antiviral Activity by Inhibiting Furin-Mediated Processing of Viral Envelope Proteins. Cell Rep. 27, 2092–2104.e10 (2019).

161. Barrett, T., et al. NCBI GEO: archive for functional genomics data sets—update. Nucleic Acids Research vol. 41 D991–D995 (2012).

162. Zheng, G. X. Y. et al. Massively parallel digital transcriptional profiling of single cells. Nat. Commun. 8, 236 (2017).

163. Li, B. et al. Cumulus: a cloud-based data analysis framework for large-scale single-cell and single-nucleus RNA-seq. bioRxiv 823682 (2019) doi:10.1101/823682.

164. Korsunsky, I. et al. Fast, sensitive and accurate integration of single-cell data with Harmony. Nature Methods vol. 16 1289–1296 (2019).

165. Wolf, F. A., Angerer, P. & Theis, F. J. SCANPY: large-scale single-cell gene expression data analysis. Genome Biol. 19, 15 (2018).

166. Seabold, S. & Perktold, J. Statsmodels: Econometric and Statistical Modeling with Python. Proceedings of the 9th Python in Science Conference (2010) doi:10.25080/majora-92bf1922-011.

167. Bates, D., Mächler, M., Bolker, B. & Walker, S. Fitting Linear Mixed-Effects Models Using lme4. *Journal of Statistical Software*, Articles 67, 1–48 (2015).

168. Stuart, T. et al. Comprehensive Integration of Single-Cell Data. Cell 177, 1888– 1902.e21 (2019).

169. Cusanovich, D. A. et al. Multiplex single cell profiling of chromatin accessibility by combinatorial cellular indexing. Science 348, 910–914 (2015).

170. McInnes, L. & Healy, J. UMAP: Uniform Manifold Approximation and Projection for Dimension Reduction. arXiv [stat.ML] (2018).

171. Fornes, O., et al. JASPAR 2020: update of the open-access database of transcription factor binding profiles. Nucleic Acids Res. 48, D87–D92 (2020).

172. Davis, C. A. et al. The Encyclopedia of DNA elements (ENCODE): data portal update. Nucleic Acids Res. 46, D794–D801 (2018).

173. Nagendran, M., Riordan, D. P., Harbury, P. B. & Desai, T. J. Automated cell-type classification in intact tissues by single-cell molecular profiling. eLife vol. 7 (2018).

174. Habermann, A. C. et al. Single-cell RNA-sequencing reveals profibrotic roles of distinct epithelial and mesenchymal lineages in pulmonary fibrosis. Genomics 1888 (2019).

175. Pedregosa, F. et al. Scikit-learn: Machine Learning in Python. J. Mach. Learn. Res. 12, 2825–2830 (2011).

176. Breiman, L., Friedman, J., Stone, C. J. & Olshen, R. A. Classification and regression trees. (CRC press, 1984).

177. Hagberg, A., Swart, P. & S Chult, D. Exploring network structure, dynamics, and function using NetworkX. http://conference.scipy.org/proceedings/SciPy2008/paper_2/full_text.pdf (2008).

178. Jacomy, M., Venturini, T., Heymann, S. & Bastian, M. ForceAtlas2, a continuous graph layout algorithm for handy network visualization designed for the Gephi software. PLoS One 9, e98679 (2014).

179. Raudvere, U., et al. g:Profiler: a web server for functional enrichment analysis and conversions of gene lists (2019 update). Nucleic Acids Research vol. 47 W191–W198 (2019).

180. Finak, G. et al. MAST: a flexible statistical framework for assessing transcriptional changes and characterizing heterogeneity in single-cell RNA sequencing data. Genome Biol. 16, 278 (2015).

181. Kuleshov, M. V. et al. Enrichr: a comprehensive gene set enrichment analysis web server 2016 update. Nucleic Acids Res. 44, W90–7 (2016).

